# *Salmonella* MgtC promotes polymyxin B resistance and systemic infection by activating LpxO-mediated lipid A hydroxylation

**DOI:** 10.64898/2026.11.23.768523

**Authors:** Nakjun Choi, Seungwoo Baek, Eunna Choi, Eun-Jin Lee

**Affiliations:** Department of Life Sciences, School of Life Sciences and Biotechnology, Korea University, Seoul, South Korea

**Author notes:** To Whom correspondence should be addressed: Eun-Jin Lee.

**Keywords:** *Salmonella*, MgtC, LpxO, Lipid A hydroxylation, Polymyxin B, Virulence

## Abstract

The intracellular pathogen *Salmonella* Typhimurium modifies the lipid A moiety of lipopolysaccharide through various lipid A-modifying enzymes, thereby acquiring resistance to host cationic antimicrobial peptides during infection. While regulation of these enzymes at the transcription level has been extensively studied, however, regulation at the activity level remains poorly understood. Here, we show that the *Salmonella* virulence factor MgtC interacts with LpxO, an enzyme that hydroxylates the 3’-myristoyl chain of hexa-acylated lipid A, identified through bacterial two-hybrid screening for potential MgtC interaction partners. MgtC binding to LpxO promotes lipid A hydroxylation in a concentration-dependent manner, and the Leu287 residue in the C-terminal helix of LpxO is required for this interaction. Substitution of Leu287 with Ala reduces MgtC binding without affecting intrinsic hydroxylase activity, consequently impairing MgtC-enhanced lipid A hydroxylation. This substitution decreases resistance to both the antimicrobial peptide polymyxin B and human serum, and attenuates virulence in a mouse infection model. These findings suggest that *Salmonella* exploits MgtC to promote LpxO hydroxylation activity, thereby enhancing survival inside the host.

**Significance statement:** How bacterial pathogens regulate lipid-modifying enzymes at the activity level during infection remains poorly understood. Here we show that the *Salmonella* virulence factor MgtC directly binds the lipid A hydroxylase LpxO through a transmembrane helix-helix interaction, activating LpxO-mediated lipid A hydroxylation in a concentration-dependent manner. This regulatory interaction occurs inside macrophages, where low magnesium and limited oxygen constrain LpxO activity, and is required for full resistance to cationic antimicrobial peptides, human serum complement, and systemic infection in mice. Because MgtC and LpxO co-occur across diverse Gram-negative pathogens, this work uncovers a broadly conserved post-translational mechanism by which intracellular bacteria remodel their outer membrane to evade innate immune defenses and promote virulence.

## Introduction

Gram-negative bacteria possess a unique outer membrane that provides an effective barrier against environmental stressors, including host-derived immune effectors. A key component of this outer membrane is lipopolysaccharide (LPS), which is located at the outer leaflet of the outer membrane, and whose lipid A moiety directly interacts with host immune molecules such as cationic antimicrobial peptides (CAMPs) and complement in human serum (1, 2). Lipid A consists of a hexa-acylated diphosphosaccharide backbone, which is further extended by core oligosaccharides and repeating O-antigen units (1). Many pathogens modify lipid A using enzymes that alter its charge and hydrophobicity, thereby reducing the binding affinity of host defense molecules to evade immune recognition and enhance survival inside the host (3, 4).

In the intracellular pathogen *Salmonella* Typhimurium (5), extensive lipid A modification also occurs after phagocytosis by macrophages (6). This modification includes addition of 4-amino-4-deoxy-L-arbinose (L-Ara4N) by ArnT (PbgP), addition of phosphoethanolamine (pEtN) by EptA (PmrC), palmitoylation by PagP, and hydroxylation of lipid A by LpxO, in response to low Mg^2+^, CAMPs, low pH, and high Fe^2+^ signals, all of which are physiologically relevant inside macrophages (Fig. S1) (7). The PmrA/PmrB two-component systems control expression of the *pbgP* and *pmrC* genes in response to high Fe^2+^, while the PhoP/PhoQ two-component system controls expression of *pagP* and *lpxO* in response to low Mg^2+^. Notably, although *lpxO* transcription is induced by PhoP/PhoQ in low Mg^2+^ conditions, previous research showed that lipid A hydroxylation is detected in both PhoP-repressing (high Mg^2+^) and PhoP-inducing (low Mg^2+^) conditions, and even in a *phoP* deletion mutant, suggesting that a basal level of *lpxO* expression exists independently of PhoP, low Mg^2+^ signal, or the intramacrophage environment (6). Adding further complexity, in low Mg^2+^, the PhoP/PhoQ two-component system controls production of the PmrD connector protein, which in turn binds PmrA response regulator and activates PmrA/PmrB-dependent signaling in this condition (8). These lipid A modifications allow *Salmonella* to resist antimicrobial peptides and serum-mediated killing (9, 10). Interestingly, among lipid A-modifying enzymes, LpxO is absent in *Escherichia coli*, although many gram-negative pathogens possess one or two *lpxO* genes (11–13). LpxO is a Fe^2+^/α-ketoglutarate-dependent dioxygenase that generates the 2-hydroxymyristoyl group on the 3’-myristoyl chain of lipid A (11), thereby contributing to resistance to polymyxin B. Lipid A hydroxylation increases the structural stability of lipid A, reducing the effectiveness of CAMPs by altering the charge and hydrophobicity of the bacterial outer membrane (14). However, unlike that of other lipid A modifying enzymes such as ArnT or EptA, the biological relevance of LpxO in the context of host-pathogen interaction has not been fully investigated (15, 16).

In addition to lipid A modifying genes, *pbgP* and *lpxO*, the PhoP/PhoQ two-component system controls many genes required for survival in intracellular conditions such as the phagosome, where magnesium limitation and antimicrobial peptides are prevalent (17). Among the genes regulated by PhoP/PhoQ, *mgtC* is located within the *mgtCBRU* operon and is required for survival within macrophages and for mouse virulence (18, 19). Over the years, MgtC has been found to interact with several proteins that contribute to *Salmonella* survival inside macrophages and to mouse virulence. Initially, the membrane F_o_F_1_ ATP synthase was identified as an interacting target, an interaction involved in maintaining bacterial cytoplasmic pH inside acidified macrophage phagosomes (19). PhoR, the histidine kinase of the phosphate-sensing PhoB/PhoR two-component system, was also identified as a binding target that allows *Salmonella* to promote phosphate uptake inside macrophages (20). Additionally, MgtC participates in interactions that lead to regulated proteolysis. MgtR binds MgtC and promotes its own MgtC degradation via the membrane-bound FtsH-mediated proteolysis (21), whereas MgtC interacts with the cytoplasmic PhoP protein of the PhoP/PhoQ two-component system and thereby protects PhoP from proteolysis by the ClpAP protease, constituting a positive feedback loop that reinforces the PhoP/PhoQ-controlled gene expression (22). Although several MgtC-interacting proteins have already been identified, there remains room for additional interactors, given that MgtC is highly induced inside macrophages or in Mg^2+^-limiting conditions (23, 24), and that other candidate interacting proteins have already been detected in previous immunoprecipitation-mass spectrometry experiments (19). In this study, we address this further by searching for MgtC-interacting proteins using a bacterial adenylate cyclase two-hybrid (BACTH) screen.

In this work, we discovered the lipid A hydroxylase LpxO as a partner protein interacting with MgtC through a bacterial adenylate cyclase two-hybrid (BACTH) screen. MgtC interacts with the C-terminal transmembrane helix of LpxO via its Leu287 residue. MgtC binding to LpxO enhanced lipid A hydroxylation in a concentration-dependent manner, whereas disruption of this interaction by the Leu287Ala substitution abolished the enhancement. Consequently, the MgtC-LpxO interaction significantly increased resistance to polymyxin B and to human serum-mediated killing, and also promoted mouse virulence. Our findings reveal, for the first time, that the MgtC virulence factor modulates lipid A modification by direct binding to a lipid A modifying enzyme. This interaction suggests a coordinated regulatory mechanism that may influence *Salmonella*’s overall adaptability and pathogenicity across diverse host environments, providing new insights into bacterial survival strategies within the host.

## Results

### Identification of LpxO as an MgtC-interacting partner by BACTH assay

To search for the interacting partners of MgtC, we utilized the BACTH screening (25). We used the pUT18 plasmid containing MgtC as bait (20) and constructed a Sau3AI-cut genomic DNA library in pKT25 or pKNT25. These plasmids were then co-transformed into BTH101 strain, and red colonies were isolated on MacConkey agar plates (Fig. 1A). Approximately 18,000 transformants were screened across three independent trials. Given the average insert size (500–2000 bp) and the size of the *S*. Typhimurium 14028s genome (4,870,265 bp), this corresponds to an estimated 4.62-fold physical coverage of the genome, or a ∼99% probability that any given genomic locus was represented at least once in the library, based on a Poisson distribution model of random fragmentation. From the initial screening, we obtained 69 positive clones, and after re-screening and sequencing, we narrowed these down to 16 positive clones (Fig. 1B, Table S1). Most of the identified clones correspond to proteins located in the inner membrane or associated with the membrane, suggesting that interaction with MgtC mostly occurs at the inner membrane. Among these clones, we further characterized the fragment containing the *lpxO* gene, because it was independently isolated four times from two different DNA libraries, confirming a reproducible interaction (Fig. 1C, L1 and L2). Sequencing analysis revealed that the partial *lpxO* gene (sequence region 4,541,552-4,540,869, NCBI accession: CP001363) was inserted into pKT25, with the LpxO^251-302^ portion expected to be translationally fused in-frame to the C-terminus of the T25 domain (Fig. 1C).

**Figure 1.**
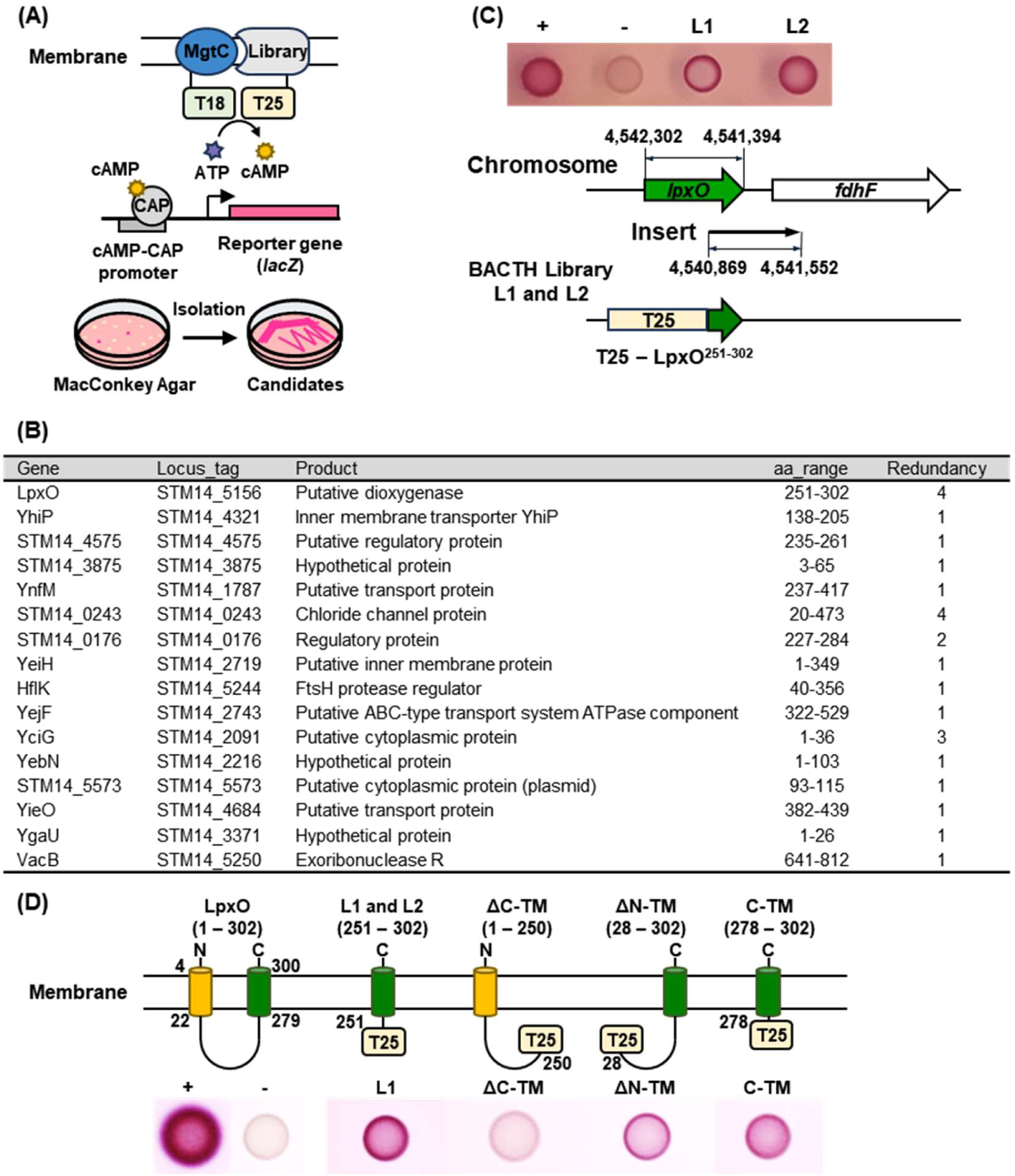
BACTH screen identifies LpxO as an MgtC-interacting protein. (A) Schematic representation of the BACTH screen. (B) List of MgtC-interacting clones identified from the BACTH screen. See also Table S1. (C) The L1 and L2 clones, which were identified through the BACTH screen to interact with MgtC, contain DNA fragments corresponding to LpxO^251-302^ fused to T25. (D) Bacterial two-hybrid assay of BTH101 cells coexpressing MgtC-T18 and C-terminally T25-fused LpxO domains: LpxO^1-250^ (ΔC-TM), LpxO^28-302^ (ΔN-TM), or LpxO^278-302^ (C-TM). Cells were spotted on MacConkey agar plates containing 1% maltose and 0.5 mM IPTG and incubated at 30°C for 48 hours. Red color indicates a positive interaction. +, positive control (MgtC-MgtR) (20, 21); -, negative control (MgtC-empty vector).

LpxO is a lipid A hydroxylase anchored in the inner membrane via two transmembrane helices (Fig. 1D). To confirm the MgtC-LpxO interaction, we cloned DNA fragments corresponding to C-terminal helix-deleted LpxO (ΔC-TM, LpxO^1-250^) and N-terminal helix-deleted LpxO (ΔN-TM, LpxO^28-302^) into pKT25, introduced them into BTH101, and spotted the cultures on MacConkey agar plates together with the previously isolated L1 clone. The reintroduced L1 clone still exhibited a strong red color on the plate, indicating a positive interaction (Fig. 1D). Removal of the C-terminal helix (ΔC-TM) abolished the red color, suggesting that this region is indeed required for the interaction (Fig. 1D). In contrast, removal of the N-terminal helix (ΔN-TM) did not affect the interaction (Fig. 1D). To further test whether the C-terminal helix alone is sufficient for the interaction, we removed most of the cytoplasmic portion and cloned only the C-terminal helix of LpxO into pKT25 (C-TM, LpxO^278-302^). This construct still maintained the interaction, indicating that LpxO interacts with MgtC via the C-terminal helix (Fig. 1D).

### The LpxO Leu287 is required for the MgtC interaction

LpxO is a lipid A hydroxylase containing two transmembrane helices and a long cytoplasmic domain (Fig. 2A). Since the LpxO C-terminal helix was found to be required for the MgtC interaction, we further narrowed down the interaction by replacing each amino acid in the C-terminal helix of LpxO (278-302, Fig. 2B) with alanine and tested which residue was critical for the MgtC interaction. Alanine scanning revealed that the Leu287-to-alanine substitution did not produce a blue color on LB X-gal plates (Fig. 2C), indicating that the Leu287 residue within the LpxO C-terminal helix is important for the MgtC interaction. To further explore this interaction, we generated strains carrying a C-terminal Myc tag on either the wild-type or the Leu287-to-Ala-substituted *lpxO* gene at its native chromosomal location (LpxO-M and L287A-M, respectively) and introduced a plasmid carrying a C-terminally FLAG-tagged *mgtC* gene driven by an IPTG-inducible promoter (pUH-*mgtC*-FLAG). Similarly to what we observed in the bacterial two hybrid assay, wild-type LpxO-Myc successfully immunoprecipitated MgtC-FLAG (Fig. 2D). However, the Leu287-to-Ala substitution significantly decreased the levels of co-precipitated MgtC protein (Fig. 2D), supporting the conclusion that LpxO Leu287 is a key residue for the MgtC interaction. Reciprocally, MgtC-FLAG also successfully immunoprecipitated chromosomally expressed wild-type LpxO-Myc, whereas MgtC-FLAG exhibited a clearly weakened interaction with the Leu287-to-Ala-substituted version of LpxO (Fig. 2E). Collectively, these data demonstrate that MgtC interacts with the lipid A hydroxylase LpxO via its C-terminal helix, within which Leu287 residue is a key residue for the interaction.

**Figure 2.**
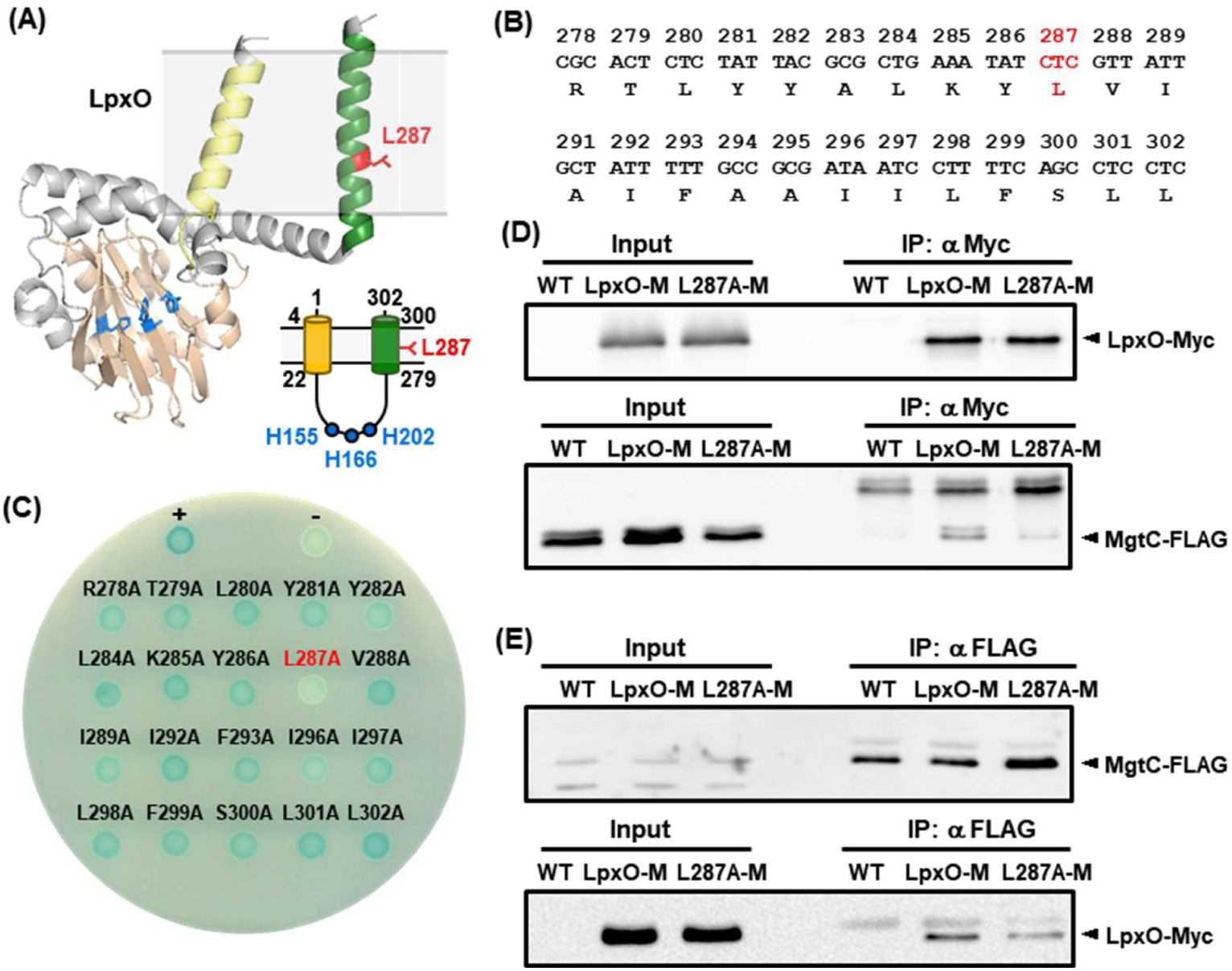
LpxO Leu287 is required for the MgtC interaction. (A) AlphaFold-assisted prediction of LpxO (AlphaFoldDB; A0A0F6BAE3) and its schematic diagram. The N-terminal helix is colored yellow (N-TM), the C-terminal helix green (C-TM), the conserved histidine residues blue (H155, H166, H202; see Fig. S2), and the Leu287 residue red. See also Fig. S2. (B) Amino acid sequence of LpxO C-terminal helix (278–302). (C) Bacterial two-hybrid assay of BTH101 cells coexpressing MgtC-T18 and C-terminally T25-fused LpxO C-terminal helix variants (278–302), with each residue individually substituted with alanine. Cells were spotted on LB agar plates containing 80 µM X-gal and 0.5 mM IPTG and incubated at 30°C for 48 hours. Blue color indicates a positive interaction. (D-E) Co-immunoprecipitation assays of C-terminally FLAG-tagged MgtC and C-terminally 8×Myc-tagged LpxO^WT^ or LpxO^L287A^, with an untagged strain as a control. FLAG-tagged MgtC was expressed from an IPTG-inducible pUHE21 plasmid, while 8×Myc-tagged LpxO (LpxO-M) and LpxO^L287A^ (L287A-M) were expressed from their native chromosomal locus. (D) Lysates were subjected to immunoprecipitation with an anti-Myc antibody, and western blot analysis using an anti-FLAG antibody detected co-precipitated MgtC. (E) Similarly, lysates were immunoprecipitated with an anti-FLAG antibody, followed by western blotting with an anti-Myc antibody to detect co-precipitated LpxO. In both cases, the MgtC-LpxO interaction was weakened in the LpxO L287A-M variant.

### The Ile69 and Ile73 residues in the 3^rd^ transmembrane helix of MgtC are required for the LpxO interaction

Next, we investigated which domain of MgtC is required for this interaction. Because the LpxO Leu287 residue is located in the middle of the C-terminal helix, we reasoned that LpxO binds to MgtC via a helix-helix interaction. MgtC is composed of five N-terminal transmembrane helices and a C-terminal cytoplasmic domain (Fig. S1B) (26). Thus, we cloned each of the five transmembrane helices into either pUT18 or pUT18C, depending on their topology, and introduced them into BTH101 cells harboring pKT25-*lpxO-*C-terminal helix, the minimal portion required for the MgtC interaction. When spotted on MacConkey agar plates, cells co-expressing an MgtC region containing transmembrane helix 3 (TM3, residues 66–82) and the LpxO C-terminal helix (C-TM) exhibited a strong red color, indicating a positive interaction (Fig. S1B).

We then replaced, one at a time, the TM3 residues expected to interact with LpxO (Q68, I69, V70, S71, I73, F75, and L76) with alanine and cloned the resulting mutant constructs into pUT18 (Fig. S1C). Alanine scanning of the residues in MgtC TM3 revealed that the Ile69-to-Ala or Ile73-to-Ala substitutions abolished the interaction with the LpxO C-TM (Fig. S1D), demonstrating that both the Ile69 and Ile73 residues in MgtC TM3 are required for the LpxO interaction. An AlphaFold-predicted model also showed that LpxO Leu287 is located in close proximity to Ile69 and Ile73 in MgtC, further supporting this interaction (Fig. S1E).

### Most pathogenic Gram-negative bacteria possess both MgtC and LpxO

To investigate whether MgtC and LpxO coexist in other species, we analyzed data from UniProtKB (www.uniprot.org). Excluding partial proteins, we identified 13,594 strains containing MgtC and 1,255 strains containing LpxO, and we then examined which genera possessed both proteins. Notably, most representative pathogenic Gram-negative bacteria were found to possess both MgtC and LpxO, except *Brucella* spp. and *Aeromonas* spp. (26–28) (Table 1, Supplementary Data). In contrast, the non-pathogenic bacterium *Escherichia coli* possessed neither protein, serving as a control. This finding suggests that co-occurrence of MgtC and LpxO may be a conserved feature among pathogenic Gram-negative bacteria that harbor MgtC.

**Table 1.** Distribution of MgtC and LpxO in representative gram-negative pathogenic bacteria. MgtC and LpxO protein sequences were obtained from the Uniprot database (www.uniprot.org). Representative Gram-negative pathogenic bacteria analyzed for MgtC included *Salmonella* spp. (n = 461), *Pseudomonas* spp. (n = 574), *Klebsiella* spp. (n = 58), *Burkholderia* spp. (n = 442), *Yersinia* spp. (n = 65), *Brucella* spp. (n = 43), *Acinetobacter* spp. (n = 173), *Enterobacter* spp. (n = 31), and *Aeromonas* spp. (n = 45). For LpxO, the dataset included *Salmonella* spp. (n = 336), *Pseudomonas* spp. (n = 329), *Klebsiella* spp. (n = 21), *Burkholderia* spp. (n = 19), *Yersinia* spp. (n = 12), *Acinetobacter* spp. (n = 120), and *Enterobacter* spp. (n = 13).

| Bacterial species | MgtC |  |  | LpxO |  |
|---|---|---|---|---|---|
|  | MgtC | 69 | 73 | LpxO | 287 |
| *Escherichia coli* | - | - | - | - | - |
| *Salmonella* spp. | + | Ile | Ile | + | Leu |
| *Pseudomonas* spp. | + | Val | Ile | + | Ala, Leu, Gly |
| *Klebsiella* spp. | + | Ile, Val | Ile | + | Leu |
| *Burkholderia* spp. | + | Val | Val | + | Ala, Leu, Val |
| *Yersinia* spp. | + | Ile, Val | Ile | + | Leu, Ile |
| *Acinetobacter* spp. | △ | Val | Ile | + | Phe, Leu |
| *Enterobacter* spp. | △ | Ile | Ile | + | Leu |
| *Brucella* spp. | + | Ile, Val | Ile, Val | - | - |
| *Aeromonas* spp. | △ | Ile | Ile | - | - |

+; presence, -; absence, △; MgtC homolog

We then further examined the residues implicated in the MgtC-LpxO interaction across these species. The amino acids at position 69 and 73 of MgtC are predominantly isoleucine or valine. Similarly, the amino acid at position 287 of LpxO is predominantly leucine, with occasional substitutions to valine, isoleucine, alanine, glycine, or phenylalanine (Table 1). Because most of these residues are branched-chain amino acids, this conservation suggests that the MgtC-LpxO interaction is mediated by hydrophobic interactions between these residues.

### Leu287 of LpxO is required for interaction with MgtC without affecting intrinsic hydroxylase activity

LpxO lipid A hydroxylase activity requires coordination of Fe^2+^ through key histidine residues, including H155, H166, and H202 (Fig. 2A and S2, blue), which are essential for its enzymatic function (28). These active sites are located in the long cytoplasmic loop flanked by two transmembrane helices. Notably, Leu287 is located in the C-terminal helix but outside this catalytic region, suggesting that it may mediate protein-protein interaction rather than catalysis. To test whether the Leu287 to Ala substitution affects the enzymatic function of LpxO, we cultured wild-type, Δ*lpxO*, and *lpxO*^L287A^ strains in N-minimal medium (NMM) containing 10 mM Mg^2+^, extracted lipid A, and analyzed the samples by MALDI-TOF MS.

*Salmonella* synthesizes di-phosphorylated hexa-acyl lipid A (1797 m/z) in its membrane. LpxO adds an oxygen atom to the 2’-myristate group of lipid A, increasing the molecular weight of the hydroxylated species by 16 Da to 1813 m/z (Fig. 3A). In the lipid A profile of wild-type strain grown in NMM containing 10 mM Mg^2+^, we detected distinct peaks corresponding to both unmodified (1797 m/z) and hydroxylated (1813 m/z) lipid A (Fig. 3B). In the Δ*lpxO* mutant, the peak corresponding to hydroxylated lipid A (1813 m/z) was absent, confirming the role of LpxO in lipid A hydroxylation. In contrast, the Leu287 to Ala substitution did not affect the hydroxylated lipid A peak, indicating that Leu287 is required for MgtC interaction but not for hydroxylase activity (Fig. 3B). However, instead of substituting Leu287 to Ala that maintains the C-terminal helix, removing the entire C-terminal helix of LpxO eliminated lipid A hydroxylase activity (Fig. S3). Likewise, removing the N-terminal helix also abolished lipid A hydroxylase activity (Fig. S3), indicating that localization of LpxO via the two transmembrane helices is required for lipid A hydroxylase activity.

**Figure 3.**
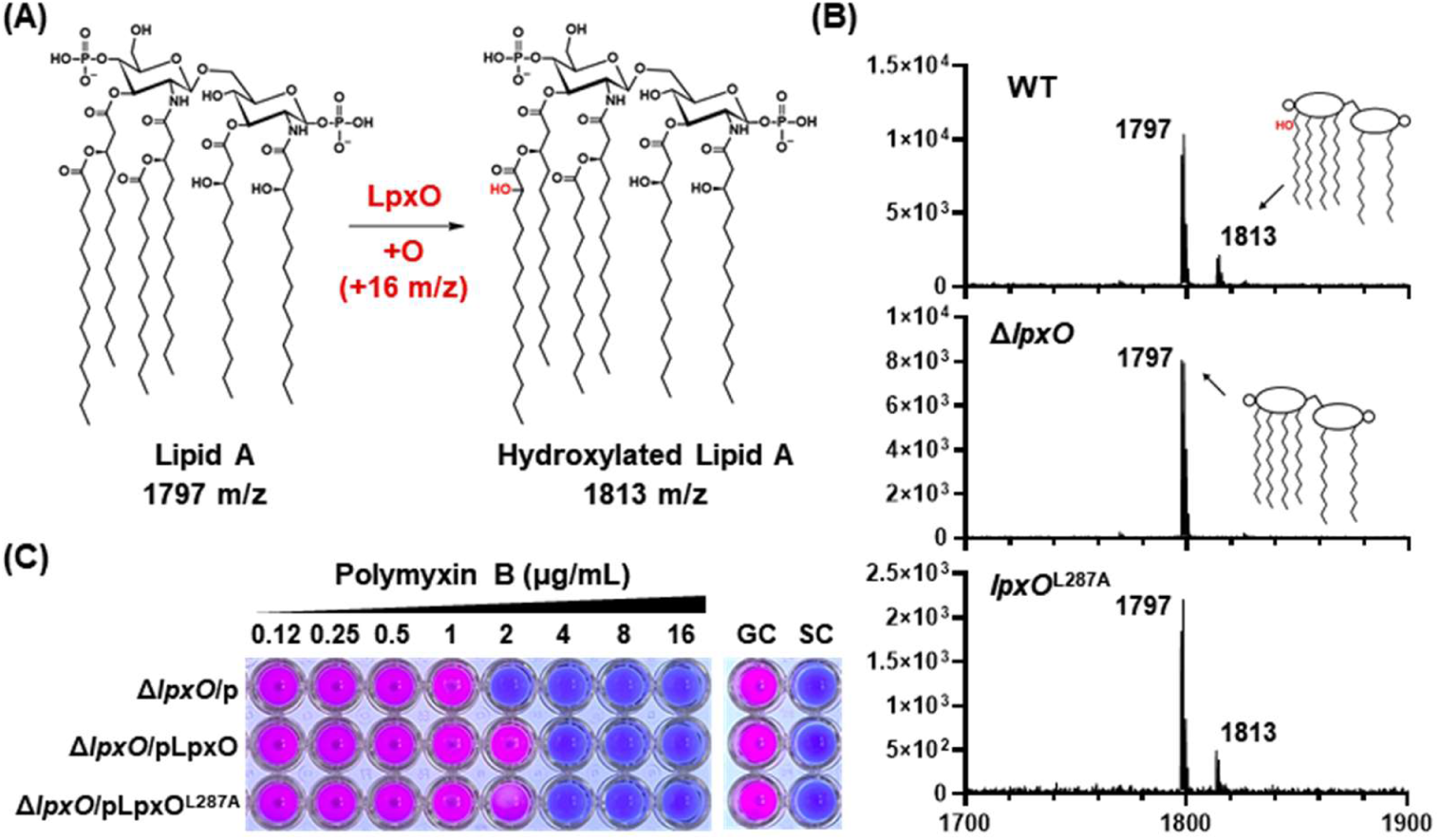
The Leu287-to-Ala substitution in LpxO maintains intrinsic lipid A hydroxylase activity. (A) Schematic representation of lipid A modification by LpxO and PagP. *Salmonella* lipid A is detected at m/z 1797. Hydroxylation by LpxO increases the molecular mass by 16 Da. (B) Lipid A profile under high Mg^2+^ conditions (10 mM Mg^2+^). Lipid A was extracted from wild-type, Δ*lpxO*, and *lpxO*^L287A^ mutant strains and analyzed by MALDI-TOF MS. The Δ*lpxO* strain exhibited only the unmodified lipid A (1797 m/z). Both the wild-type and *lpxO*^L287A^ mutant displayed the hydroxylated lipid A (1813 m/z), with no significant difference in lipid A profiles between the two strains. (C) To assess the impact of LpxO and LpxO^L287A^ on polymyxin B resistance, the Δ*lpxO* mutant was complemented with *lpxO* or *lpxO*^L287A^ cloned into the arabinose-inducible pBAD33 plasmid, and a minimal inhibitory concentration (MIC) assay was performed. The MIC assay was visualized colorimetrically using 0.015% resazurin (29). GC, growth control; SC, sterility control.

Because LpxO-mediated lipid A hydroxylation contributes to resistance against cationic antimicrobial peptides (CAMPs) such as polymyxin B and colistin (12, 13), we next assessed whether the Leu287Ala substitution affects this phenotype. The minimal inhibitory concentration (MIC) of polymyxin B against the Δ*lpxO* mutant was 2 μg/ml, whereas expression of wild-type LpxO in the Δ*lpxO* mutant restored resistance to 4 μg/ml. Expression of LpxO^L287A^ similarly restored resistance to 4 μg/ml (Fig. 3C). Together, these results indicate that although Leu287 of LpxO is important for interaction with MgtC, it is not required for intrinsic hydroxylase activity or polymyxin B-resistance.

### MgtC promotes the hydroxylation of lipid A by interacting with LpxO

To understand the physiological significance of the MgtC-LpxO interaction, we first tested its effect on lipid A hydroxylation. To examine this in a controllable manner, we constructed a Δ*mgtC* Δ*lpxO* double deletion mutant and provided the *mgtC* and *lpxO* genes on inducible plasmids. *mgtC* was cloned into an IPTG-inducible plasmid, while *lpxO* or *lpxO*^L287A^ was cloned into an arabinose-inducible plasmid. After introducing these two plasmids into the Δ*mgtC* Δ*lpxO* double deletion mutant, we monitored lipid A profiles while inducing MgtC expression with increasing concentrations of IPTG and maintaining basal expression of LpxO or LpxO^L287A^ (Fig. 4A and 4B). Western blot analysis confirmed that MgtC protein levels increased with increasing IPTG concentrations (0 to 500 μM) in N-minimal medium containing 10 mM Mg^2+^ (Fig. 4B) (30). This experimental setup allowed comparative analysis of lipid A hydroxylation levels in response to MgtC expression by MALDI-TOF MS.

**Figure 4.**
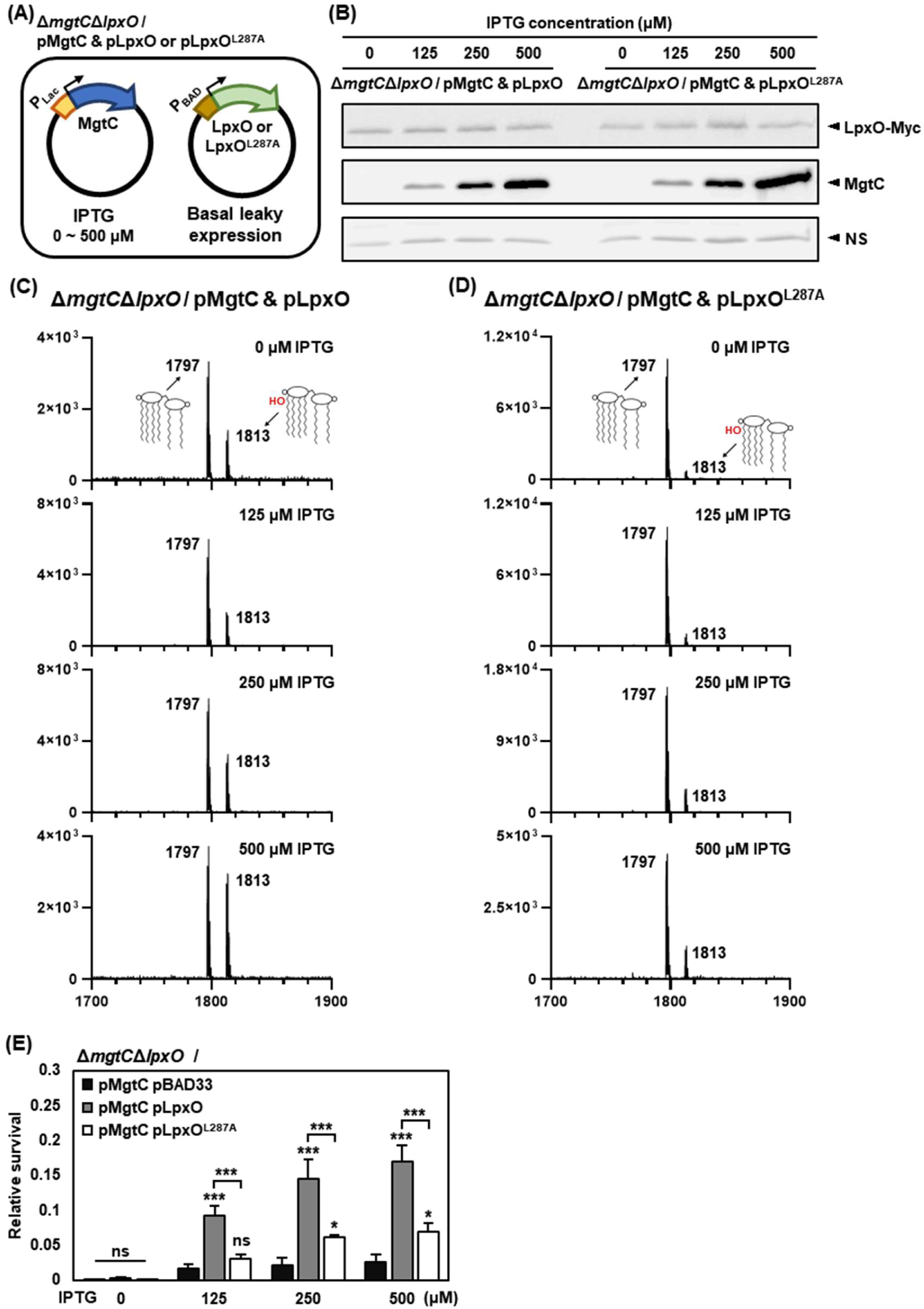
MgtC promotes LpxO-mediated lipid A hydroxylation in a concentration-dependent manner. (A) Schematic of the strain used in this study. The Δ*mgtC*Δ*lpxO* double-knockout mutant was co-transformed with plasmids allowing IPTG-inducible expression of MgtC and basal-level expression of LpxO. Bacteria were grown in N-minimal media containing 10 mM Mg^2+^ for 3 hours and cells were then normalized to OD_600_ = 0.1, after which 0 to 500 μM IPTG was added to induce MgtC, and incubation continued for 2 hours. Cells were then harvested for lipid A extraction and subsequent analysis. (B) Western blot analysis of strains carrying Myc-tagged *lpxO*^WT^ or *lpxO*^L287A^ genes and an IPTG-inducible *mgtC* gene. LpxO was expressed at a basal level, and MgtC expression was induced with different IPTG concentrations. Nonspecific bands in the same blot are shown as a loading control. (C-D) Lipid A profiles of strains expressing LpxO^WT^ (C) or LpxO^L287A^ (D) as MgtC levels increased with increasing IPTG concentrations. (E) Polymyxin B killing assay. After 2 hours of MgtC induction with increasing IPTG concentrations, cells were treated with 2 μg/ml polymyxin B for 1 hour. Survival was calculated as the ratio of CFUs in polymyxin B-treated to untreated samples. Data represent the means ± SD from three independent experiments. Statistical analysis was performed using two-way ANOVA followed by Tukey’s multiple comparisons test.

As MgtC expression increased, the hydroxylated lipid A peak generated by wild-type LpxO increased proportionally (Fig. 4C), whereas the hydroxylated lipid A peak generated by LpxO^L287A^ was significantly lower than that of wild-type LpxO and failed to increase even with elevated MgtC expression (Fig. 4D). Similarly, under N-minimal medium containing 10 μM Mg^2+^, a PhoP/PhoQ-inducing infection-relevant medium (31, 32), LpxO^L287A^ expression exhibited a delayed increase in lipid A hydroxylation compared to wild-type LpxO in response to MgtC expression (Fig. S4).

To assess whether MgtC-mediated enhancement of LpxO-dependent lipid A hydroxylation also correlates with increased resistance to polymyxin B, we performed a polymyxin B killing assay under the same conditions. In the strain expressing basal levels of wild-type LpxO, MgtC induction proportionally enhanced survival in the presence of polymyxin B (PMB). In contrast, the strain expressing LpxO^L287A^ exhibited a significantly lower PMB survival and did not further increase with MgtC induction, consistent with the MALDI-TOF MS results (Fig. 4E). Together, these results demonstrate that MgtC enhances lipid A hydroxylation in a dose-dependent manner via direct interaction with LpxO, and that this regulatory interaction is essential for conferring resistance to polymyxin B.

### The *lpxO*^L287A^ mutation fails to induce polymyxin B resistance under an MgtC-inducing condition

In the above section, we found that MgtC interacts with LpxO via the Leu287 residue and promotes lipid A hydroxylation, thereby conferring resistance to polymyxin B. To address this regulation in a physiological context, we constructed a chromosomal mutant in which the Leu287 residue of LpxO is substituted with Ala, and tested whether this chromosomal *lpxO*^L287A^ mutant behaved as expected with respect to polymyxin B resistance. Because the *mgtC* gene is regulated by the PhoP/PhoQ two-component system, which is activated in Mg^2+^-limiting conditions (32), we performed a time-course polymyxin B killing assay under Mg^2+^-limiting conditions to determine whether the chromosomal *lpxO*^L287A^ mutant exhibits altered resistance when MgtC is endogenously expressed (Fig. 5A). Wild-type, Δ*mgtC*, Δ*lpxO*, and *lpxO*^L287A^ *Salmonella* strains were grown in N-minimal medium containing 10 mM Mg^2+^ (PhoP-repressing) and then transferred to the same medium containing 10 μM Mg^2+^ (PhoP-activating), a condition that induces *mgtC* expression over time. Bacterial samples were harvested at 1-hour intervals from 3 to 6 hours post-inoculation, washed with PBS, and treated with polymyxin B at a final concentration of 2 μg/ml for 1 hour. Following treatment, cells were serially diluted and plated on LB agar to determine CFUs (Fig. 5B).

**Figure 5.**
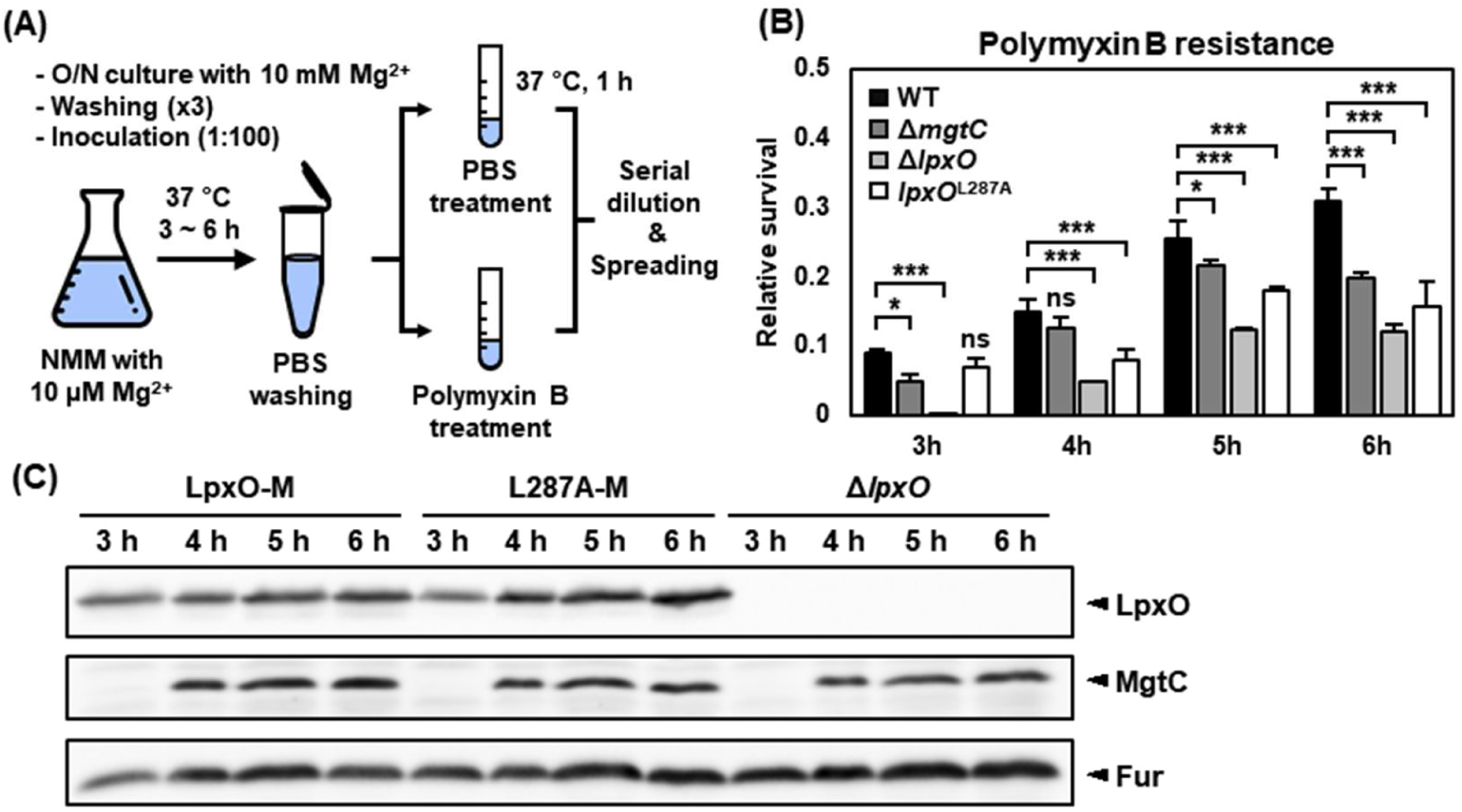
The *lpxO*^L287A^ mutation fails to induce polymyxin B resistance, phenocopying the *mgtC* mutant. (A) Schematic diagram of the time-course polymyxin B killing assay. Bacterial cultures were transferred from 10 mM to 10 μM Mg^2+^ in NMM, and survival following polymyxin B treatment was assessed at hourly intervals from 3 to 6 hours. (B) Results of the time-course polymyxin B killing assay. Data represent means ± SD of three independent experiments. Statistical analysis was performed using two-way ANOVA followed by Tukey’s multiple comparisons test. (C) Western blot analysis of MgtC and LpxO expression in strains with a chromosomal, C-terminally Myc-tagged *lpxO* or *lpxO*^L287A^ gene, and in the Δ*lpxO* strain, during the time-course experiment. LpxO was consistently detected at all time points, whereas MgtC expression became detectable after 4 hours.

In the wild-type strain, polymyxin B survival increased over time from 3 to 6 hours. By contrast, the Δ*lpxO* mutant exhibited significantly lower survival than the wild-type across all time points (Fig. 5B). Interestingly, the *lpxO*^L287A^ mutant exhibited comparable survival to wild-type at 3 hours. However, from 4-5 hours onward, as *mgtC* expression began to increase (Fig. 5C), the *lpxO*^L287A^ mutant became significantly more susceptible to polymyxin B than wild-type and remained susceptible through 6 hours (Fig. 5B). Similarly, the Δ*mgtC* mutant did not show a clear survival difference from wild-type at 3 and 4 hours but displayed a significant decrease at 5 and 6 hours, to a degree similar to that observed in the *lpxO*^L287A^ mutant (Fig. 5B). As control experiments, the *lpxO*^L287A^ mutant exhibited increased susceptibility only under low Mg^2+^ conditions, not under high Mg^2+^ conditions (Fig. S5). In contrast, Δ*lpxO* mutant and three additional *lpxO* chromosomal mutants carrying substitutions in active-site residues (*lpxO*^H155A^, *lpxO*^H166A^, or *lpxO*^H202A^; Fig. S2) exhibited susceptibility under both high and low Mg^2+^ conditions, corresponding to PhoP-repressing and PhoP-activating conditions, respectively (Fig. S5). These results suggest that the interaction between MgtC and LpxO is necessary to achieve optimal polymyxin B resistance under an infection-relevant condition.

### MgtC-LpxO interaction is required for full lipid A hydroxylation inside macrophages, serum resistance, and mouse virulence

Given that MgtC promotes LpxO-dependent lipid A hydroxylation and that *mgtC* is one of the highly expressed genes inside macrophages (33), we further investigated whether this regulation occurs during *Salmonella* infection within macrophages. To explore this, J774A.1 macrophage-like cells were infected with wild-type, Δ*mgtC*, Δ*lpxO*, or *lpxO*^L287A^ *Salmonella* strains for 6 hours, and lipid A was extracted and analyzed by MALDI-TOF MS (Fig. 6A). Both the PhoP/PhoQ and PmrA/PmrB two-component systems control lipid A modification during infection. PhoP/PhoQ controls LpxO-dependent hydroxylation and PagP-dependent palmitoylation, while PmrA/PmrB controls PmrC-dependent addition of phosphoethanolamine (pEtN) and PbgP-dependent addition of 4-amino-4-deoxy-L-aminoarbinose (L-Ara4N) (34–36). Thus, one can expect to observe MS peaks representing combinations of these modifications from lipid A profiles from *Salmonella* during host infection (Fig. 6A and 6B). In the lipid A profile isolated from wild-type *Salmonella* recovered from infected macrophages, 1) *Salmonella* lipid A was predominantly present as hepta-acylated (2035 m/z) or hydroxylated hepta-acylated (2051 m/z) species and 2) the hydroxylated hepta-acylated species was present at relatively higher abundance than the non-hydroxylated hepta-acylated species (Fig. 6C). These data indicate that PagP-dependent palmitoylation and LpxO-dependent hydroxylation constitute the major lipid A modifications under this condition, and that *Salmonella* exhibits strong LpxO-dependent hydroxylation activity inside macrophages (Fig. 6C). As expected, the MS peak corresponding to hydroxylated lipid A species was completely absent in the Δ*lpxO* mutant, confirming that the peaks indicated by red arrows are derived from LpxO-dependent lipid A hydroxylation (Fig. 6B and 6C). Interestingly, both the Δ*mgtC* and *lpxO*^L287A^ mutants exhibited residual levels of lipid A hydroxylation, but these levels were substantially reduced relative to the wild-type strain (Fig. 6C). These data indicate that the MgtC-LpxO interaction is required for full lipid A hydroxylation inside macrophages.

**Figure 6.**
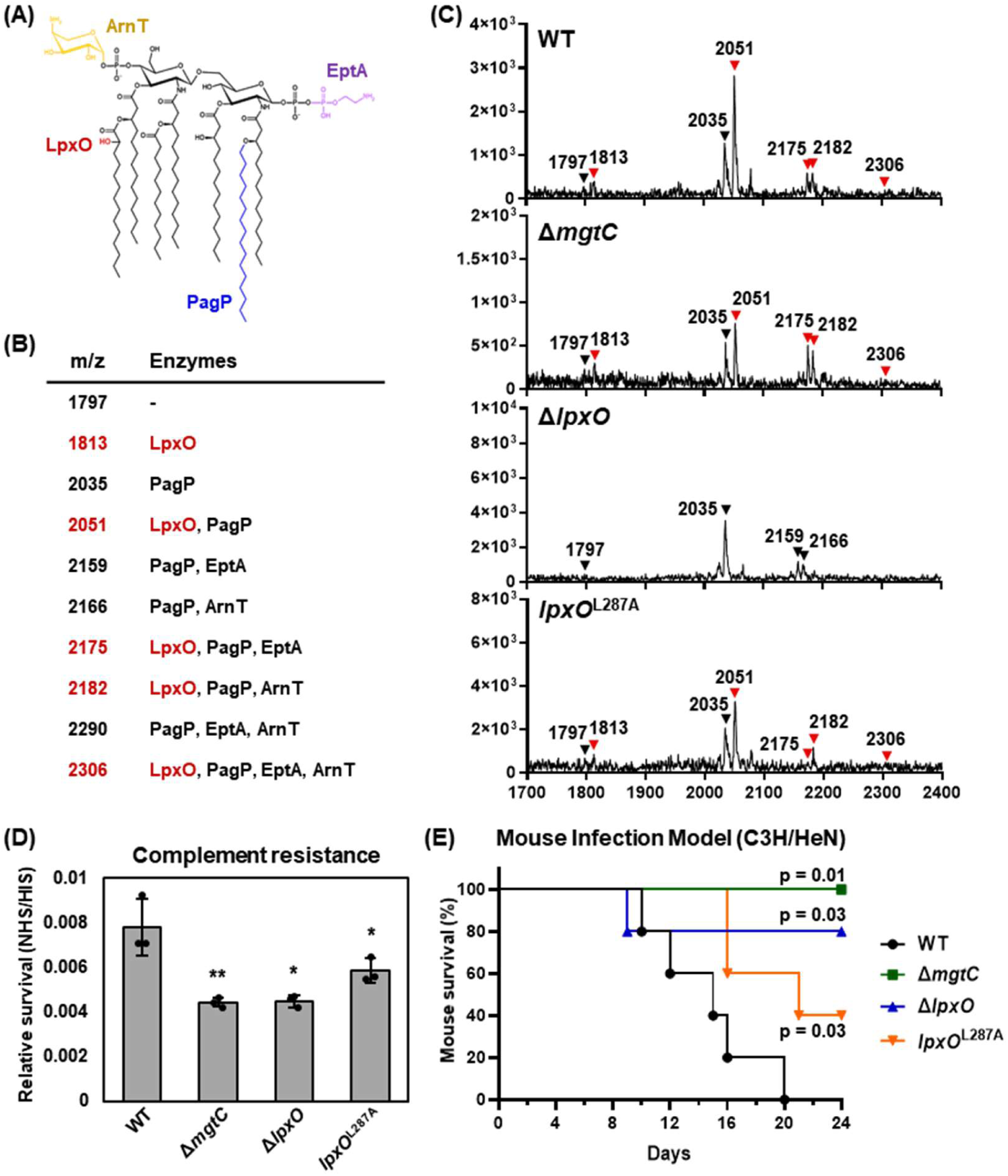
The *lpxO*^L287A^ mutation attenuates lipid A hydroxylation inside macrophages, serum resistance, and virulence in systemic infection. (A) Schematic diagram of PhoP/PhoQ- and PmrA/PmrB-regulated lipid A modifications. LpxO adds a hydroxyl group (+16 m/z) and PagP adds a palmitoyl group (+238 m/z) to lipid A. ArnT and EptA are regulated by the PmrA/PmrB system and add 4-amino-4-deoxy-L-arabinose (+131 m/z) and phosphoethanolamine (+123 m/z) to lipid A, respectively, conferring a positive charge. (B) Changes in lipid A molecular mass resulting from each lipid A-modifying enzyme. (C) J774A.1 cells were infected with wild-type, Δ*mgtC*, Δ*lpxO*, and *lpxO*^L287A^ mutant strains for 6 hours, and the lipid A profiles of intracellular *Salmonella* were analyzed by MALDI-TOF MS to assess lipid A modification. LpxO-mediated lipid A modifications are indicated by red arrows. (D) Complement resistance assay. The Δ*lpxO* mutant showed markedly reduced survival relative to wild-type, and the *lpxO*^L287A^ mutant and the Δ*mgtC* mutant exhibited a similarly reduced resistance. NHS, normal human serum; IHS, inactivated human serum. Data represent means ± SD of three independent experiments. Statistical significance was determined by one-way ANOVA followed by Tukey’s test. (E) C3H/HeN mice were injected intraperitoneally with wild-type, Δ*mgtC*, Δ*lpxO*, and *lpxO*^L287A^ *Salmonella* strains, and survival was monitored over 24 days. By day 20, all mice injected with the wild-type strain had died, while all mice injected with the Δ*mgtC* mutant survived (*x*^2^ = 6.087 vs. wild-type). Mice injected with the Δ*lpxO* mutant exhibited reduced virulence, with an 80% survival rate (*x^2^* = 4.485 vs. wild-type), whereas those injected with the *lpxO*^L287A^ mutant showed a 40% survival rate (*x*^2^ = 4.991 vs. wild-type). Statistical significance was determined using the Log-rank (Mantel–Cox) test.

Alterations in lipid A structure and charge are known to affect bacterial resistance to serum-mediated killing, an important component of host innate defense during systemic infection (37, 38). To determine whether the MgtC-dependent increase in lipid A hydroxylation observed inside macrophages affects serum resistance, we next assessed bacterial killing in human serum. Normal human serum (NHS) contains both CAMPs and complement proteins, representing major innate immune defenses encountered in the bloodstream (38). The above bacterial strains were incubated in NHS for 2 hours, and survival was determined relative to heat-inactivated serum (IHS) controls. Both the Δ*mgtC* and *lpxO*^L287A^ mutants exhibited significantly lower serum resistance than the wild-type strain (Fig. 6D). As a control, the Δ*lpxO* mutant also showed reduced survival, similar to the Δ*mgtC* and *lpxO*^L287A^ mutants (Fig. 6D). These results indicate that both LpxO function and its interaction with MgtC contribute to complement resistance.

Next, to assess the effect of the MgtC-LpxO interaction on mouse virulence, we conducted a mouse virulence assay using C3H/HeN mice. Approximately 10^3^ CFUs of *Salmonella* were injected intraperitoneally, and mouse survival was monitored for 24 days. Mice infected with the wild-type strain showed a decrease in survival starting from day 9, reaching 0% survival by day 20 (Fig. 6E). In contrast, mice infected with the Δ*lpxO* strain showed a slight decrease in survival at day 8, with no further decline thereafter, maintaining 80% survival through day 24 (Fig. 6E). Survival of mice infected with the *lpxO*^L287A^ mutant strain was significantly attenuated compared to wild-type and fell between that of Δ*lpxO* and wild-type, reaching 40% by the end of the experiment (Fig. 6E). As a control, mice infected with Δ*mgtC* strain were avirulent, consistent with previous findings that MgtC is required for virulence through interactions with additional targets (18–22). Together, these findings indicate that LpxO-mediated lipid A hydroxylation is important for *Salmonella* virulence, and that full lipid A hydroxylation and virulence require the interaction between MgtC and LpxO.

## Discussion

We established here that the *Salmonella* virulence factor MgtC interacts with the lipid A-modifying enzyme LpxO identified from a bacterial two-hybrid screen (Fig. 1). The coexistence of MgtC and LpxO among gram-negative bacterial pathogens further supports the rationale that MgtC controls lipid A modification via LpxO (Table 1). MgtC interacts with LpxO through a transmembrane helix-helix interaction between MgtC TM3 and the C-terminal helix of LpxO, and LpxO Leu287 was identified as a key residue for this interaction (Fig. 2 and S1). MgtC binding to LpxO promotes lipid A hydroxylation, and substitution of the Leu287 residue decreased both the MgtC-LpxO interaction and, consequently, lipid A hydroxylation (Fig. 4). This decrease in lipid A hydroxylation also reduced resistance to the antimicrobial peptide polymyxin B, as well as resistance to serum-mediated killing and mouse virulence (Fig. 6). Thus, these data indicate that the MgtC-LpxO interaction is required for *Salmonella* pathogenesis by promoting lipid A hydroxylation.

Interestingly, we noticed that although MgtC activates lipid A hydroxylation (Fig. 4), (i) hydroxylated lipid A species were detected under all tested conditions, including high Mg^2+^ (PhoP-repressing conditions), and (ii) the *lpxO* L287A substitution mutant or the *mgtC* mutant each exhibited a significant but partial decrease in hydroxylated lipid A species (Fig. 4 and 6). A similar phenomenon has been previously described that *lpxO* transcription is induced in low Mg^2+^ (PhoP-inducing conditions), but hydroxylated lipid A species were detected in both low and high Mg^2+^ conditions (PhoP-inducing and PhoP-repressing, respectively), and even in a *phoP* mutant (6). These data indicate a basal level of *lpxO* expression independent of PhoP, and the existence of additional regulatory pathways (Fig. S6) (6). LpxO is an oxygen-dependent hydroxylase (28), and its expression is also regulated by the aerobic transcriptional regulators Fnr and ArcA in response to oxygen availability (39). Studies using intracellular infection models have shown that *Salmonella*-infected tissues experience substantial drops in oxygen tension (40). Similarly, the phagosomal environment within macrophages is known to be micro-oxygenic or hypoxic (33, 41), conditions that could compromise the activity of oxygen-dependent enzymes such as LpxO. We speculate that under such conditions, *Salmonella* employs MgtC to compensate for the reduced activity of LpxO. Within the intracellular niche, MgtC may help preserve LpxO activity through direct interaction, thereby maintaining lipid A hydroxylation.

MgtC is an inner-membrane protein with five transmembrane helices and a cytoplasmic region containing a ligand-binding ACT domain, the nature of whose ligand is still unclear (26). Here, we found that MgtC TM3 is involved in the interaction with the C-terminal helix of LpxO via a helix-helix interaction (Fig. 7). LpxO has two transmembrane helices, one at each end, connected by a long cytoplasmic loop containing key residues for lipid A hydroxylase activity (H155, H166, and H202) (11). Initial bacterial two-hybrid screening identified only the C-terminal 55 amino acids, including the C-terminal helix, as sufficient for the interaction, and we further verified that a minimal fragment corresponding to just the 24 amino acid C-terminal helix is sufficient and the L287A substitution within the helix weakens the MgtC-LpxO interaction and consequently lipid A hydroxylation (Fig. 2 and 4). Given that the active site for hydroxylation resides within the cytoplasmic loop, a question remains as to how the helix-helix interaction between MgtC-LpxO activates hydroxylation activity. One clue comes from the observation that other Fe^2+^/α-ketoglutarate-dependent hydroxylases lack these two transmembrane helices, suggesting that correct membrane localization of LpxO via these helices may determine substrate specificity toward lipid A (11). Our finding that LpxO lacking either transmembrane helix completely loses lipid A hydroxylase activity (Fig. S3) further supports the notion that correct localization to the inner membrane is critical for full lipid A hydroxylase activity, and that MgtC binding may facilitate this process through its helix-helix interaction.

**Figure 7.**
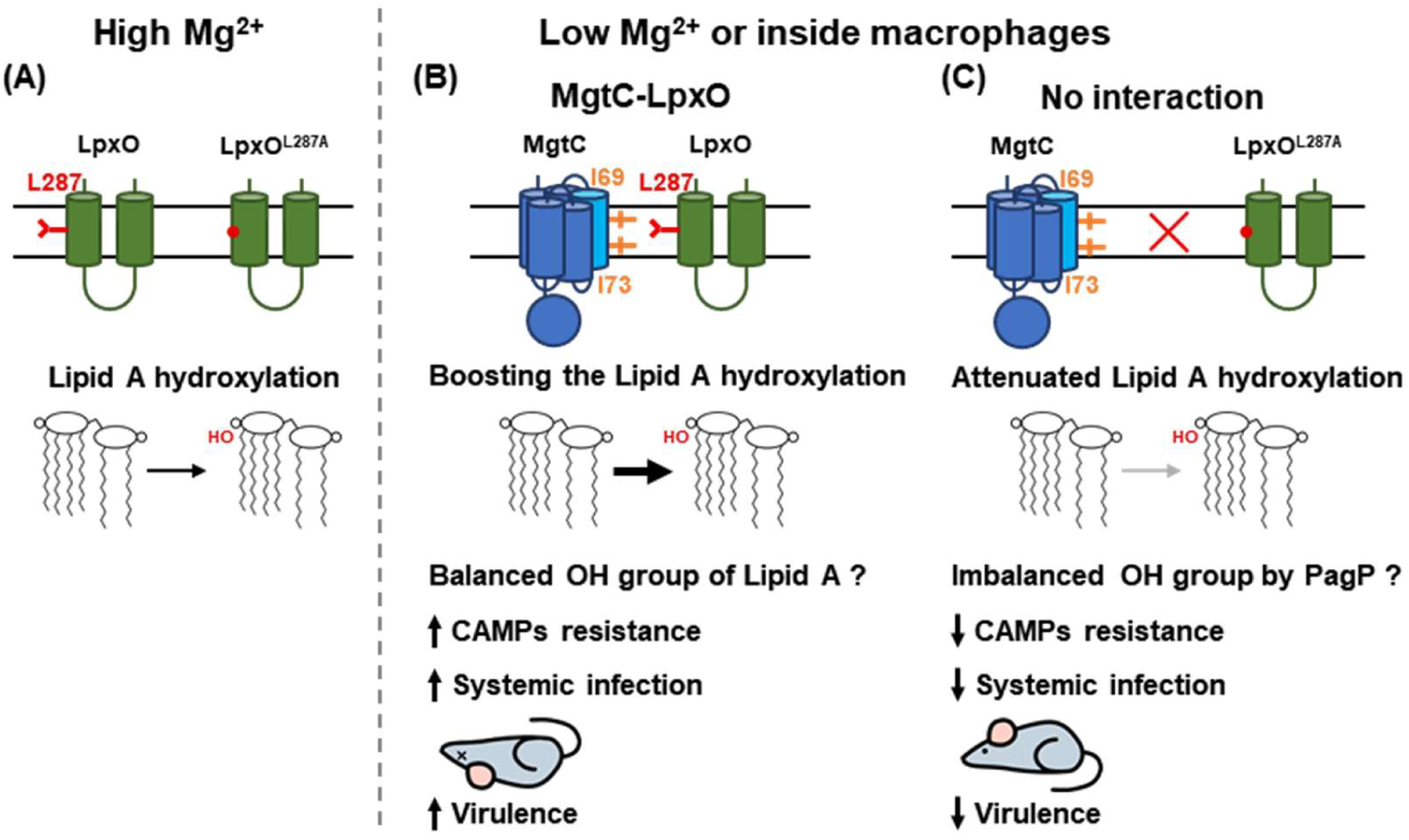
A model of MgtC-LpxO interaction governing lipid A modification. (A) Under a high Mg^2+^ condition where MgtC is not expressed, both wild-type LpxO and LpxO^L287A^ catalyze lipid A hydroxylation at comparable levels. (B) Under low Mg^2+^ or intramacrophage conditions characterized by limited magnesium and oxygen availability, MgtC is highly expressed and LpxO activity is presumed to be reduced. However, MgtC interaction to LpxO enhances LpxO-mediated lipid A hydroxylation. This maintains hydroxyl group balance, supports lipid A palmitoylation by PagP, and increases resistance to CAMPs, ultimately promoting systemic infection and full virulence. (C) In contrast, the LpxO^L287A^ mutant shows reduced interaction with MgtC, resulting in reduced hydroxylation efficiency. This leads to disrupted hydroxyl group balance, reduced lipid A palmitoylation, decreased resistance to CAMPs, and diminished systemic infection and attenuated virulence.

In support of the idea that LpxO activity can be modulated by other proteins, it has also been reported that VisP also directly interacts with LpxO. VisP is required for *Salmonella* survival against multiple stressors, as well as for intramacrophage survival and virulence in a murine infection model (42). However, unlike the MgtC-LpxO interaction that promotes lipid A hydroxylation, VisP inhibits it. Notably, 1) VisP is a periplasmic protein containing an oligonucleotide/oligosaccharide-binding (OB) domain, which potentially interacts with N-acetylmuramic acid (NAM) in the peptidoglycan, and 2) LpxO has no residues extending into the periplasmic space; thus, how VisP interacts with LpxO to inhibit lipid A hydroxylation remains unclear, given that lipid A hydroxylation occurs on the cytoplasmic face of the inner membrane (43). Hydroxylated lipid A species are strongly induced and predominate inside macrophages (Fig. 6) (6), and the Δ*lpxO* mutant is sensitive to polymyxin B and attenuated in mouse virulence, indicating that LpxO-mediated lipid A hydroxylation is required for resistance to antimicrobial peptides and for mouse virulence. This raises another question of how VisP-mediated inhibition of lipid A hydroxylation contributes to *Salmonella* virulence, given that *visP* is also required for virulence. Currently, we speculate that MgtC and VisP may act at different stages of infection, a scenario that needs to be further investigated.

Both LpxO-mediated hydroxylation and PagP-mediated palmitoylation act on lipid A under low Mg^2+^, a PhoP/PhoQ-activating condition. There is a possible explanation linking MgtC-mediated lipid A hydroxylation to the palmitoylation process catalyzed by PagP. Under low magnesium conditions (10 μM Mg^2+^), where *pagP* is transcriptionally upregulated, the strain expressing LpxO^L287A^ displayed reduced hydroxylation due to weakened MgtC-LpxO interaction, but palmitoylation levels remained unchanged (Fig. S4E). Although PagP is capable of palmitoylating both hydroxylated and non-hydroxylated lipid A species, our data suggest that the overall efficiency or distribution of palmitoylation is influenced by the hydroxylation state of lipid A. This observation points to a potential balancing mechanism in lipid A modification (Fig. S7). These lipid A modifications increase membrane rigidity or alter the membrane’s net charge to reduce affinity for CAMPs (4). However, LpxO-mediated lipid A hydroxylation presents an interesting paradox, as it is expected to increase the negative charge of lipid A. Notably, palmitoylation esterifies the free hydroxyl group on lipid A, which would be expected to reduce intermolecular hydrogen bonding while increasing acyl-chain packing density. One hypothesis to resolve this paradox is that LpxO compensates for the hydroxyl group consumed by PagP-mediated palmitoylation, thereby restoring the capacity for hydrogen bonding and maintaining a balance that stabilizes the membrane (44). Together, these two modifications may contribute to outer membrane barrier function through complementary effects that together reduce susceptibility to CAMPs (44), providing indirect evidence that LpxO replenishes the hydroxyl group consumed by PagP to maintain balance in lipid A modification (Fig. S7).

In conclusion, our study uncovers a new role for MgtC in lipid A modification, demonstrating its direct involvement in enhancing LpxO-mediated hydroxylation. The *lpxO*^L287A^ mutant, which retains intrinsic hydroxylation activity but weakens the interaction with MgtC, exhibited a significantly reduced resistance to host antimicrobial peptides and attenuated virulence, supporting the notion that the MgtC-LpxO interaction is a crucial determinant of systemic infection. This interaction is essential for resistance to CAMPs and complement during systemic infection, thereby contributing to the overall virulence of *Salmonella*. These findings offer new insights into *Salmonella* pathogenicity and reinforce the importance of outer layer modification by a bacterial virulence factor in adaptation and survival inside the host.

## Materials and Methods

### Bacterial strains and growth conditions

Bacterial strains used in this study are listed in Table S2. All *Salmonella* strains used in this study were derived from the wild-type *Salmonella enterica* serovar Typhimurium strain 14028s (45). Bacteria were grown at 37°C in Luria-Bertani (LB) medium or N-minimal medium (pH 7.7) supplemented with 10 mM or 10 μM of MgCl_2_ (46).

### Construction of Δ*lpxO* and Δ*mgtC*Δ*lpxO* mutant strains

We used the Δ*mgtC* (EL4) strain from a previous study (19). Each deletion mutant was constructed via the one-step inactivation method (47). Using the pKD4 plasmid, a kanamycin-resistance cassette flanked by 40-nt of *lpxO* homology regions was amplified by PCR (NJP036/NJP037) and integrated into the chromosome of the wild-type strain and Δ*mgtC* mutant. After inactivation, the kanamycin-resistance cassette was removed by thermal induction of FLP recombinase from the pCP20 plasmid (47).

### Construction of the *lpxO*^L287A^, *lpxO*^H155A^, *lpxO*^H166A^, and *lpxO*^H202A^ mutants, and 8xMyc tagging of *lpxO* and *lpxO*^L287A^

To generate mutants carrying a missense mutation in the *lpxO* gene, we used a counterselection method based on the fusaric acid sensitivity (48, 49).

A Tn10 tetracycline-resistance cassette with 40-bp overhangs of the *lpxO* gene was amplified by PCR using the NJP099/NJP100 primer pair and integrated into the chromosome using λ recombinase from the pKD46 plasmid. We then replaced the cassette with PCR products containing the desired mutations (primer pairs: NJP101/NJP102 for L287A, NJP109/NJP110 for H155A, NJP107/NJP108 for H166A, and NJP105/NJP106 for H202A), also using λ recombinase, and isolated colonies with the cassette removed via fusaric acid selection. Each mutation was confirmed by sequencing.

For tagging *lpxO* with 8xMyc, we followed a similar approach. We transformed the wild-type and *lpxO*^L287A^ mutant strains with the pKD46 plasmid, amplified the 40 bp overhangs near the *lpxO* C-terminus (using NJP171/NJP172 primer pair), and integrated the tetracycline cassette. Subsequently, we integrated the PCR product encoding an 8xMyc tagged C-terminus, amplified from the pBOP508 plasmid (50), into the chromosome using λ recombinase, isolated colonies after cassette removal with fusaric acid, and confirmed the presence of the 8xMyc tag by sequencing.

### Construction of plasmids

For the BACTH assay to identify the interacting residues of LpxO with MgtC, the plasmids pKNT25-*lpxO*^1-250^, pKT25-*lpxO*^28-302^, pKT25-*lpxO*^278-302^, pKT25-*lpxO*^278-302-R278A^, pKT25-*lpxO*^278-302-T279A^, pKT25-*lpxO*^278-302-L280A^, pKT25-*lpxO*^278-302-Y281A^, pKT25-*lpxO*^278-302-T282A^, pKT25-*lpxO*^278-302-L284A^, pKT25-*lpxO*^278-302-K285A^, pKT25-*lpxO*^278-302-Y286A^, pKT25-*lpxO*^278-302-L287A^, pKT25-*lpxO*^278-302-V288A^, pKT25-*lpxO*^278-302-I289A^, pKT25-*lpxO*^278-302-I292A^, pKT25-*lpxO*^278-302-F293A^, pKT25-*lpxO*^278-302-I296A^, pKT25-*lpxO*^278-302-I297A^, pKT25-*lpxO*^278-302-L298A^, pKT25-*lpxO*^278-302-F299A^, pKT25-*lpxO*^278-302-S300A^, pKT25-*lpxO*^278-302-L301A^ and pKT25-*lpxO*^278-302-L302A^ were constructed as follows: each DNA fragment was amplified by PCR using the primer pairs NJP084/NJP085 (for *lpxO*^1-250^), NJP045/NJP066 (for *lpxO*^28-302^), NJP042/NJP066 (for *lpxO*^278-302^), NJP046 to NJP056/NJP066 (for *lpxO*^278-302-R278A^ to *lpxO*^278-302-I289A^), and NJP042/NJP057 to NJP065 (for *lpxO*^278-302-I292A^ to *lpxO*^278-302-L302A^), using 14028s genomic DNA as a template.

On the other hand, to identify the interacting residues of MgtC with LpxO, the plasmids pUT18-*mgtC*-TM1, pUT18C-*mgtC*-TM2, pUT18-*mgtC*-TM3, pUT18C-*mgtC*-TM4, pUT18-*mgtC*-TM5, pUT18-*mgtC*-TM3^Q68A^, pUT18-*mgtC*-TM3^I69A^, pUT18-*mgtC*-TM3^V70A^, pUT18-*mgtC*-TM3^S71A^, pUT18-*mgtC*-TM3^I73A^, pUT18-*mgtC*-TM3^F75A^, and pUT18-*mgtC*-TM3^L76A^ were constructed as follows: each DNA fragment was amplified by PCR using the primer pairs NJP111/NJP112 (for *mgtC*-TM1), NJP113/NJP114 (for *mgtC*-TM2), NJP115/NJP116 (for *mgtC*-TM3), NJP117/NJP118 (for *mgtC*-TM4), NJP119/NJP120 (for *mgtC*-TM5), NJP121/NJP116 (for *mgtC*-TM3^Q68A^), NJP122/NJP116 (for *mgtC*-TM3^S71A^), NJP115/NJP123 (for *mgtC*-TM3^F75A^), NJP225/NJP116 (for *mgtC*-TM3^I69A^), NJP226/NJP116 (for *mgtC*-TM3^V70A^), NJP228/NJP116 (for *mgtC*-TM3^I73A^), and NJP115/NJP230 (for *mgtC*-TM3^L76A^), using 14028s genomic DNA as a template. All PCR products were digested with XbaI and KpnI and cloned into pKNT25, pKT25, pUT18, and pUT18C plasmids, respectively.

For MgtC and LpxO protein expression, pUHE21-*mgtC* and pBAD33-*lpxO,* pBAD33_*lpxO*^28-302^, pBAD33_*lpxO*^1-277^, or pBAD33-*lpxO*^L287A^ (as well as pBAD33-*lpxO*-8xMyc or pBAD33-*lpxO*^L287A^-8xMyc) were constructed as follows. Insert DNAs were amplified by PCR using the primer pairs NJP233/NJP159 (for *mgtC*) and NJP078/NJP034 (for *lpxO*, *lpxO*-8xMyc, *lpxO*^L287A^, and *lpxO*^L287A^-8xMyc), NJP315/NJP316 (for pBAD33_*lpxO*^28-302^), and NJP078/NJP081 (for pBAD33_*lpxO*^1-277^) using genomic DNA from wild-type 14028s or the 8xMyc-tagged *lpxO* and *lpxO*^L287A^ strains (NJ207 and NJ209, respectively) as templates. The amplified *mgtC* gene was digested with BamHI and PstI and cloned into pUHE21, and the amplified *lpxO, lpxO*^28-302^, *lpxO*^1-277^, or *lpxO*^L287A^ genes were digested with KpnI, HindIII, or XbaI and cloned into pBAD33.

For the Co-IP assay, pUHE21-MgtC-FLAG was constructed using the NJP233/NJP234 primer pair.

### Finding MgtC interaction partners using BACTH assay

To identify the protein-protein interaction *in vivo*, a BACTH assay was performed (25). Chromosomal DNA of *S*. Typhimurium 14028s was extracted using G-spin^TM^ genomic DNA extraction kit (iNtRON, Korea) and digested with Sau3AI restriction enzyme (Enzynomics, Korea). DNA fragments ranging from 500 to 2000 bp were purified via gel electrophoresis. The DNA libraries were cloned into BamHI-digested pKT25 and pKNT25 plasmids, which were then transformed into *E. coli* BTH101 containing the pUT18 plasmid with cloned *mgtC* as bait (20). Transformants were spread on MacConkey agar plates with 0.5 mM IPTG, 1% maltose, 50 μg/ml ampicillin, 50 μg/ml kanamycin, and incubated at 30°C for 48 to 72 hours. Red colonies were re-streaked 2 to 3 times. Library DNA was amplified by colony PCR and confirmed by sequencing.

To identify interacting residues of MgtC and LpxO, site-directed alanine substitution mutants were constructed and subjected to BACTH analysis. Five microliters of overnight cultures of BTH101 strains containing the respective plasmid pairs were spotted onto MacConkey or LB X-gal agar plate (supplemented with 40 μg/ml X-gal, 0.5 mM IPTG, 50 μg/mL ampicillin, and 50 μg/mL kanamycin) and incubated at 30 °C for 48–72 h.

### Immunoprecipitation assay

The interaction between MgtC and LpxO was investigated in *Salmonella* strains expressing either wild-type LpxO, LpxO-8xMyc, or LpxO^L287A^-8xMyc, each containing the pUHE21-*mgtC*-FLAG plasmid, where MgtC is under IPTG-inducible expression. Cells were grown in 50 ml of N-minimal medium supplemented with 10 mM MgCl_2_ for 3.5 hours at 37°C, followed by induction of MgtC expression with 500 μM IPTG for 2 hours. After induction, cells were harvested and normalized by OD_600_. For cell lysis, pellets were washed once with 1x TBS and lysed by sonication in TBS buffer. For the pull-down assay, 40 μl of the crude extracts were kept as input controls, while 200 μl of the protein extracts were mixed with beads conjugated to anti-FLAG or anti-Myc antibodies. The samples were incubated overnight at 4°C with gentle mixing. After washing, the bound proteins were eluted in SDS sample buffer, separated on a 12% SDS-polyacrylamide gel, and analyzed by western blotting with anti-FLAG antibody (1:10,000 dilution, Sigma-Aldrich, Cat#F2426) and anti-Myc antibody (1:10,000 dilution, MBL life science, Cat#M192-3).

### Distribution and conservation of MgtC and LpxO

All MgtC and LpxO amino acid sequences were obtained from Uniprot database (www.uniprot.org, supplementary data). Partial protein sequences were excluded, and true MgtC sequences were identified using MUSCLE alignment in MEGA11, following established criteria (26, 27).

### Lipid A extraction and MALDI-TOF MS/MS

Bacterial lipid A was extracted as described (51, 52) with a few modifications. First, bacterial cells were grown in NMM with 10 mM Mg^2+^ overnight. These pre-cultured bacteria were diluted 1:100 and inoculated in NMM with 10 mM or 10 μM MgCl_2_ for 4 to 5 hours (OD_600_ = 0.3 to 0.4). Cells were harvested and washed with phosphate-buffered saline (PBS). The remaining pellets were resuspended in 500 μl of Tri-Reagent, vortexed vigorously and incubated at room temperature for 10 min. One hundred microliters of chloroform were added, vortexed vigorously, and incubated for another 10 min. The mixtures were centrifuged at 13,000 rpm for 10 min and the aqueous phase was transferred into a new microcentrifuge tube. One hundred microliters of distilled water were added to the organic phase, mixed by vortex, incubated at room temperature for 10 min, and centrifuged at 13,000 rpm for 10 min. Upper aqueous phase was collected and combined with same microcentrifuge tube as described above. This water extraction step was performed twice. These aqueous phases were dried by SpeedVac and finally lipopolysaccharide (LPS) was extracted.

LPS was then dissolved in mild acid hydrolysis buffer (12.5 mM sodium acetate, 1% SDS, pH 4.5), incubated at room temperature for 30 min, boiled at 100 °C for 1 hour for releasing the lipid A, and dried by SpeedVac again. Lipid A was then washed with the mixture of 100 μl of distilled water and 500 μl of acidified 95% ethanol (prepared by combining 100 μl of 4 M HCl with 20 ml of 95% ethanol). The resuspensions were centrifuged at 2100 g for 10 min and the supernatant was discarded very carefully. The white solid pellets were washed twice with 500 μl of 95% ethanol. The supernatant was removed very carefully, and the lipid A pellet was air-dried.

Lipid A pellet was dissolved in the chloroform and methanol (3:1) solution for matrix-assisted laser desorption-ionization time-of-flight (MALDI-TOF) mass spectrometry analysis. MALDI-TOF analysis was outsourced to GBSA, Korea (www.gbsa.or.kr). Two microliters of dissolved lipid A sample were mixed with an equal volume of matrix solution (10 mg/ml of 2,5-Dihydroxybenzoic acid (DHB) in 80% acetonitrile (ACN) with 0.1% trifluoroacetic acid (TFA)) directly on the MALDI target and air-dried under vacuum. The analysis was performed in negative-ion linear mode, with an m/z detection range of 1,000 to 4,000, a laser repetition rate of 2,000 Hz, and an accelerating voltage of 20 kV using an Autoflex maX (Bruker Daltonics, USA).

### Minimal inhibitory concentration assay

The strains were cultured in Mueller-Hinton (MH) broth. Polymyxin B was diluted in MH broth to the 4× top concentration desired in the test. Using a multi-pipette, 50 μl of MH broth was dispensed into a 96-well plate and 50 μl of diluted antibiotics were mixed into the wells of column 1 by pipetting up and down 6-8 times. Fifty microliters of sample from column 1 were withdrawn and added to the next column. This procedure was repeated through column 10, such that each well contained 50 μl of serially diluted antibiotic. Overnight-grown bacteria were diluted to the concentration suggested by CLSI guideline (about 1/100 diluted 0.5 McFarland) and dispensed in 50 μl volumes into the plate except column 11 (sterilized control). This 96-well plate was incubated overnight at 37 °C, added 30 μl of 0.015% resazurin and incubated for 2 to 4 hours to distinguish the MIC (29).

### Western blot

Cells were grown for 5 hours in N-minimal medium supplemented with either 10 mM or 10 μM Mg^2+^ at 37°C. Cultures were normalized based on OD_600_ and harvested by centrifugation. Crude extracts were prepared by sonication in 1x TBS buffer, and whole-cell lysates were resolved by SDS-PAGE on 12% polyacrylamide gels. Proteins were transferred onto PVDF membranes and blocked with 5% skim milk in PBS. Membranes were incubated overnight at 4°C with either monoclonal anti-Myc antibody (1:5,000 dilution, MBL) for detecting LpxO or polyclonal anti-MgtC antibody (1:10,000 dilution, in-house) for detecting MgtC. After washing, blots were incubated for 1 hour with horseradish peroxidase-conjugated secondary antibodies: anti-mouse IgG (1:10,000 dilution, ThermoFisher) or anti-rabbit IgG (1:10,000 dilution, ThermoFisher), respectively. Protein bands were visualized using the ECL detection system (SuperSignal™ West Femto Maximum Sensitivity Substrate, ThermoFisher).

### Polymyxin B killing assay

The polymyxin B killing assay was performed as described (53) with a few modifications. Briefly, overnight cultures of bacteria grown in NMM with 10 mM MgCl_2_ at 37°C were washed three times with magnesium-free NMM and inoculated (1:100 dilution) into NMM with 10 μM MgCl_2_ at 37°C for 4 to 5 hours. These bacteria were normalized to an OD_600_ of 0.3 to 0.4 and diluted 1:100 in LB broth. Polymyxin B solution was prepared at 2-fold of final concentration in LB broth. Next, 500 μl of diluted bacteria were added to 500 μl of polymyxin B solution and incubated at 37°C for 1 hour with aeration. Polymyxin B treated and untreated bacteria were serially diluted and plated on LB agar plates. The survival rate was calculated as the ratio of CFU of polymyxin B treated samples to CFU of untreated samples.

For the time-course polymyxin B killing assay, overnight cultures grown in NMM with 10 mM MgCl_2_ were washed twice with magnesium-free NMM and inoculated into NMM containing 10 μM MgCl_2_ at 1:100 dilution. Bacterial samples were harvested every hour from 3 to 6 hours post-inoculation, washed with PBS, and adjusted to an OD_600_ of 0.1. Cells were then treated with 2 μg/ml of polymyxin B at 37°C for 1 hour. CFUs were determined by plating serial dilutions on LB agar, and survival rate was calculated.

### Complement resistance assay

The complement resistance assay was conducted based on previously described methods (38, 54) with minor modifications. Normal human serum was purchased from Sigma-Aldrich (catalog no. H4522). To inactivate complement, half of the serum was incubated at 56 °C for 30 minutes. Bacterial overnight cultures grown in NMM supplemented with 10 mM MgCl_2_ at 37 °C were washed three times with magnesium-free NMM and then diluted 1:100 into NMM containing 10 μM MgCl_2_. The cultures were further incubated at 37 °C for 4–5 hours. Cells were adjusted to an OD_600_ of 0.1, washed with PBS, and resuspended. Next, 50 μl of normal human serum or inactivated human serum was mixed with 40 μl of PBS and 10 μl of washed bacterial suspension. The mixtures were incubated at 37 °C for 2 hours. CFUs were determined by plating serial dilutions on LB agar, and the survival rate was calculated as the ratio of CFU of normal human serum treated samples to CFU of inactivated human serum treated controls.

### Gentamicin protection assay

Gentamicin protection assay was conducted using the macrophage-like cell line J774A.1 as described (18). First, J774A.1 cells were cultured in Dulbecco’s modified Eagle’s medium (DMEM) supplemented with 10% fetal bovine serum (FBS) and 5 × 10^5^ cells were seeded in 24-well plates and cultured with 5% CO_2_ at 37°C for overnight. On the other hand, bacteria for infection were grown in LB at 37°C for overnight. Then, seeded J774A.1 were infected with overnight grown bacteria at a multiplicity of infection (MOI) of 20:1, and plates were centrifuged at 1,500 rpm for 5 min at room temperature. These bacteria were plated on LB agar plates to calculate CFUs used for infection (T_0_). Cells were incubated with 5% CO_2_ for 30 min at 37°C to allow phagocytosis, and the extracellular bacteria were washed twice with Dullbecco’s phosphate-buffered saline (DPBS) and killed by incubation for 1 hour within the DMEM containing 10% FBS and 150 μg/ml gentamicin. To measure the CFUs of invasion (T_1_) into J774A.1, wells were treated with PBS containing 0.1% Triton X-100 and lysed cells were plated on LB agar plates at serial dilutions. For measuring intracellular replication, wells were washed with DPBS and DMEM containing 10% FBS and 15 μg/ml gentamicin were added. Cells were cultured another 21 hours and similarly, wells were treated with DPBS containing 0.1% Triton X-100 and lysed cells were plated on LB agar plates at serial dilutions (T_21_). The invasion rates were calculated by dividing the T_1_ by T_0_ and replication rates were calculated by dividing the T_21_ by T_1_. All experiments were conducted in triplicate, and the results presented are representative of at least three independent replicates.

### Extraction of bacterial lipid A within the macrophage

To extract lipid A from bacteria within infected macrophages, J774A.1 cells were cultured in DMEM medium supplemented with 10% FBS in a T25 cell culture flask until reaching confluency (approximately 8.4 × 10^6^ cells), maintained in 5% CO_2_ at 37°C. J774A.1 cells were then infected with overnight-grown bacteria at a MOI of 100:1, followed by centrifugation at 1,500 rpm for 5 min at room temperature. The cells were incubated in 5% CO_2_ at 37°C for 30 min to allow phagocytosis, after which extracellular bacteria were washed twice with DPBS and killed by incubating the cells for 1 hour in DMEM containing 10% FBS and 150 μg/ml gentamicin. After washing the cells with DPBS, DMEM containing 10% FBS and 15 μg/ml gentamicin was added, and the culture was incubated for an additional 6 hours to allow lipid A modification in the bacteria. Following two DPBS washes, the cells were treated with DPBS containing 0.1% Triton X-100 and harvested. The subsequent steps for lipid A extraction followed the lipid A extraction protocol above.

### Mouse infection

Six- to eight-week-old female C3H/HeN mice were inoculated intraperitoneally with ∼10^3^ CFUs of *Salmonella* strains. Mouse survival was followed for 24 days. Virulence assays were conducted three times with similar outcomes and the data correspond to groups of five mice. All animals were housed in a temperature- and humidity-controlled room, in which a 12 h light/12 h dark cycle was maintained. The experimental procedures were approved by the Korea University Institutional Animal Care and Use Committee.

### Real-time quantitative reverse transcription PCR (qRT-PCR)

Total RNA was isolated using the RNeasy kit (Qiagen) according to the manufacturer’s instructions. The purified RNA was quantified using a NanoDrop spectrophotometer (Thermo Fisher). Five hundred nanograms of cDNA were synthesized using a PrimeScript RT reagent kit (TaKaRa). Transcripts were quantified by real-time PCR using the SYBR green PCR master mix (Applied Biosystems) on a StepOnePlus real-time PCR system (Applied Biosystems) with the following PCR protocol: 95°C for 20 sec followed by 40 cycles of 95°C for 2 sec and 60°C for 30 sec. The mRNA levels of each target gene were calculated using a standard curve of *Salmonella* Typhimurium 14028s genomic DNA at known concentrations, and the data were normalized to 16S rRNA levels of the *rrsH* gene. The primers used for the detection of transcripts of each gene are listed in Table S2.

### Molecular structures and protein modelling

Lipid A molecular structures were drawn using ACD/ChemSketch (Freeware, File Version C45E41) and all 3D protein structures were generated by AlphaFold 3 (55) and visualized by PyMOL 3.0 (56).

## Data, Materials, and Software Availability

All data are included in the manuscript and/or supporting information.

## Acknowledgements

This work was supported by Basic Science Research Program through the National Research Foundation of Korea (NRF) funded by the Ministry of Science, ICT and Future Planning [NRF-2022R1A2B5B02002256, NRF-RS-2025-00561488, and NRF-2022R1A4A1025913 to E.-J.L.] and a grant from Korea University. We thank the undergraduate students who contributed to the bacterial two-hybrid screening as part of the summer research program.

## Author contributions

E.-J.L. conceptualized the study, visualized the study, acquired funding, and wrote the original draft. N.C. conducted the investigation, performed formal analysis, and wrote the original draft. S.B. and E.C. conducted the investigation.

## Supporting information

**Figure S1.**
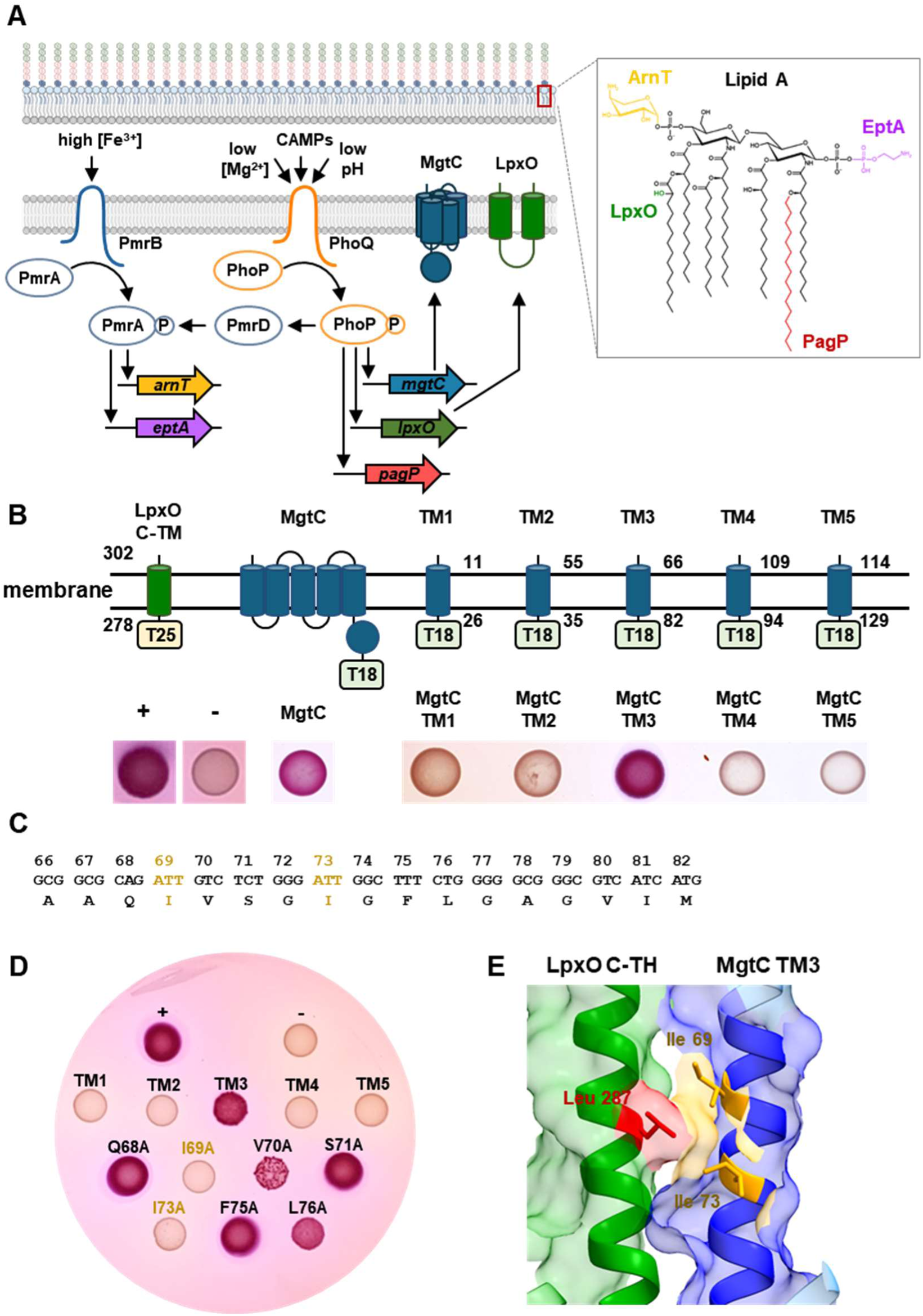
Ile69 and Ile73 in the MgtC 3^rd^ transmembrane helix are required for LpxO interaction. (A) Control of lipid A modification in *Salmonella* Typhimurium. In response to low Mg^2+^, antimicrobial peptides (CAMPs), and pH, the PhoP/PhoQ two-component system controls transcription of the *lpxO* and *pagP* genes, encoding a lipid A hydroxylase and a palmitoyltransferase, respectively. In addition, in response to high Fe^3+^, the PmrA/PmrB two-component system controls transcription of the *pmrC* and *pbgP* genes, adding 4-amino-4-deoxy-L-arbinose (L-Ara4N) or phosphoethanolamine to lipid A, respectively. PhoP also controls production of PmrD, which binds to phosphorylated PmrA and activates PmrC- and PbgP-mediated lipid A modification under PhoP-inducing conditions. (B) Schematic representation of constructs used in the BACTH assay. Each transmembrane helix (TM) of MgtC was cloned into pUT18 or pUT18c plasmids according to its topology, and interaction was assessed by bacterial two-hybrid assay of BTH101 cells coexpressing LpxO C-TM-T25 and C-terminally T18-fused MgtC or MgtC transmembrane domains: TM1 (11–26), TM2 (35–55), TM3 (66–82), TM4 (94–109), and TM5 (114–129). Cells were spotted on MacConkey agar plates containing 1% maltose and 0.5 mM IPTG and incubated at 30°C for 48 hours. Red color indicates a positive interaction. +, positive control (MgtC-MgtR), (1, 2), -, negative control (MgtC-empty vector). (C) Amino acid sequence of MgtC TM3 (66–82). (D) Alanine scanning mutagenesis was performed on the amino acids in TM3 of MgtC, followed by a BACTH assay to test interaction with LpxO’s C-TH. Substitution of I69 or I73 with alanine abolished this interaction. (E) 3D structural model of TM3 of MgtC (blue) and C-TM of LpxO (green). All models were generated by AlphaFold3, and the image was produced using UCSF ChimeraX (3).

**Figure S2.**
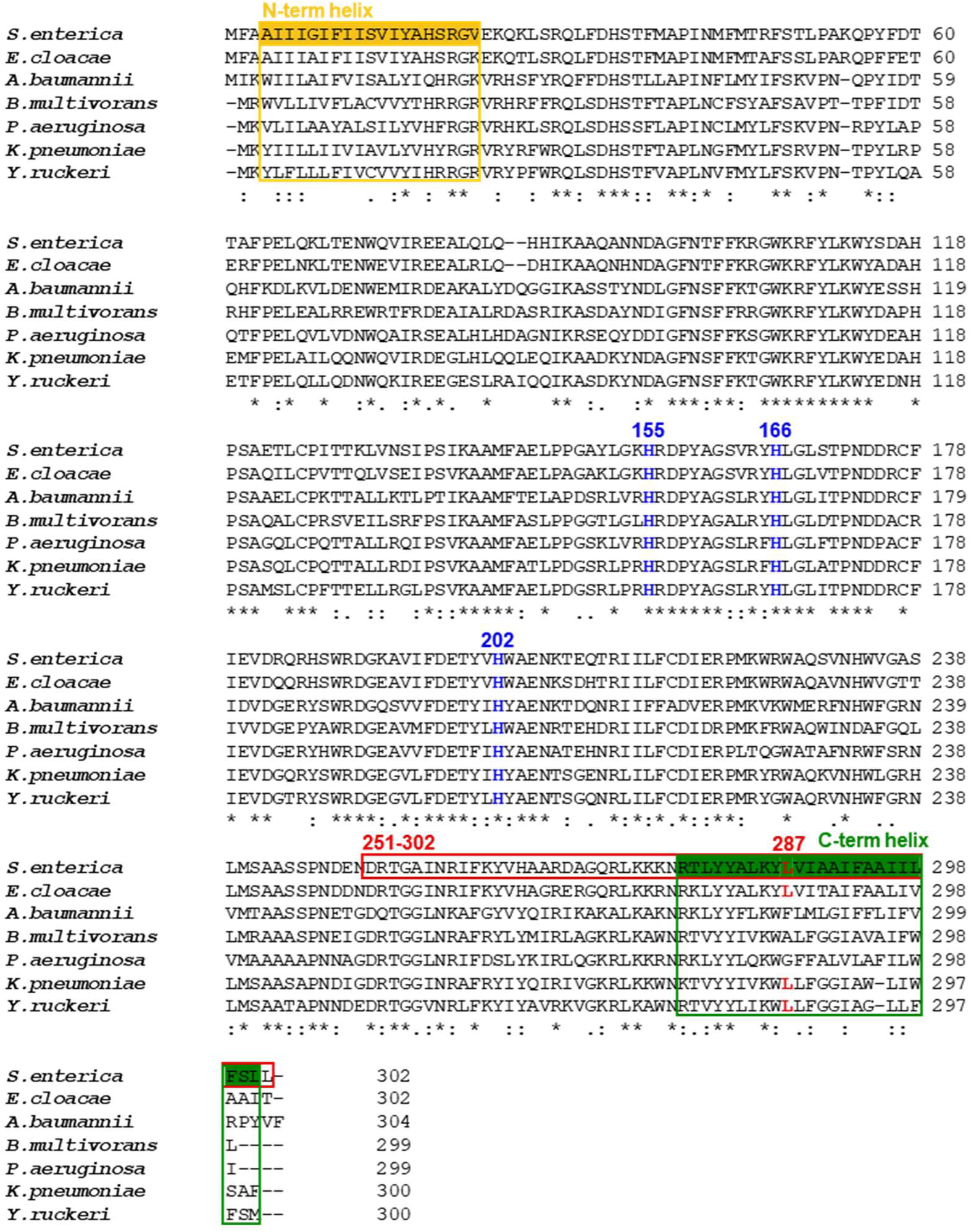
Multiple sequence alignment of LpxO proteins from representative Gram-negative bacteria that possess both LpxO and MgtC. Histidine residues (H155, H166, and H202; blue), previously reported (4) as essential for LpxO hydroxylase activity, are well conserved across species. The residue corresponding to Leu287 in *Salmonella* Typhimurium is occupied predominantly by leucine, with occasional substitutions to glycine or phenylalanine. Sequence alignment was performed using the CLUSTALW web tool (www.genome.jp/tools-bin/clustalw). *S. enterica*, *Salmonella enterica* serovar Typhimurium 14028s (GenBank; ACY91495.1), *E. cloacae*, *Enterobacter cloacae* (GenBank; ADF59904.1), *A. baumannii*, *Acinetobacter baumannii* (GenBank; EEX03680.1), *B. multivorans*, *Burkholderia multivorans* (GenBank; PRE55428.1), *P. aeruginosa*, *Pseudomonas aeruginosa* (GenBank; AAG07900.1), *K. pneumoniae*, *Klebsiella pneumoniae* (GenBank; AEW62766.1), *Y. ruckeri*, *Yersinia ruckeri* (GenBank; CEK28262.1). The originally identified fragment from BACTH screening is indicated by a red box and the C-terminal helix is indicated by a green box. The Leu287 residue is highlighted in red.

**Figure S3.**
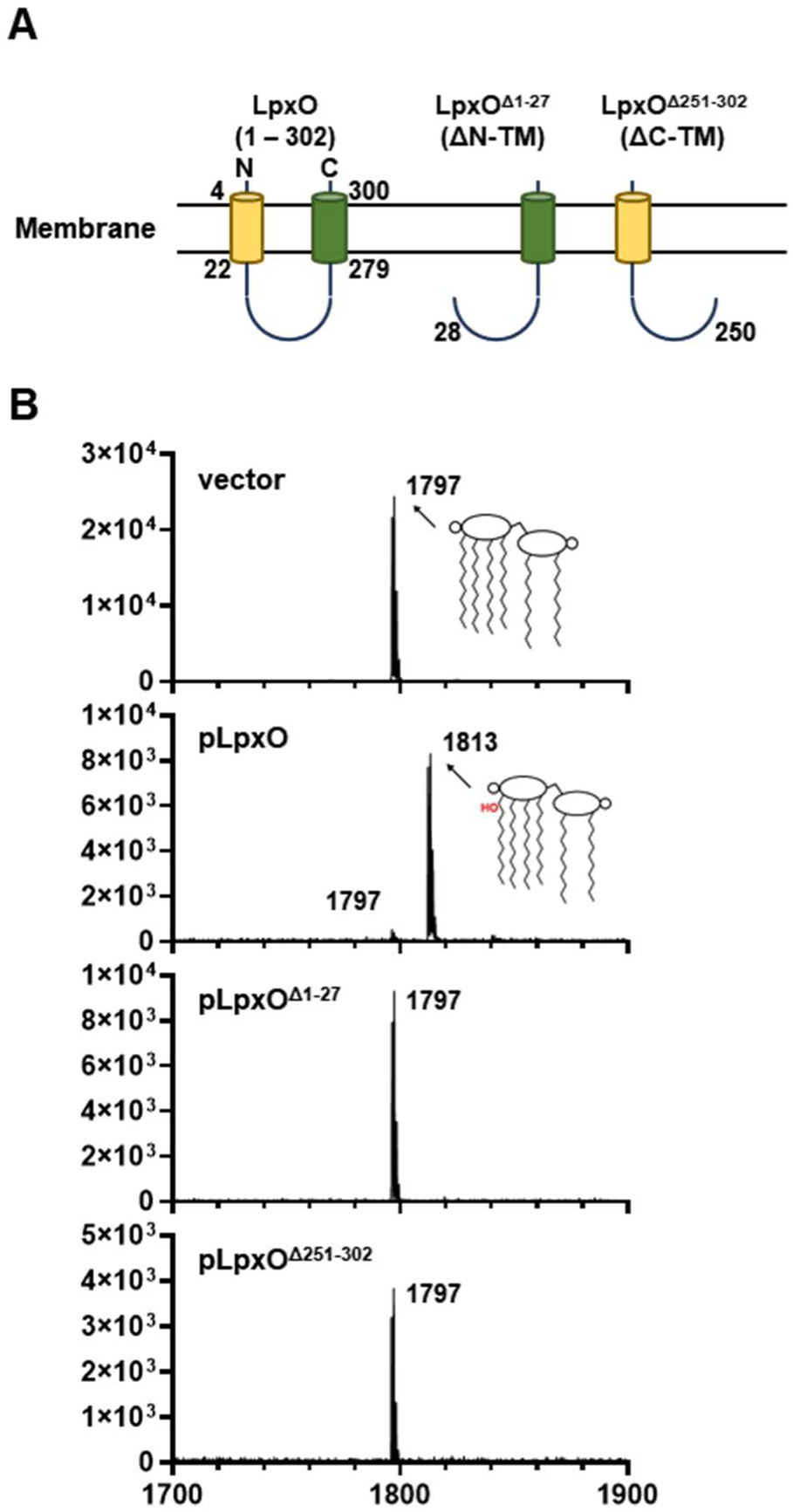
Both the N- and C-terminal helices of LpxO are required for lipid A hydroxylase activity. Lipid A profile under high Mg^2+^ conditions. Lipid A was extracted from Δ*lpxO Salmonella* strain carrying pBAD33 empty vector (vector), pLpxO^WT^, pLpxO^Δ1-27^ (N-terminal helix deletion), or pLpxO^Δ251-302^ (C-terminal helix deletion) and analyzed by MALDI-TOF MS. Strains were grown in N-minimal medium containing 10 mM Mg^2+^, and 1 mM arabinose was added to induce LpxO^WT^, LpxO^Δ1-27^, or LpxO^Δ251-302^ for 3 hours.

**Figure S4.**
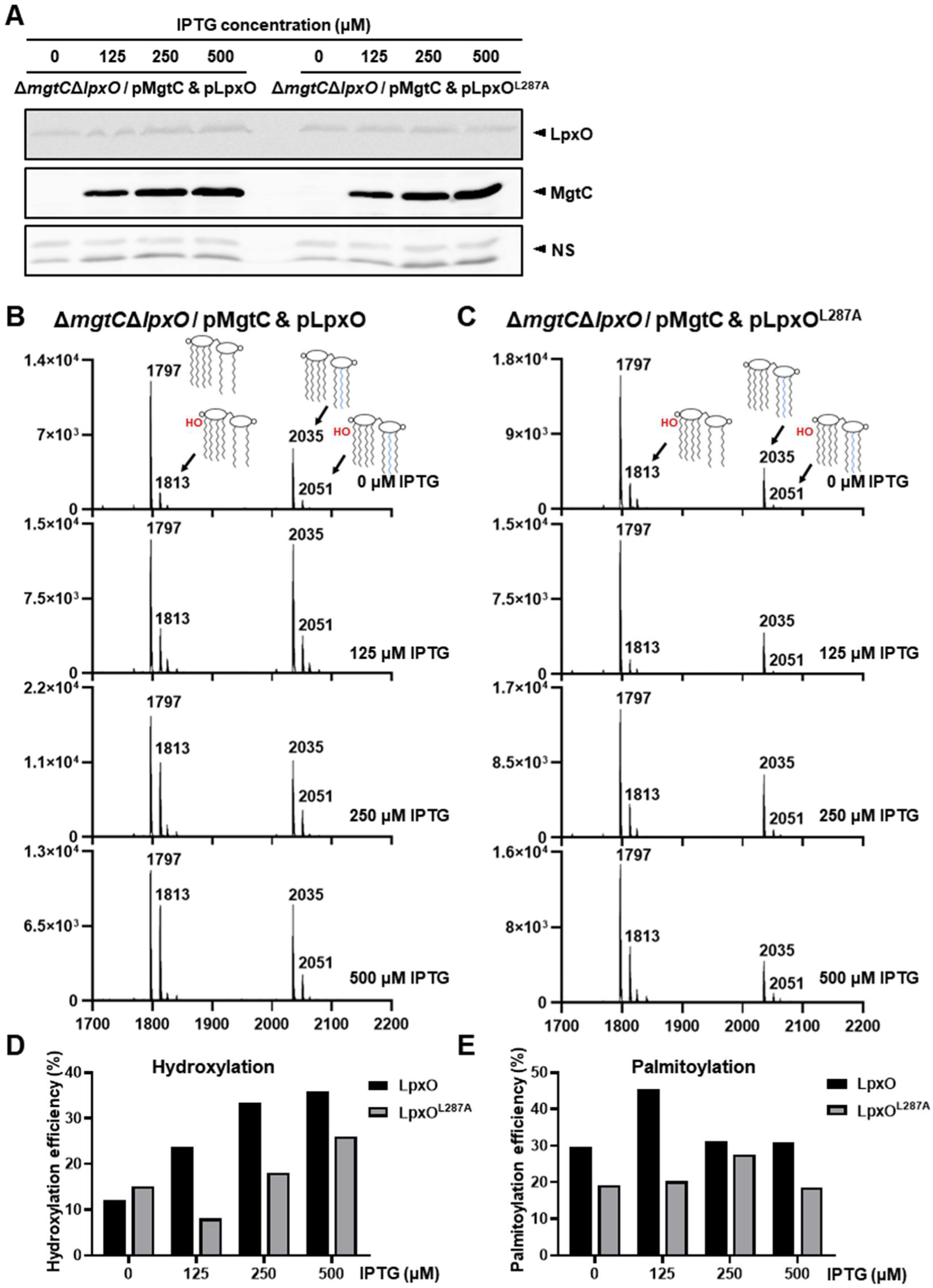
MgtC increases LpxO-mediated lipid A hydroxylation in a concentration-dependent manner even at low magnesium concentrations. (A) Western blot analysis of a strain carrying a Myc-tagged *lpxO*^WT^ or *lpxO*^L287A^ genes and an IPTG-inducible *mgtC* gene. LpxO was expressed at a basal level, and MgtC expression was induced with different IPTG concentrations. Nonspecific bands in the same blot were shown as a control. (B-C) The Δ*mgtC*Δ*lpxO* double knock-out mutant was co-transformed with plasmids allowing IPTG-inducible expression of MgtC and basal level expression of LpxO (B) or LpxO^L287A^ (C). Bacteria were grown in N-minimal medium containing 10 μM Mg^2+^ and incubated for 3 to 4 hours (OD_600_ = 0.1). After the addition of 0 to 500 μM IPTG to induce MgtC, incubation continued for 2 hours. Cells were then harvested for lipid A extraction and subsequent analysis. (D) Lipid A hydroxylation efficiency in response to MgtC expression. Hydroxylation efficiency (%) = [sum of intensities of hydroxylated lipid A/sum of intensities of unmodified lipid A]×100. (E) Lipid A palmitoylation efficiency in response to MgtC expression. Palmitoylation efficiency (%) = [sum of intensities of palmitoylated lipid A/ sum of intensities of unmodified lipid A]×100.

**Figure S5.**
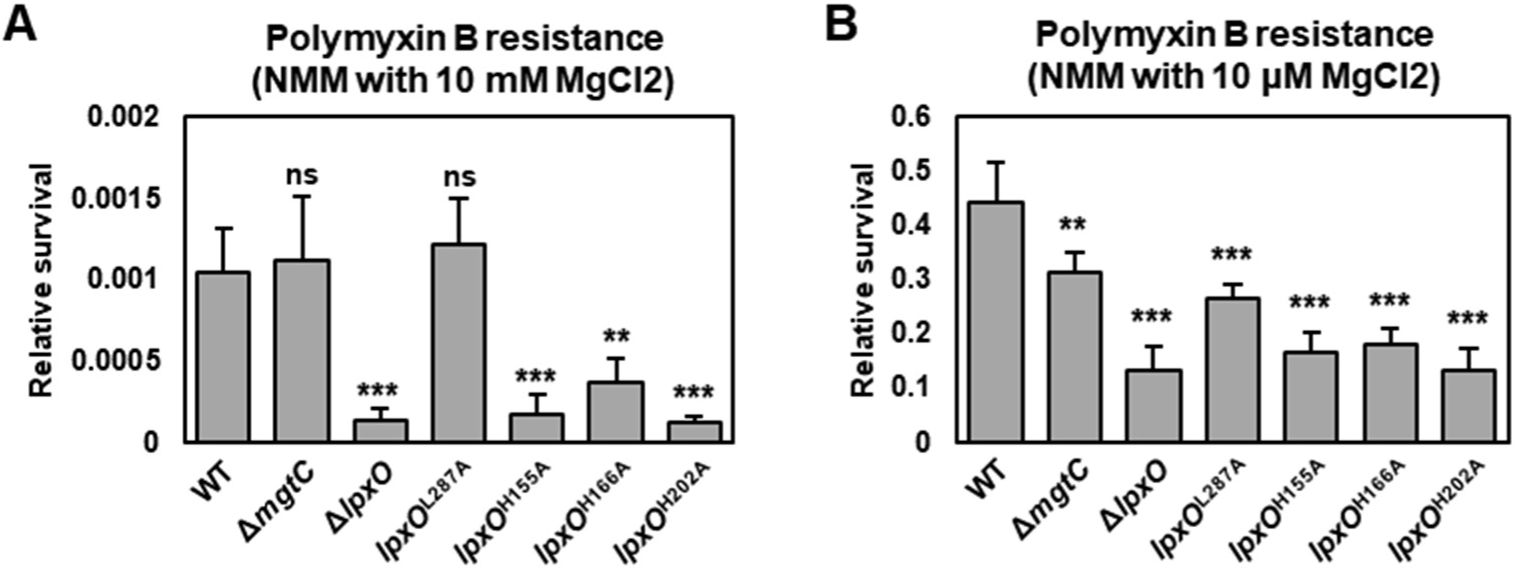
The Leu287-to-Ala substitution in LpxO decreases polymyxin B resistance in low Mg^2+^ but not in high Mg^2+^. (A-B) Polymyxin B killing assay under MgtC-repressing (A) (10 mM Mg^2+^) and MgtC-inducing (B) conditions (10 μM Mg^2+^) (5). Cells were treated with 2 μg/ml polymyxin B for 1 hour. Survival was calculated as the ratio of CFU in treated versus untreated samples. Data represent means ± SD of five independent experiments. Statistical analysis was performed using one-way ANOVA followed by Dunnett’s multiple comparisons test.

**Figure S6.**
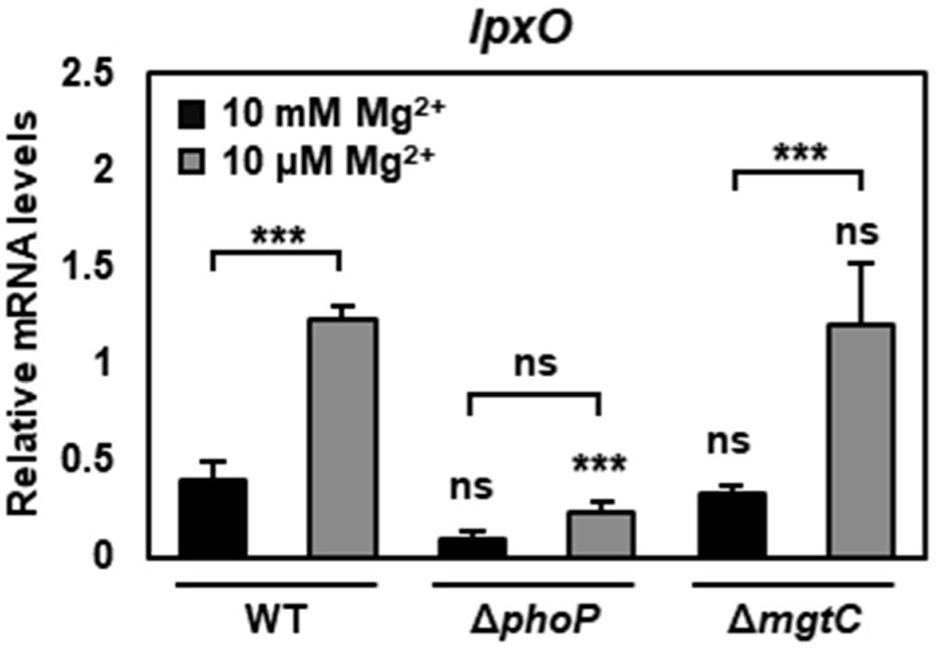
*lpxO* transcription increases in response to low Mg^2+^ in a PhoP-dependent manner. Quantitative real-time PCR (qRT-PCR) analysis of *lpxO* transcript levels in wild-type, Δ*mgtC* and Δ*phoP* cultured in NMM containing either 10 mM or 10 μM Mg^2+^. The mRNA levels were normalized to the 16S rRNA gene (*rrsH*) and are presented as means ± SD from three independent experiments.

**Figure S7.**
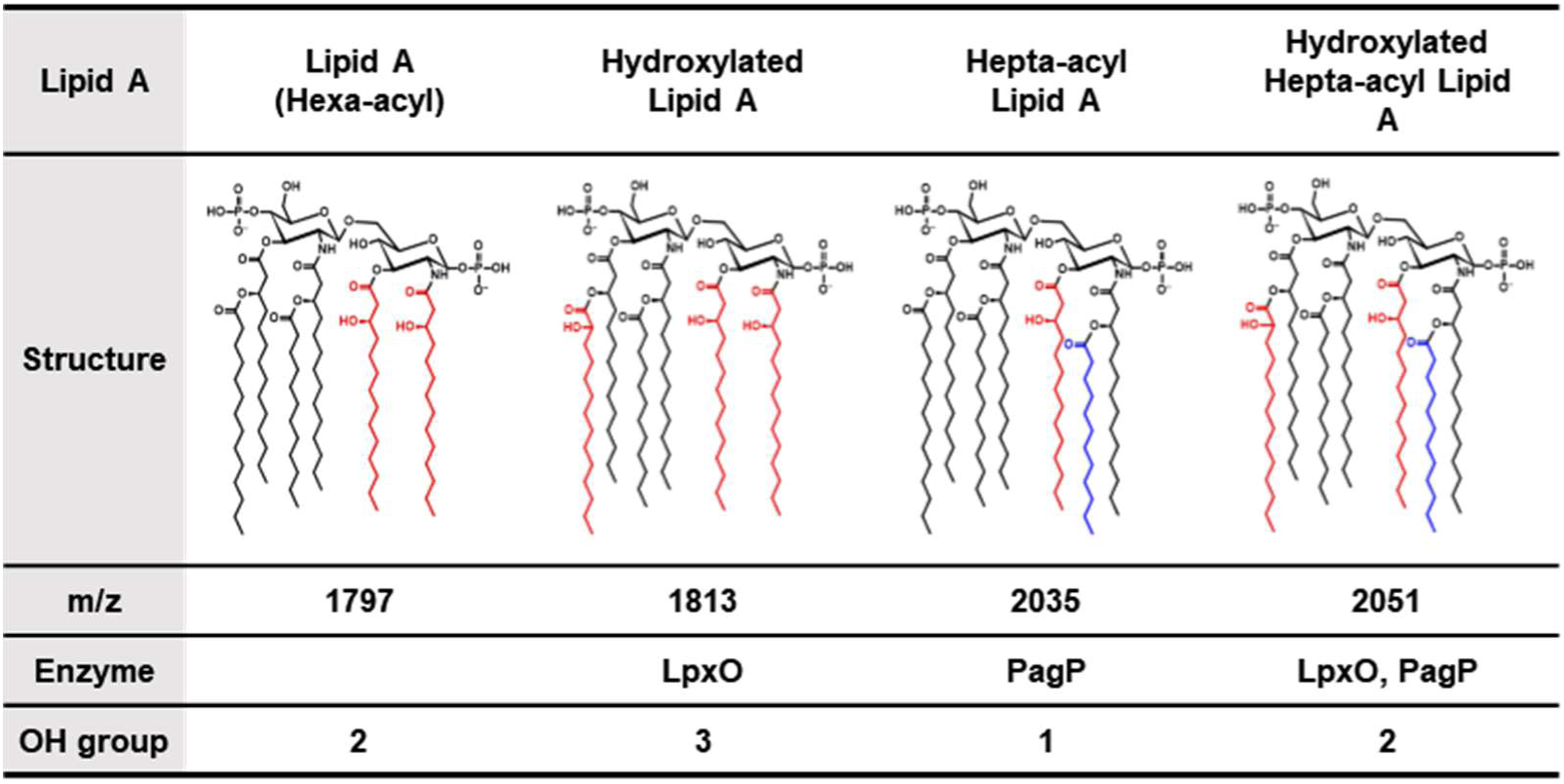
Number of the hydroxyl group in lipid A changes depending on LpxO and PagP. Unmodified lipid A contains 2 acyl chains with a hydroxyl group. When lipid A is hydroxylated by LpxO, the number of hydroxyl groups increases to 3. PagP adds a palmitoyl group to the hydroxyl group of the acyl chain, resulting in hepta-acyl lipid A having only 1 hydroxyl group. When further hydroxylated by LpxO, the number of hydroxyl groups returns to 2.

**Table S1.**
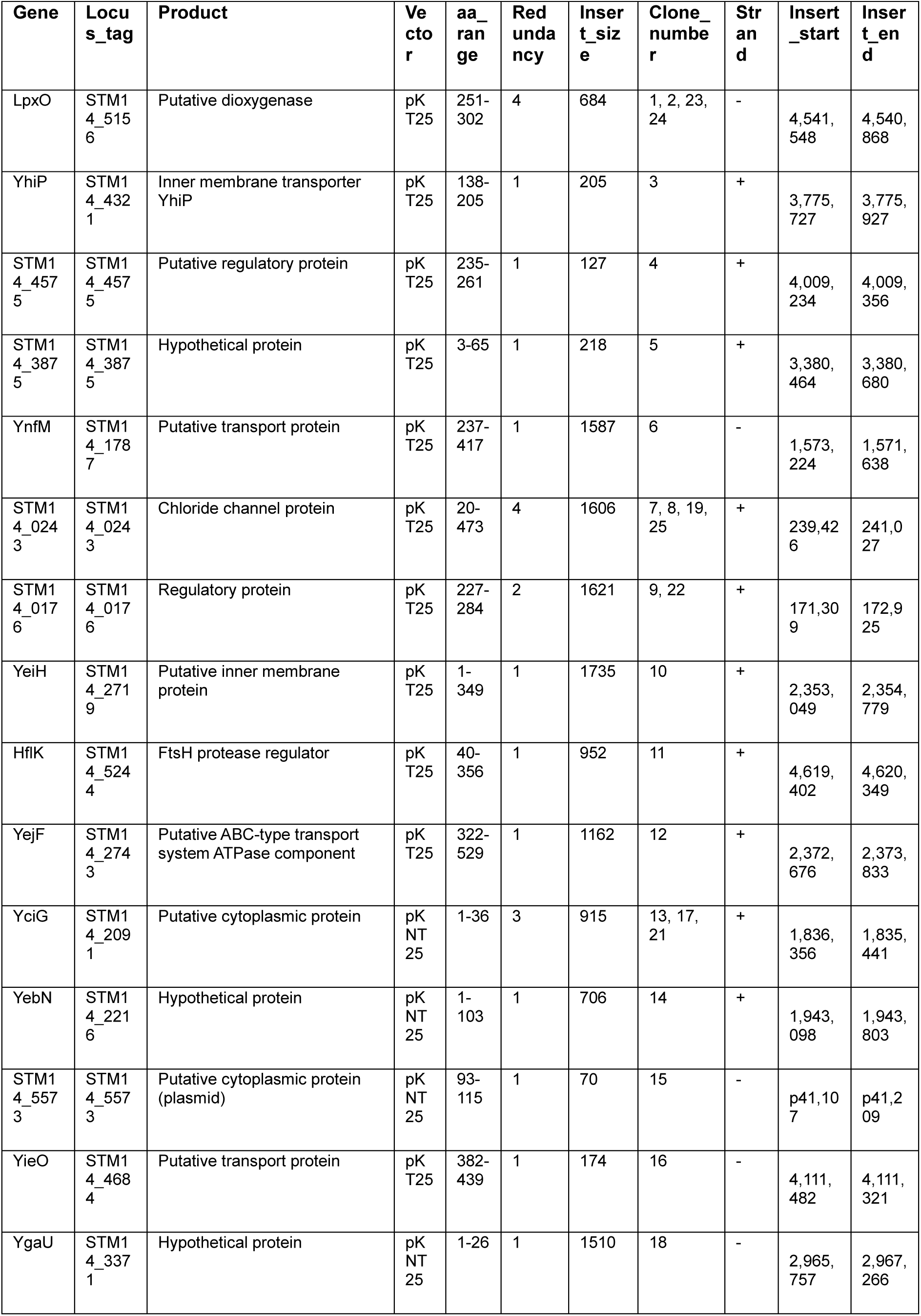
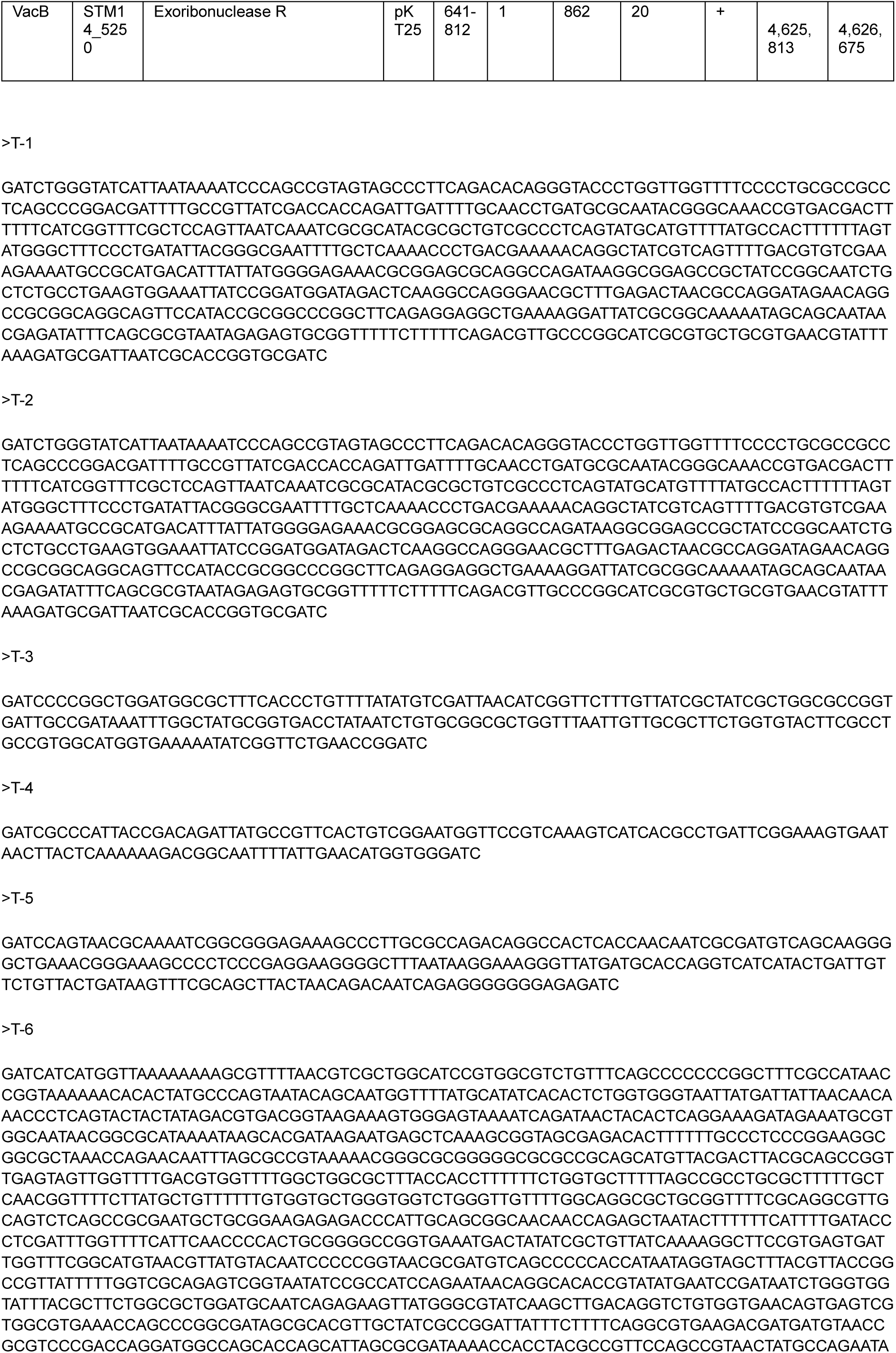
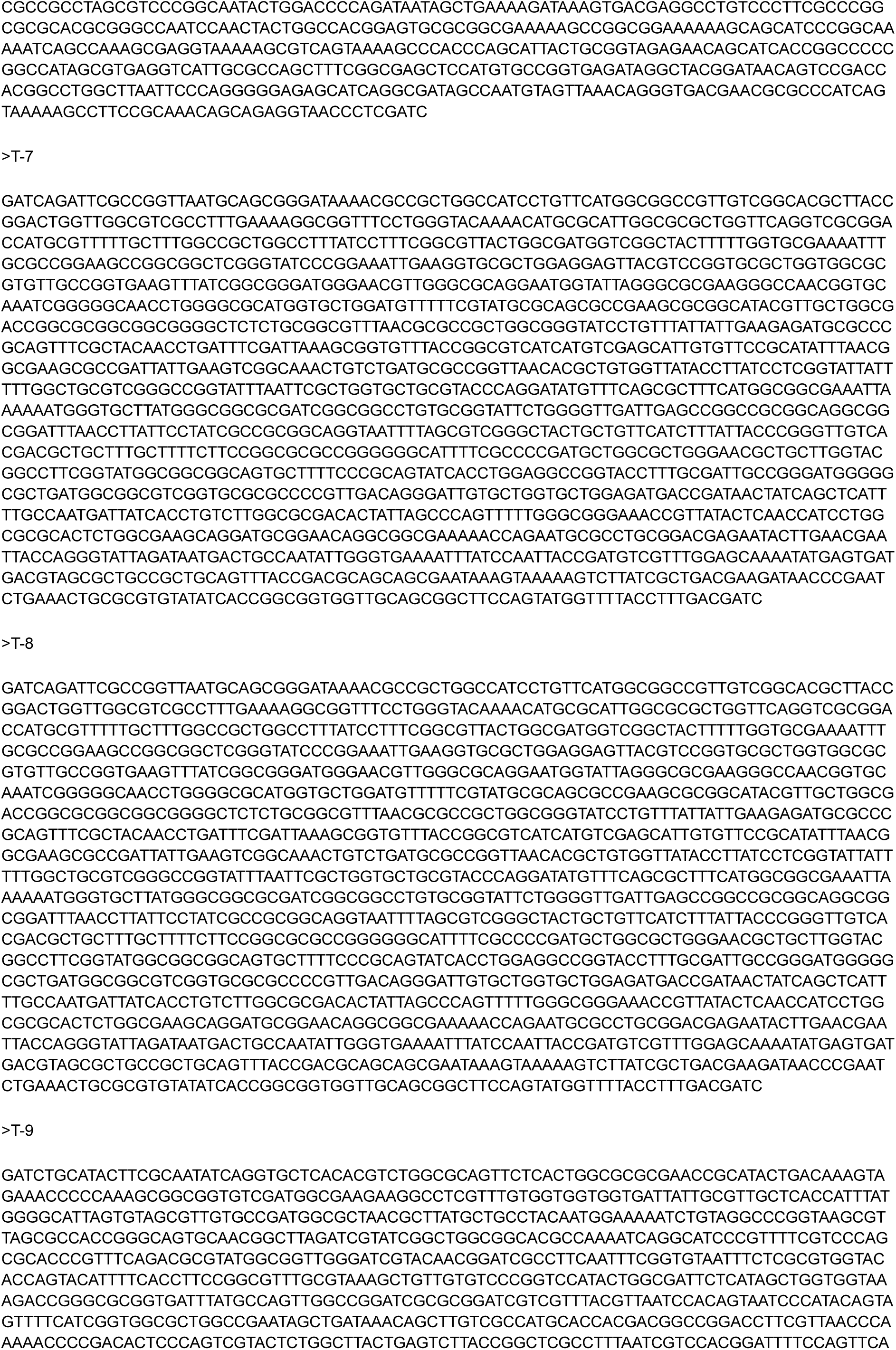
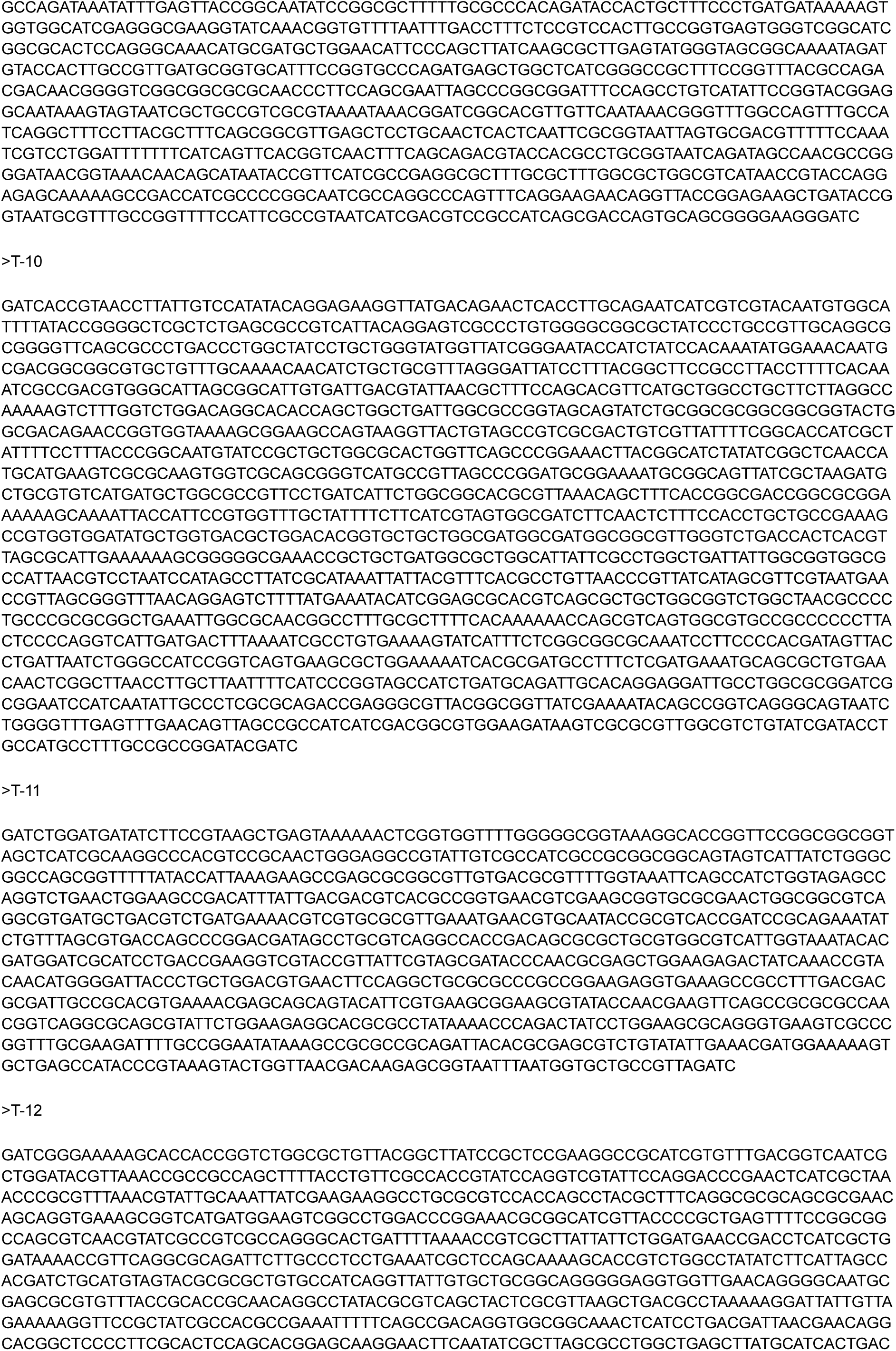
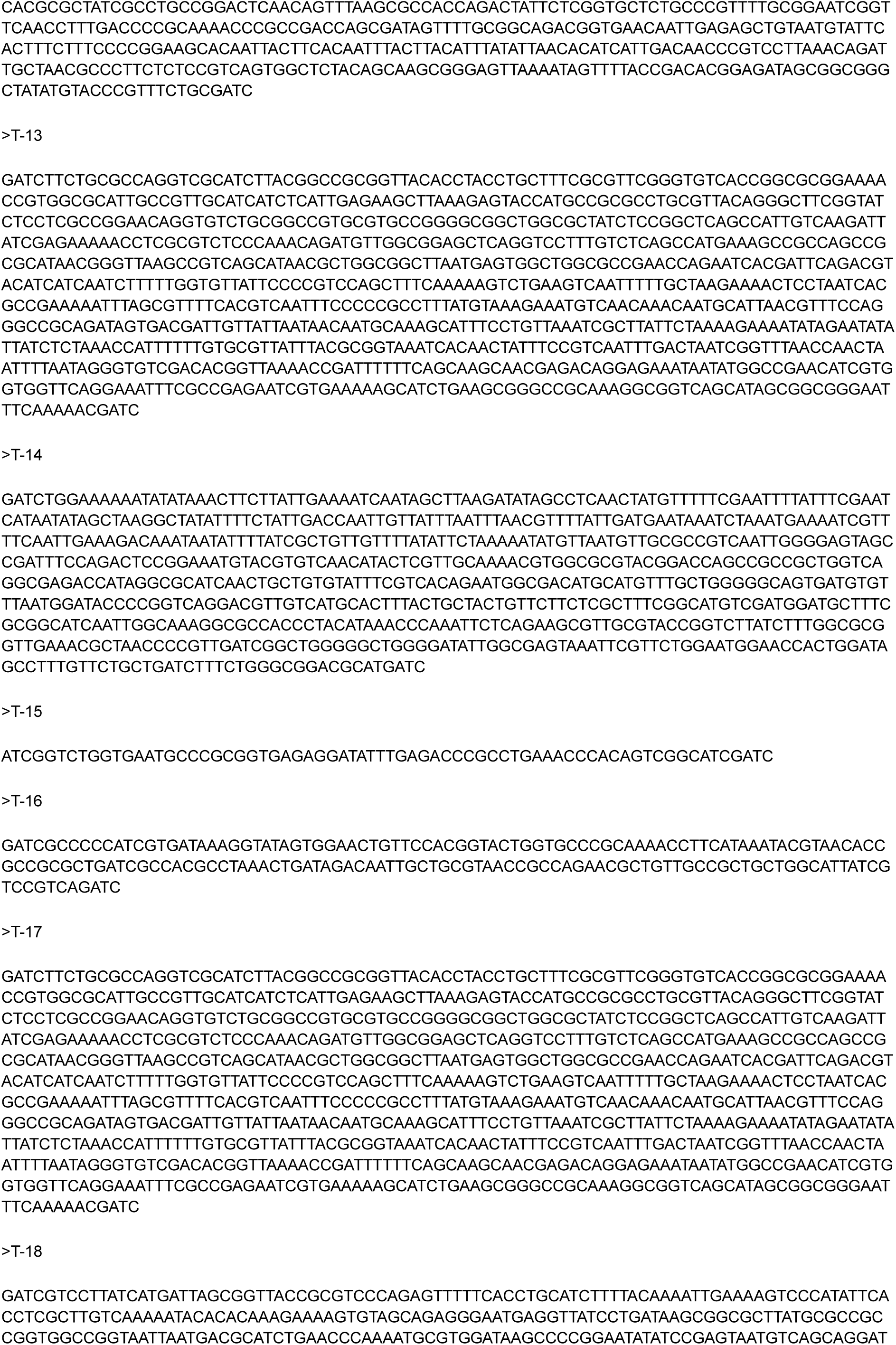
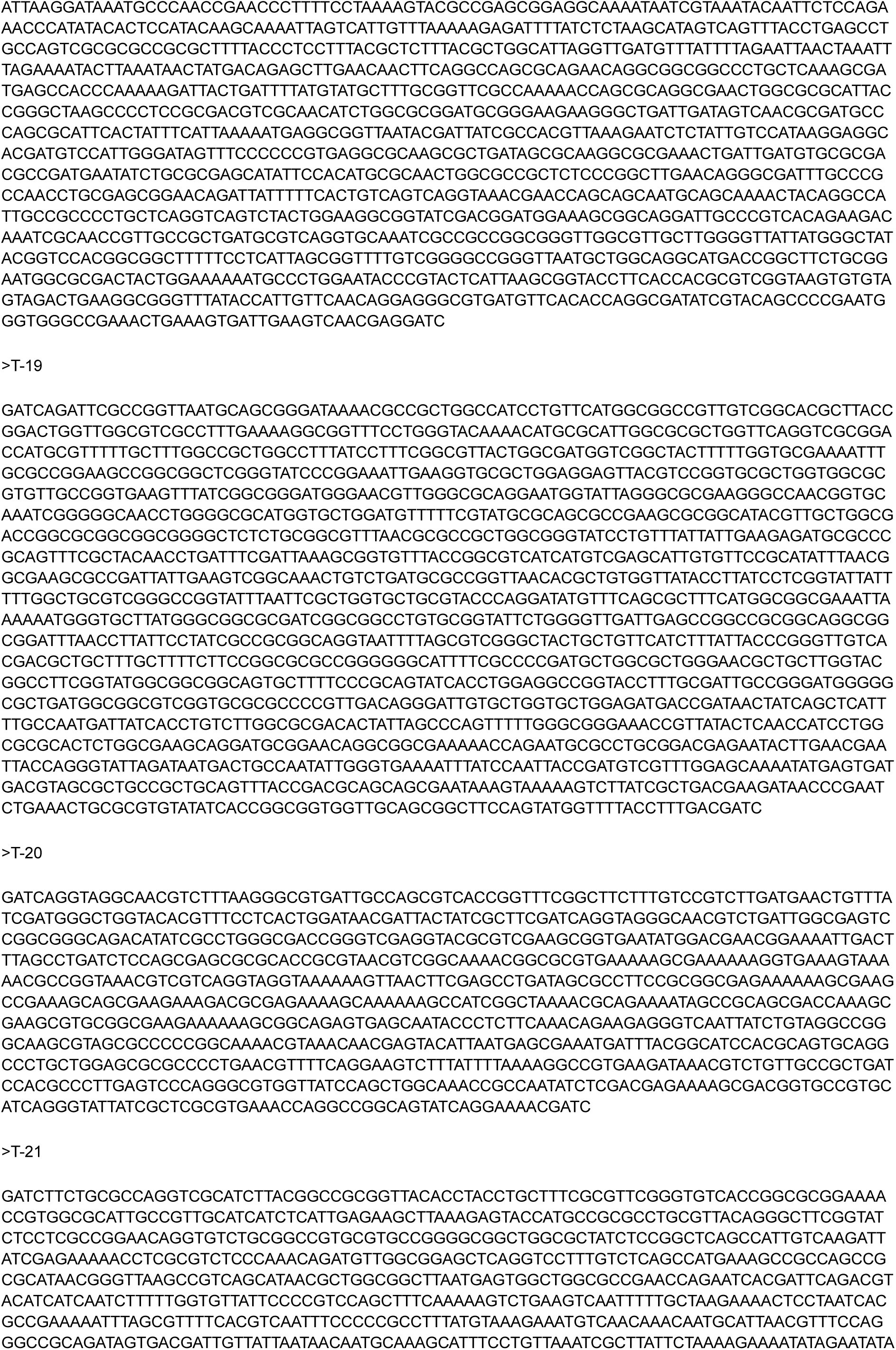
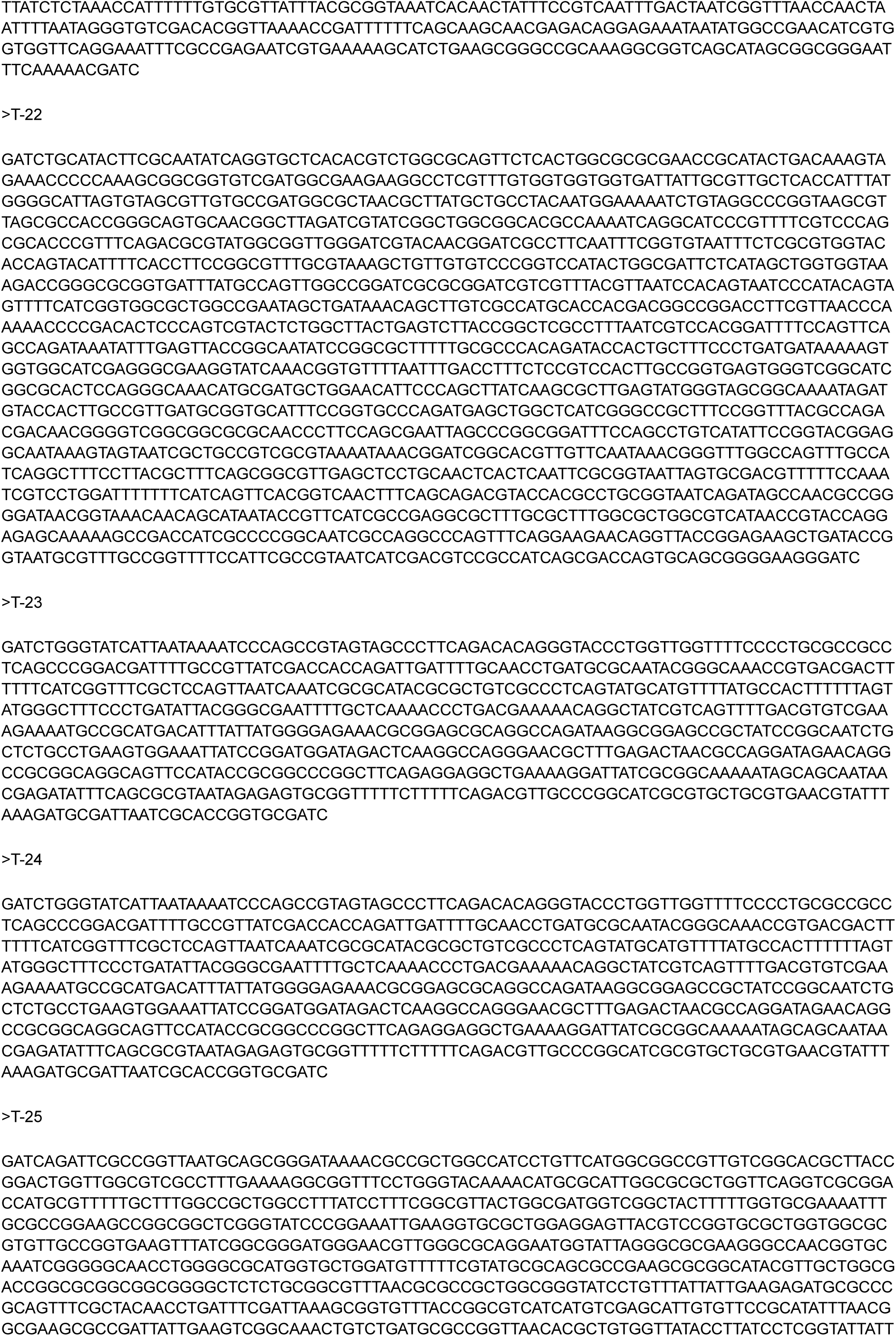
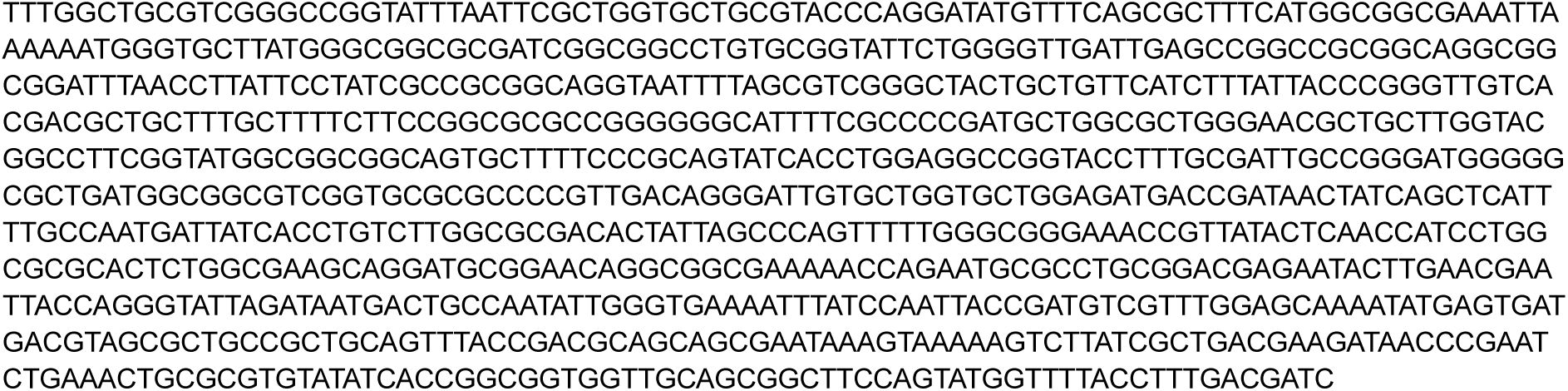
List of MgtC-interacting proteins identified by BACTH genomic library screening. MgtC-interacting proteins were identified through three independent BACTH library screening trials. The aa_range indicates the amino acid coordinates of the insert in frame with the T25 fragment. Redundancy indicates the number of independent clones recovered across three screening trials. DNA sequences of 25 identified clones are shown below.

| Gene | Locus_tag | Product | Vector | aa_range | Redundancy | Insert_size | Clone_number | Strand | Insert_start | Insert_end |
|---|---|---|---|---|---|---|---|---|---|---|
| LpxO | STM14_5156 | Putative dioxygenase | pKT25 | 251-302 | 4 | 684 | 1, 2, 23, 24 | - | 4,541,548 | 4,540,868 |
| YhiP | STM14_4321 | Inner membrane transporter YhiP | pKT25 | 138-205 | 1 | 205 | 3 | + | 3,775,727 | 3,775,927 |
| STM14_4575 | STM14_4575 | Putative regulatory protein | pKT25 | 235-261 | 1 | 127 | 4 | + | 4,009,234 | 4,009,356 |
| STM14_3875 | STM14_3875 | Hypothetical protein | pKT25 | 3-65 | 1 | 218 | 5 | + | 3,380,464 | 3,380,680 |
| YnfM | STM14_1787 | Putative transport protein | pKT25 | 237-417 | 1 | 1587 | 6 | - | 1,573,224 | 1,571,638 |
| STM14_0243 | STM14_0243 | Chloride channel protein | pKT25 | 20-473 | 4 | 1606 | 7, 8, 19, 25 | + | 239,426 | 241,027 |
| STM14_0176 | STM14_0176 | Regulatory protein | pKT25 | 227-284 | 2 | 1621 | 9, 22 | + | 171,309 | 172,925 |
| YeiH | STM14_2719 | Putative inner membrane protein | pKT25 | 1-349 | 1 | 1735 | 10 | + | 2,353,049 | 2,354,779 |
| HflK | STM14_5244 | FtsH protease regulator | pKT25 | 40-356 | 1 | 952 | 11 | + | 4,619,402 | 4,620,349 |
| YejF | STM14_2743 | Putative ABC-type transport system ATPase component | pKT25 | 322-529 | 1 | 1162 | 12 | + | 2,372,676 | 2,373,833 |
| YciG | STM14_2091 | Putative cytoplasmic protein | pKNT25 | 1-36 | 3 | 915 | 13, 17, 21 | + | 1,836,356 | 1,835,441 |
| YebN | STM14_2216 | Hypothetical protein | pKNT25 | 1-103 | 1 | 706 | 14 | + | 1,943,098 | 1,943,803 |
| STM14_5573 | STM14_5573 | Putative cytoplasmic protein (plasmid) | pKNT25 | 93-115 | 1 | 70 | 15 | - | p41,107 | p41,209 |
| YieO | STM14_4684 | Putative transport protein | pKT25 | 382-439 | 1 | 174 | 16 | - | 4,111,482 | 4,111,321 |
| YgaU | STM14_3371 | Hypothetical protein | pKNT25 | 1-26 | 1 | 1510 | 18 | - | 2,965,757 | 2,967,266 |
| VacB | STM14_5250 | Exoribonuclease R | pKT25 | 641-812 | 1 | 862 | 20 | + | 4,625,813 | 4,626,675 |

>T-1
GATCTGGGTATCATTAATAAAATCCCAGCCGTAGTAGCCCTTCAGACACAGGGTACCCTGGTTGGTTTTCCCCTGCGCCGCCTCAGCCCGGACGATTTTGCCGTTATCGACCACCAGATTGATTTTGCAACCTGATGCGCAATACGGGCAAACCGTGACGACTTTTTTCATCGGTTTCGCTCCAGTTAATCAAATCGCGCATACGCGCTGTCGCCCTCAGTATGCATGTTTTATGCCACTTTTTTAGTATGGGCTTTCCCTGATATTACGGGCGAATTTTGCTCAAAACCCTGACGAAAAACAGGCTATCGTCAGTTTTGACGTGTCGAAAGAAAATGCCGCATGACATTTATTATGGGGAGAAACGCGGAGCGCAGGCCAGATAAGGCGGAGCCGCTATCCGGCAATCTGCTCTGCCTGAAGTGGAAATTATCCGGATGGATAGACTCAAGGCCAGGGAACGCTTTGAGACTAACGCCAGGATAGAACAGGCCGCGGCAGGCAGTTCCATACCGCGGCCCGGCTTCAGAGGAGGCTGAAAAGGATTATCGCGGCAAAAATAGCAGCAATAACGAGATATTTCAGCGCGTAATAGAGAGTGCGGTTTTTCTTTTTCAGACGTTGCCCGGCATCGCGTGCTGCGTGAACGTATTTAAAGATGCGATTAATCGCACCGGTGCGATC
>T-2
GATCTGGGTATCATTAATAAAATCCCAGCCGTAGTAGCCCTTCAGACACAGGGTACCCTGGTTGGTTTTCCCCTGCGCCGCCTCAGCCCGGACGATTTTGCCGTTATCGACCACCAGATTGATTTTGCAACCTGATGCGCAATACGGGCAAACCGTGACGACTTTTTTCATCGGTTTCGCTCCAGTTAATCAAATCGCGCATACGCGCTGTCGCCCTCAGTATGCATGTTTTATGCCACTTTTTTAGTATGGGCTTTCCCTGATATTACGGGCGAATTTTGCTCAAAACCCTGACGAAAAACAGGCTATCGTCAGTTTTGACGTGTCGAAAGAAAATGCCGCATGACATTTATTATGGGGAGAAACGCGGAGCGCAGGCCAGATAAGGCGGAGCCGCTATCCGGCAATCTGCTCTGCCTGAAGTGGAAATTATCCGGATGGATAGACTCAAGGCCAGGGAACGCTTTGAGACTAACGCCAGGATAGAACAGGCCGCGGCAGGCAGTTCCATACCGCGGCCCGGCTTCAGAGGAGGCTGAAAAGGATTATCGCGGCAAAAATAGCAGCAATAACGAGATATTTCAGCGCGTAATAGAGAGTGCGGTTTTTCTTTTTCAGACGTTGCCCGGCATCGCGTGCTGCGTGAACGTATTTAAAGATGCGATTAATCGCACCGGTGCGATC
>T-3
GATCCCCGGCTGGATGGCGCTTTCACCCTGTTTTATATGTCGATTAACATCGGTTCTTTGTTATCGCTATCGCTGGCGCCGGTGATTGCCGATAAATTTGGCTATGCGGTGACCTATAATCTGTGCGGCGCTGGTTTAATTGTTGCGCTTCTGGTGTACTTCGCCTGCCGTGGCATGGTGAAAAATATCGGTTCTGAACCGGATC
>T-4
GATCGCCCATTACCGACAGATTATGCCGTTCACTGTCGGAATGGTTCCGTCAAAGTCATCACGCCTGATTCGGAAAGTGAATAACTTACTCAAAAAAGACGGCAATTTTATTGAACATGGTGGGATC
>T-5
GATCCAGTAACGCAAAATCGGCGGGAGAAAGCCCTTGCGCCAGACAGGCCACTCACCAACAATCGCGATGTCAGCAAGGGGCTGAAACGGGAAAGCCCCTCCCGAGGAAGGGGCTTTAATAAGGAAAGGGTTATGATGCACCAGGTCATCATACTGATTGTTCTGTTACTGATAAGTTTCGCAGCTTACTAACAGACAATCAGAGGGGGGGAGAGATC
>T-6
GATCATCATGGTTAAAAAAAAGCGTTTTAACGTCGCTGGCATCCGTGGCGTCTGTTTCAGCCCCCCGGCTTTCGCCATAACCGGTAAAAAACACACTATGCCCAGTAATACAGCAATGGTTTTATGCATATCACACTCTGGTGGGTAATTATGATTATTAACAACAAACCCTCAGTACTACTATAGACGTGACGGTAAGAAAGTGGGAGTAAAATCAGATAACTACACTCAGGAAAGATAGAAATGCGTGGCAATAACGGCGCATAAAATAAGCACGATAAGAATGAGCTCAAAGCGGTAGCGAGACACTTTTTTGCCCTCCCGGAAGGCGGCGCTAAACCAGAACAATTTAGCGCCGTAAAAACGGGCGCGGGGGCGCGCCGCAGCATGTTACGACTTACGCAGCCGGTTGAGTAGTTGGTTTTGACGTGGTTTTGGCTGGCGCTTTACCACCTTTTTTCTGGTGCTTTTTAGCCGCCTGCGCTTTTTGCTCAACGGTTTTCTTATGCTGTTTTTTGTGGTGCTGGGTGGTCTGGGTTGTTTTGGCAGGCGCTGCGGTTTTCGCAGGCGTTGCAGTCTCAGCCGCGAATGCTGCGGAAGAGAGACCCATTGCAGCGGCAACAACCAGAGCTAATACTTTTTTCATTTTGATACCCTCGATTTGGTTTTCATTCAACCCCACTGCGGGGCCGGTGAAATGACTATATCGCTGTTATCAAAAGGCTTCCGTGAGTGATTGGTTTCGGCATGTAACGTTATGTACAATCCCCCGGTAACGCGATGTCAGCCCCCACCATAATAGGTAGCTTTACGTTACCGGCCGTTATTTTTTGGTCGCAGAGTCGGTAATATCCGCCATCCAGAATAACAGGCACACCGTATATGAATCCGATAATCTGGGTGGTATTTACGCTTCTGGCGCTGGATGCAATCAGAGAAGTTATGGGCGTATCAAGCTTGACAGGTCTGTGGTGAACAGTGAGTCGTGGCGTGAAACCAGCCCGGCGATAGCGCACGTTGCTATCGCCGGATTATTTCTTTTCAGGCGTGAAGACGATGATGTAACCGCGTCCCGACCAGGATGGCCAGCACCAGCATTAGCGCGATAAAACCACCTACGCCGTTCCAGCCGTAACTATGCCAGAATACGCCGCCTAGCGTCCCGGCAATACTGGACCCCAGATAATAGCTGAAAAGATAAAGTGACGAGGCCTGTCCCTTCGCCCGGCGCGCACGCGGGCCAATCCAACTACTGGCCACGGAGTGCGCGGCGAAAAAGCCGGCGGAAAAAAGCAGCATCCCGGCAAAAATCAGCCAAAGCGAGGTAAAAAGCGTCAGTAAAAGCCCACCCAGCATTACTGCGGTAGAGAACAGCATCACCGGCCCCCGGCCATAGCGTGAGGTCATTGCGCCAGCTTTCGGCGAGCTCCATGTGCCGGTGAGATAGGCTACGGATAACAGTCCGACCACGGCCTGGCTTAATTCCCAGGGGGAGAGCATCAGGCGATAGCCAATGTAGTTAAACAGGGTGACGAACGCGCCCATCAGTAAAAAGCCTTCCGCAAACAGCAGAGGTAACCCTCGATC
>T-7
GATCAGATTCGCCGGTTAATGCAGCGGGATAAAACGCCGCTGGCCATCCTGTTCATGGCGGCCGTTGTCGGCACGCTTACCGGACTGGTTGGCGTCGCCTTTGAAAAGGCGGTTTCCTGGGTACAAAACATGCGCATTGGCGCGCTGGTTCAGGTCGCGGACCATGCGTTTTTGCTTTGGCCGCTGGCCTTTATCCTTTCGGCGTTACTGGCGATGGTCGGCTACTTTTTGGTGCGAAAATTTGCGCCGGAAGCCGGCGGCTCGGGTATCCCGGAAATTGAAGGTGCGCTGGAGGAGTTACGTCCGGTGCGCTGGTGGCGCGTGTTGCCGGTGAAGTTTATCGGCGGGATGGGAACGTTGGGCGCAGGAATGGTATTAGGGCGCGAAGGGCCAACGGTGCAAATCGGGGGCAACCTGGGGCGCATGGTGCTGGATGTTTTTCGTATGCGCAGCGCCGAAGCGCGGCATACGTTGCTGGCGACCGGCGCGGCGGCGGGGCTCTCTGCGGCGTTTAACGCGCCGCTGGCGGGTATCCTGTTTATTATTGAAGAGATGCGCCCGCAGTTTCGCTACAACCTGATTTCGATTAAAGCGGTGTTTACCGGCGTCATCATGTCGAGCATTGTGTTCCGCATATTTAACGGCGAAGCGCCGATTATTGAAGTCGGCAAACTGTCTGATGCGCCGGTTAACACGCTGTGGTTATACCTTATCCTCGGTATTATTTTTGGCTGCGTCGGGCCGGTATTTAATTCGCTGGTGCTGCGTACCCAGGATATGTTTCAGCGCTTTCATGGCGGCGAAATTAAAAAATGGGTGCTTATGGGCGGCGCGATCGGCGGCCTGTGCGGTATTCTGGGGTTGATTGAGCCGGCCGCGGCAGGCGGCGGATTTAACCTTATTCCTATCGCCGCGGCAGGTAATTTTAGCGTCGGGCTACTGCTGTTCATCTTTATTACCCGGGTTGTCACGACGCTGCTTTGCTTTTCTTCCGGCGCGCCGGGGGGCATTTTCGCCCCGATGCTGGCGCTGGGAACGCTGCTTGGTACGGCCTTCGGTATGGCGGCGGCAGTGCTTTTCCCGCAGTATCACCTGGAGGCCGGTACCTTTGCGATTGCCGGGATGGGGGCGCTGATGGCGGCGTCGGTGCGCGCCCCGTTGACAGGGATTGTGCTGGTGCTGGAGATGACCGATAACTATCAGCTCATTTTGCCAATGATTATCACCTGTCTTGGCGCGACACTATTAGCCCAGTTTTTGGGCGGGAAACCGTTATACTCAACCATCCTGGCGCGCACTCTGGCGAAGCAGGATGCGGAACAGGCGGCGAAAAACCAGAATGCGCCTGCGGACGAGAATACTTGAACGAATTACCAGGGTATTAGATAATGACTGCCAATATTGGGTGAAAATTTATCCAATTACCGATGTCGTTTGGAGCAAAATATGAGTGATGACGTAGCGCTGCCGCTGCAGTTTACCGACGCAGCAGCGAATAAAGTAAAAAGTCTTATCGCTGACGAAGATAACCCGAATCTGAAACTGCGCGTGTATATCACCGGCGGTGGTTGCAGCGGCTTCCAGTATGGTTTTACCTTTGACGATC
>T-8
GATCAGATTCGCCGGTTAATGCAGCGGGATAAAACGCCGCTGGCCATCCTGTTCATGGCGGCCGTTGTCGGCACGCTTACCGGACTGGTTGGCGTCGCCTTTGAAAAGGCGGTTTCCTGGGTACAAAACATGCGCATTGGCGCGCTGGTTCAGGTCGCGGACCATGCGTTTTTGCTTTGGCCGCTGGCCTTTATCCTTTCGGCGTTACTGGCGATGGTCGGCTACTTTTTGGTGCGAAAATTTGCGCCGGAAGCCGGCGGCTCGGGTATCCCGGAAATTGAAGGTGCGCTGGAGGAGTTACGTCCGGTGCGCTGGTGGCGCGTGTTGCCGGTGAAGTTTATCGGCGGGATGGGAACGTTGGGCGCAGGAATGGTATTAGGGCGCGAAGGGCCAACGGTGCAAATCGGGGGCAACCTGGGGCGCATGGTGCTGGATGTTTTTCGTATGCGCAGCGCCGAAGCGCGGCATACGTTGCTGGCGACCGGCGCGGCGGCGGGGCTCTCTGCGGCGTTTAACGCGCCGCTGGCGGGTATCCTGTTTATTATTGAAGAGATGCGCCCGCAGTTTCGCTACAACCTGATTTCGATTAAAGCGGTGTTTACCGGCGTCATCATGTCGAGCATTGTGTTCCGCATATTTAACGGCGAAGCGCCGATTATTGAAGTCGGCAAACTGTCTGATGCGCCGGTTAACACGCTGTGGTTATACCTTATCCTCGGTATTATTTTTGGCTGCGTCGGGCCGGTATTTAATTCGCTGGTGCTGCGTACCCAGGATATGTTTCAGCGCTTTCATGGCGGCGAAATTAAAAAATGGGTGCTTATGGGCGGCGCGATCGGCGGCCTGTGCGGTATTCTGGGGTTGATTGAGCCGGCCGCGGCAGGCGGCGGATTTAACCTTATTCCTATCGCCGCGGCAGGTAATTTTAGCGTCGGGCTACTGCTGTTCATCTTTATTACCCGGGTTGTCACGACGCTGCTTTGCTTTTCTTCCGGCGCGCCGGGGGGCATTTTCGCCCCGATGCTGGCGCTGGGAACGCTGCTTGGTACGGCCTTCGGTATGGCGGCGGCAGTGCTTTTCCCGCAGTATCACCTGGAGGCCGGTACCTTTGCGATTGCCGGGATGGGGGCGCTGATGGCGGCGTCGGTGCGCGCCCCGTTGACAGGGATTGTGCTGGTGCTGGAGATGACCGATAACTATCAGCTCATTTTGCCAATGATTATCACCTGTCTTGGCGCGACACTATTAGCCCAGTTTTTGGGCGGGAAACCGTTATACTCAACCATCCTGGCGCGCACTCTGGCGAAGCAGGATGCGGAACAGGCGGCGAAAAACCAGAATGCGCCTGCGGACGAGAATACTTGAACGAATTACCAGGGTATTAGATAATGACTGCCAATATTGGGTGAAAATTTATCCAATTACCGATGTCGTTTGGAGCAAAATATGAGTGATGACGTAGCGCTGCCGCTGCAGTTTACCGACGCAGCAGCGAATAAAGTAAAAAGTCTTATCGCTGACGAAGATAACCCGAATCTGAAACTGCGCGTGTATATCACCGGCGGTGGTTGCAGCGGCTTCCAGTATGGTTTTACCTTTGACGATC
>T-9
GATCTGCATACTTCGCAATATCAGGTGCTCACACGTCTGGCGCAGTTCTCACTGGCGCGCGAACCGCATACTGACAAAGTAGAAACCCCCAAAGCGGCGGTGTCGATGGCGAAGAAGGCCTCGTTTGTGGTGGTGGTGATTATTGCGTTGCTCACCATTTATGGGGCATTAGTGTAGCGTTGTGCCGATGGCGCTAACGCTTATGCTGCCTACAATGGAAAAATCTGTAGGCCCGGTAAGCGTTAGCGCCACCGGGCAGTGCAACGGCTTAGATCGTATCGGCTGGCGGCACGCCAAAATCAGGCATCCCGTTTTCGTCCCAGCGCACCCGTTTCAGACGCGTATGGCGGTTGGGATCGTACAACGGATCGCCTTCAATTTCGGTGTAATTTCTCGCGTGGTACACCAGTACATTTTCACCTTCCGGCGTTTGCGTAAAGCTGTTGTGTCCCGGTCCATACTGGCGATTCTCATAGCTGGTGGTAAAGACCGGGCGCGGTGATTTATGCCAGTTGGCCGGATCGCGCGGATCGTCGTTTACGTTAATCCACAGTAATCCCATACAGTAGTTTTCATCGGTGGCGCTGGCCGAATAGCTGATAAACAGCTTGTCGCCATGCACCACGACGGCCGGACCTTCGTTAACCCAAAAACCCCGACACTCCCAGTCGTACTCTGGCTTACTGAGTCTTACCGGCTCGCCTTTAATCGTCCACGGATTTTCCAGTTCAGCCAGATAAATATTTGAGTTACCGGCAATATCCGGCGCTTTTTGCGCCCACAGATACCACTGCTTTCCCTGATGATAAAAAGTGGTGGCATCGAGGGCGAAGGTATCAAACGGTGTTTTAATTTGACCTTTCTCCGTCCACTTGCCGGTGAGTGGGTCGGCATCGGCGCACTCCAGGGCAAACATGCGATGCTGGAACATTCCCAGCTTATCAAGCGCTTGAGTATGGGTAGCGGCAAAATAGATGTACCACTTGCCGTTGATGCGGTGCATTTCCGGTGCCCAGATGAGCTGGCTCATCGGGCCGCTTTCCGGTTTACGCCAGACGACAACGGGGTCGGCGGCGCGCAACCCTTCCAGCGAATTAGCCCGGCGGATTTCCAGCCTGTCATATTCCGGTACGGAGGCAATAAAGTAGTAATCGCTGCCGTCGCGTAAAATAAACGGATCGGCACGTTGTTCAATAAACGGGTTTGGCCAGTTTGCCATCAGGCTTTCCTTACGCTTTCAGCGGCGTTGAGCTCCTGCAACTCACTCAATTCGCGGTAATTAGTGCGACGTTTTTCCAAATCGTCCTGGATTTTTTTCATCAGTTCACGGTCAACTTTCAGCAGACGTACCACGCCTGCGGTAATCAGATAGCCAACGCCGGGGATAACGGTAAACAACAGCATAATACCGTTCATCGCCGAGGCGCTTTGCGCTTTGGCGCTGGCGTCATAACCGTACCAGGAGAGCAAAAAGCCGACCATCGCCCCGGCAATCGCCAGGCCCAGTTTCAGGAAGAACAGGTTACCGGAGAAGCTGATACCGGTAATGCGTTTGCCGGTTTTCCATTCGCCGTAATCATCGACGTCCGCCATCAGCGACCAGTGCAGCGGGGAAGGGATC
>T-10
GATCACCGTAACCTTATTGTCCATATACAGGAGAAGGTTATGACAGAACTCACCTTGCAGAATCATCGTCGTACAATGTGGCATTTTATACCGGGGCTCGCTCTGAGCGCCGTCATTACAGGAGTCGCCCTGTGGGGCGGCGCTATCCCTGCCGTTGCAGGCGCGGGGTTCAGCGCCCTGACCCTGGCTATCCTGCTGGGTATGGTTATCGGGAATACCATCTATCCACAAATATGGAAACAATGCGACGGCGGCGTGCTGTTTGCAAAACAACATCTGCTGCGTTTAGGGATTATCCTTTACGGCTTCCGCCTTACCTTTTCACAAATCGCCGACGTGGGCATTAGCGGCATTGTGATTGACGTATTAACGCTTTCCAGCACGTTCATGCTGGCCTGCTTCTTAGGCCAAAAAGTCTTTGGTCTGGACAGGCACACCAGCTGGCTGATTGGCGCCGGTAGCAGTATCTGCGGCGCGGCGGCGGTACTGGCGACAGAACCGGTGGTAAAAGCGGAAGCCAGTAAGGTTACTGTAGCCGTCGCGACTGTCGTTATTTTCGGCACCATCGCTATTTTCCTTTACCCGGCAATGTATCCGCTGCTGGCGCACTGGTTCAGCCCGGAAACTTACGGCATCTATATCGGCTCAACCATGCATGAAGTCGCGCAAGTGGTCGCAGCGGGTCATGCCGTTAGCCCGGATGCGGAAAATGCGGCAGTTATCGCTAAGATGCTGCGTGTCATGATGCTGGCGCCGTTCCTGATCATTCTGGCGGCACGCGTTAAACAGCTTTCACCGGCGACCGGCGCGGAAAAAAGCAAAATTACCATTCCGTGGTTTGCTATTTTCTTCATCGTAGTGGCGATCTTCAACTCTTTCCACCTGCTGCCGAAAGCCGTGGTGGATATGCTGGTGACGCTGGACACGGTGCTGCTGGCGATGGCGATGGCGGCGTTGGGTCTGACCACTCACGTTAGCGCATTGAAAAAAGCGGGGGCGAAACCGCTGCTGATGGCGCTGGCATTATTCGCCTGGCTGATTATTGGCGGTGGCGCCATTAACGTCCTAATCCATAGCCTTATCGCATAAATTATTACGTTTCACGCCTGTTAACCCGTTATCATAGCGTTCGTAATGAACCGTTAGCGGGTTTAACAGGAGTCTTTTATGAAATACATCGGAGCGCACGTCAGCGCTGCTGGCGGTCTGGCTAACGCCCCTGCCCGCGCGGCTGAAATTGGCGCAACGGCCTTTGCGCTTTTCACAAAAAACCAGCGTCAGTGGCGTGCCGCCCCCCTTACTCCCCAGGTCATTGATGACTTTAAAATCGCCTGTGAAAAGTATCATTTCTCGGCGGCGCAAATCCTTCCCCACGATAGTTACCTGATTAATCTGGGCCATCCGGTCAGTGAAGCGCTGGAAAAATCACGCGATGCCTTTCTCGATGAAATGCAGCGCTGTGAACAACTCGGCTTAACCTTGCTTAATTTCATCCCGGTAGCCATCTGATGCAGATTGCACAGGAGGATTGCCTGGCGCGGATCGCGGAATCCATCAATATTGCCCTCGCGCAGACCGAGGGCGTTACGGCGGTTATCGAAAATACAGCCGGTCAGGGCAGTAATCTGGGGTTTGAGTTTGAACAGTTAGCCGCCATCATCGACGGCGTGGAAGATAAGTCGCGCGTTGGCGTCTGTATCGATACCTGCCATGCCTTTGCCGCCGGATACGATC
>T-11
GATCTGGATGATATCTTCCGTAAGCTGAGTAAAAAACTCGGTGGTTTTGGGGGCGGTAAAGGCACCGGTTCCGGCGGCGGTAGCTCATCGCAAGGCCCACGTCCGCAACTGGGAGGCCGTATTGTCGCCATCGCCGCGGCGGCAGTAGTCATTATCTGGGCGGCCAGCGGTTTTTATACCATTAAAGAAGCCGAGCGCGGCGTTGTGACGCGTTTTGGTAAATTCAGCCATCTGGTAGAGCCAGGTCTGAACTGGAAGCCGACATTTATTGACGACGTCACGCCGGTGAACGTCGAAGCGGTGCGCGAACTGGCGGCGTCAGGCGTGATGCTGACGTCTGATGAAAACGTCGTGCGCGTTGAAATGAACGTGCAATACCGCGTCACCGATCCGCAGAAATATCTGTTTAGCGTGACCAGCCCGGACGATAGCCTGCGTCAGGCCACCGACAGCGCGCTGCGTGGCGTCATTGGTAAATACACGATGGATCGCATCCTGACCGAAGGTCGTACCGTTATTCGTAGCGATACCCAACGCGAGCTGGAAGAGACTATCAAACCGTACAACATGGGGATTACCCTGCTGGACGTGAACTTCCAGGCTGCGCGCCCGCCGGAAGAGGTGAAAGCCGCCTTTGACGACGCGATTGCCGCACGTGAAAACGAGCAGCAGTACATTCGTGAAGCGGAAGCGTATACCAACGAAGTTCAGCCGCGCGCCAACGGTCAGGCGCAGCGTATTCTGGAAGAGGCACGCGCCTATAAAACCCAGACTATCCTGGAAGCGCAGGGTGAAGTCGCCCGGTTTGCGAAGATTTTGCCGGAATATAAAGCCGCGCCGCAGATTACACGCGAGCGTCTGTATATTGAAACGATGGAAAAAGTGCTGAGCCATACCCGTAAAGTACTGGTTAACGACAAGAGCGGTAATTTAATGGTGCTGCCGTTAGATC
>T-12
GATCGGGAAAAAGCACCACCGGTCTGGCGCTGTTACGGCTTATCCGCTCCGAAGGCCGCATCGTGTTTGACGGTCAATCGCTGGATACGTTAAACCGCCGCCAGCTTTTACCTGTTCGCCACCGTATCCAGGTCGTATTCCAGGACCCGAACTCATCGCTAAACCCGCGTTTAAACGTATTGCAAATTATCGAAGAAGGCCTGCGCGTCCACCAGCCTACGCTTTCAGGCGCGCAGCGCGAACAGCAGGTGAAAGCGGTCATGATGGAAGTCGGCCTGGACCCGGAAACGCGGCATCGTTACCCCGCTGAGTTTTCCGGCGGCCAGCGTCAACGTATCGCCGTCGCCAGGGCACTGATTTTAAAACCGTCGCTTATTATTCTGGATGAACCGACCTCATCGCTGGATAAAACCGTTCAGGCGCAGATTCTTGCCCTCCTGAAATCGCTCCAGCAAAAGCACCGTCTGGCCTATATCTTCATTAGCCACGATCTGCATGTAGTACGCGCGCTGTGCCATCAGGTTATTGTGCTGCGGCAGGGGGAGGTGGTTGAACAGGGGCAATGCGAGCGCGTGTTTACCGCACCGCAACAGGCCTATACGCGTCAGCTACTCGCGTTAAGCTGACGCCTAAAAAGGATTATTGTTAGAAAAAGGTTCCGCTATCGCCACGCCGAAATTTTTCAGCCGACAGGTGGCGGCAAACTCATCCTGACGATTAACGAACAGGCACGGCTCCCCTTCGCACTCCAGCACGGAGCAAGGAACTTCAATATCGCTTAGCGCCTGGCTGAGCTTATGCATCACTGACCACGCGCTATCGCCTGCCGGACTCAACAGTTTAAGCGCCACCAGACTATTCTCGGTGCTCTGCCCGTTTTGCGGAATCGGTTCAACCTTTGACCCCGCAAAACCCGCCGACCAGCGATAGTTTTGCGGCAGACGGTGAACAATTGAGAGCTGTAATGTATTCACTTTCTTTCCCCGGAAGCACAATTACTTCACAATTTACTTACATTTATATTAACACATCATTGACAACCCGTCCTTAAACAGATTGCTAACGCCCTTCTCTCCGTCAGTGGCTCTACAGCAAGCGGGAGTTAAAATAGTTTTACCGACACGGAGATAGCGGCGGGCTATATGTACCCGTTTCTGCGATC
>T-13
GATCTTCTGCGCCAGGTCGCATCTTACGGCCGCGGTTACACCTACCTGCTTTCGCGTTCGGGTGTCACCGGCGCGGAAAACCGTGGCGCATTGCCGTTGCATCATCTCATTGAGAAGCTTAAAGAGTACCATGCCGCGCCTGCGTTACAGGGCTTCGGTATCTCCTCGCCGGAACAGGTGTCTGCGGCCGTGCGTGCCGGGGCGGCTGGCGCTATCTCCGGCTCAGCCATTGTCAAGATTATCGAGAAAAACCTCGCGTCTCCCAAACAGATGTTGGCGGAGCTCAGGTCCTTTGTCTCAGCCATGAAAGCCGCCAGCCGCGCATAACGGGTTAAGCCGTCAGCATAACGCTGGCGGCTTAATGAGTGGCTGGCGCCGAACCAGAATCACGATTCAGACGTACATCATCAATCTTTTTGGTGTTATTCCCCGTCCAGCTTTCAAAAAGTCTGAAGTCAATTTTTGCTAAGAAAACTCCTAATCACGCCGAAAAATTTAGCGTTTTCACGTCAATTTCCCCCGCCTTTATGTAAAGAAATGTCAACAAACAATGCATTAACGTTTCCAGGGCCGCAGATAGTGACGATTGTTATTAATAACAATGCAAAGCATTTCCTGTTAAATCGCTTATTCTAAAAGAAAATATAGAATATATTATCTCTAAACCATTTTTTGTGCGTTATTTACGCGGTAAATCACAACTATTTCCGTCAATTTGACTAATCGGTTTAACCAACTAATTTTAATAGGGTGTCGACACGGTTAAAACCGATTTTTTCAGCAAGCAACGAGACAGGAGAAATAATATGGCCGAACATCGTGGTGGTTCAGGAAATTTCGCCGAGAATCGTGAAAAAGCATCTGAAGCGGGCCGCAAAGGCGGTCAGCATAGCGGCGGGAATTTCAAAAACGATC
>T-14
GATCTGGAAAAAATATATAAACTTCTTATTGAAAATCAATAGCTTAAGATATAGCCTCAACTATGTTTTTCGAATTTTATTTCGAATCATAATATAGCTAAGGCTATATTTTCTATTGACCAATTGTTATTTAATTTAACGTTTTATTGATGAATAAATCTAAATGAAAATCGTTTTCAATTGAAAGACAAATAATATTTTATCGCTGTTGTTTTATATTCTAAAAATATGTTAATGTTGCGCCGTCAATTGGGGAGTAGCCGATTTCCAGACTCCGGAAATGTACGTGTCAACATACTCGTTGCAAAACGTGGCGCGTACGGACCAGCCGCCGCTGGTCAGGCGAGACCATAGGCGCATCAACTGCTGTGTATTTCGTCACAGAATGGCGACATGCATGTTTGCTGGGGGCAGTGATGTGTTTAATGGATACCCCGGTCAGGACGTTGTCATGCACTTTACTGCTACTGTTCTTCTCGCTTTCGGCATGTCGATGGATGCTTTCGCGGCATCAATTGGCAAAGGCGCCACCCTACATAAACCCAAATTCTCAGAAGCGTTGCGTACCGGTCTTATCTTTGGCGCGGTTGAAACGCTAACCCCGTTGATCGGCTGGGGGCTGGGGATATTGGCGAGTAAATTCGTTCTGGAATGGAACCACTGGATAGCCTTTGTTCTGCTGATCTTTCTGGGCGGACGCATGATC
>T-15
ATCGGTCTGGTGAATGCCCGCGGTGAGAGGATATTTGAGACCCGCCTGAAACCCACAGTCGGCATCGATC
>T-16
GATCGCCCCCATCGTGATAAAGGTATAGTGGAACTGTTCCACGGTACTGGTGCCCGCAAAACCTTCATAAATACGTAACACCGCCGCGCTGATCGCCACGCCTAAACTGATAGACAATTGCTGCGTAACCGCCAGAACGCTGTTGCCGCTGCTGGCATTATCGTCCGTCAGATC
>T-17
GATCTTCTGCGCCAGGTCGCATCTTACGGCCGCGGTTACACCTACCTGCTTTCGCGTTCGGGTGTCACCGGCGCGGAAAACCGTGGCGCATTGCCGTTGCATCATCTCATTGAGAAGCTTAAAGAGTACCATGCCGCGCCTGCGTTACAGGGCTTCGGTATCTCCTCGCCGGAACAGGTGTCTGCGGCCGTGCGTGCCGGGGCGGCTGGCGCTATCTCCGGCTCAGCCATTGTCAAGATTATCGAGAAAAACCTCGCGTCTCCCAAACAGATGTTGGCGGAGCTCAGGTCCTTTGTCTCAGCCATGAAAGCCGCCAGCCGCGCATAACGGGTTAAGCCGTCAGCATAACGCTGGCGGCTTAATGAGTGGCTGGCGCCGAACCAGAATCACGATTCAGACGTACATCATCAATCTTTTTGGTGTTATTCCCCGTCCAGCTTTCAAAAAGTCTGAAGTCAATTTTTGCTAAGAAAACTCCTAATCACGCCGAAAAATTTAGCGTTTTCACGTCAATTTCCCCCGCCTTTATGTAAAGAAATGTCAACAAACAATGCATTAACGTTTCCAGGGCCGCAGATAGTGACGATTGTTATTAATAACAATGCAAAGCATTTCCTGTTAAATCGCTTATTCTAAAAGAAAATATAGAATATATTATCTCTAAACCATTTTTTGTGCGTTATTTACGCGGTAAATCACAACTATTTCCGTCAATTTGACTAATCGGTTTAACCAACTAATTTTAATAGGGTGTCGACACGGTTAAAACCGATTTTTTCAGCAAGCAACGAGACAGGAGAAATAATATGGCCGAACATCGTGGTGGTTCAGGAAATTTCGCCGAGAATCGTGAAAAAGCATCTGAAGCGGGCCGCAAAGGCGGTCAGCATAGCGGCGGGAATTTCAAAAACGATC
>T-18
GATCGTCCTTATCATGATTAGCGGTTACCGCGTCCCAGAGTTTTTCACCTGCATCTTTTACAAAATTGAAAAGTCCCATATTCACCTCGCTTGTCAAAAATACACACAAAGAAAAGTGTAGCAGAGGGAATGAGGTTATCCTGATAAGCGGCGCTTATGCGCCGCCGGTGGCCGGTAATTAATGACGCATCTGAACCCAAAATGCGTGGATAAGCCCCGGAATATATCCGAGTAATGTCAGCAGGATATTAAGGATAAATGCCCAACCGAACCCTTTTCCTAAAAGTACGCCGAGCGGAGGCAAAATAATCGTAAATACAATTCTCCAGAAACCCATATACACTCCATACAAGCAAAATTAGTCATTGTTTAAAAAGAGATTTTATCTCTAAGCATAGTCAGTTTACCTGAGCCTGCCAGTCGCGCGCCGCGCTTTTACCCTCCTTTACGCTCTTTACGCTGGCATTAGGTTGATGTTTATTTTAGAATTAACTAAATTTAGAAAATACTTAAATAACTATGACAGAGCTTGAACAACTTCAGGCCAGCGCAGAACAGGCGGCGGCCCTGCTCAAAGCGATGAGCCACCCAAAAAGATTACTGATTTTATGTATGCTTTGCGGTTCGCCAAAAACCAGCGCAGGCGAACTGGCGCGCATTACCGGGCTAAGCCCCTCCGCGACGTCGCAACATCTGGCGCGGATGCGGGAAGAAGGGCTGATTGATAGTCAACGCGATGCCCAGCGCATTCACTATTTCATTAAAAATGAGGCGGTTAATACGATTATCGCCACGTTAAAGAATCTCTATTGTCCATAAGGAGGCACGATGTCCATTGGGATAGTTTCCCCCCGTGAGGCGCAAGCGCTGATAGCGCAAGGCGCGAAACTGATTGATGTGCGCGACGCCGATGAATATCTGCGCGAGCATATTCCACATGCGCAACTGGCGCCGCTCTCCCGGCTTGAACAGGGCGATTTGCCCGCCAACCTGCGAGCGGAACAGATTATTTTTCACTGTCAGTCAGGTAAACGAACCAGCAGCAATGCAGCAAAACTACAGGCCATTGCCGCCCCTGCTCAGGTCAGTCTACTGGAAGGCGGTATCGACGGATGGAAAGCGGCAGGATTGCCCGTCACAGAAGACAAATCGCAACCGTTGCCGCTGATGCGTCAGGTGCAAATCGCCGCCGGCGGGTTGGCGTTGCTTGGGGTTATTATGGGCTATACGGTCCACGGCGGCTTTTTCCTCATTAGCGGTTTTGTCGGGGCCGGGTTAATGCTGGCAGGCATGACCGGCTTCTGCGGAATGGCGCGACTACTGGAAAAAATGCCCTGGAATACCCGTACTCATTAAGCGGTACCTTCACCACGCGTCGGTAAGTGTGTAGTAGACTGAAGGCGGGTTTATACCATTGTTCAACAGGAGGGCGTGATGTTCACACCAGGCGATATCGTACAGCCCCGAATGGGTGGGCCGAAACTGAAAGTGATTGAAGTCAACGAGGATC
>T-19
GATCAGATTCGCCGGTTAATGCAGCGGGATAAAACGCCGCTGGCCATCCTGTTCATGGCGGCCGTTGTCGGCACGCTTACCGGACTGGTTGGCGTCGCCTTTGAAAAGGCGGTTTCCTGGGTACAAAACATGCGCATTGGCGCGCTGGTTCAGGTCGCGGACCATGCGTTTTTGCTTTGGCCGCTGGCCTTTATCCTTTCGGCGTTACTGGCGATGGTCGGCTACTTTTTGGTGCGAAAATTTGCGCCGGAAGCCGGCGGCTCGGGTATCCCGGAAATTGAAGGTGCGCTGGAGGAGTTACGTCCGGTGCGCTGGTGGCGCGTGTTGCCGGTGAAGTTTATCGGCGGGATGGGAACGTTGGGCGCAGGAATGGTATTAGGGCGCGAAGGGCCAACGGTGCAAATCGGGGGCAACCTGGGGCGCATGGTGCTGGATGTTTTTCGTATGCGCAGCGCCGAAGCGCGGCATACGTTGCTGGCGACCGGCGCGGCGGCGGGGCTCTCTGCGGCGTTTAACGCGCCGCTGGCGGGTATCCTGTTTATTATTGAAGAGATGCGCCCGCAGTTTCGCTACAACCTGATTTCGATTAAAGCGGTGTTTACCGGCGTCATCATGTCGAGCATTGTGTTCCGCATATTTAACGGCGAAGCGCCGATTATTGAAGTCGGCAAACTGTCTGATGCGCCGGTTAACACGCTGTGGTTATACCTTATCCTCGGTATTATTTTTGGCTGCGTCGGGCCGGTATTTAATTCGCTGGTGCTGCGTACCCAGGATATGTTTCAGCGCTTTCATGGCGGCGAAATTAAAAAATGGGTGCTTATGGGCGGCGCGATCGGCGGCCTGTGCGGTATTCTGGGGTTGATTGAGCCGGCCGCGGCAGGCGGCGGATTTAACCTTATTCCTATCGCCGCGGCAGGTAATTTTAGCGTCGGGCTACTGCTGTTCATCTTTATTACCCGGGTTGTCACGACGCTGCTTTGCTTTTCTTCCGGCGCGCCGGGGGGCATTTTCGCCCCGATGCTGGCGCTGGGAACGCTGCTTGGTACGGCCTTCGGTATGGCGGCGGCAGTGCTTTTCCCGCAGTATCACCTGGAGGCCGGTACCTTTGCGATTGCCGGGATGGGGGCGCTGATGGCGGCGTCGGTGCGCGCCCCGTTGACAGGGATTGTGCTGGTGCTGGAGATGACCGATAACTATCAGCTCATTTTGCCAATGATTATCACCTGTCTTGGCGCGACACTATTAGCCCAGTTTTTGGGCGGGAAACCGTTATACTCAACCATCCTGGCGCGCACTCTGGCGAAGCAGGATGCGGAACAGGCGGCGAAAAACCAGAATGCGCCTGCGGACGAGAATACTTGAACGAATTACCAGGGTATTAGATAATGACTGCCAATATTGGGTGAAAATTTATCCAATTACCGATGTCGTTTGGAGCAAAATATGAGTGATGACGTAGCGCTGCCGCTGCAGTTTACCGACGCAGCAGCGAATAAAGTAAAAAGTCTTATCGCTGACGAAGATAACCCGAATCTGAAACTGCGCGTGTATATCACCGGCGGTGGTTGCAGCGGCTTCCAGTATGGTTTTACCTTTGACGATC
>T-20
GATCAGGTAGGCAACGTCTTTAAGGGCGTGATTGCCAGCGTCACCGGTTTCGGCTTCTTTGTCCGTCTTGATGAACTGTTTATCGATGGGCTGGTACACGTTTCCTCACTGGATAACGATTACTATCGCTTCGATCAGGTAGGGCAACGTCTGATTGGCGAGTCCGGCGGGCAGACATATCGCCTGGGCGACCGGGTCGAGGTACGCGTCGAAGCGGTGAATATGGACGAACGGAAAATTGACTTTAGCCTGATCTCCAGCGAGCGCGCACCGCGTAACGTCGGCAAAACGGCGCGTGAAAAAGCGAAAAAAGGTGAAAGTAAAAACGCCGGTAAACGTCGTCAGGTAGGTAAAAAAGTTAACTTCGAGCCTGATAGCGCCTTCCGCGGCGAGAAAAAAGCGAAGCCGAAAGCAGCGAAGAAAGACGCGAGAAAAGCAAAAAAGCCATCGGCTAAAACGCAGAAAATAGCCGCAGCGACCAAAGCGAAGCGTGCGGCGAAGAAAAAAGCGGCAGAGTGAGCAATACCCTCTTCAAACAGAAGAGGGTCAATTATCTGTAGGCCGGGCAAGCGTAGCGCCCCCGGCAAAACGTAAACAACGAGTACATTAATGAGCGAAATGATTTACGGCATCCACGCAGTGCAGGCCCTGCTGGAGCGCGCCCCTGAACGTTTTCAGGAAGTCTTTATTTTAAAAGGCCGTGAAGATAAACGTCTGTTGCCGCTGATCCACGCCCTTGAGTCCCAGGGCGTGGTTATCCAGCTGGCAAACCGCCAATATCTCGACGAGAAAAGCGACGGTGCCGTGCATCAGGGTATTATCGCTCGCGTGAAACCAGGCCGGCAGTATCAGGAAAACGATC
>T-21
GATCTTCTGCGCCAGGTCGCATCTTACGGCCGCGGTTACACCTACCTGCTTTCGCGTTCGGGTGTCACCGGCGCGGAAAACCGTGGCGCATTGCCGTTGCATCATCTCATTGAGAAGCTTAAAGAGTACCATGCCGCGCCTGCGTTACAGGGCTTCGGTATCTCCTCGCCGGAACAGGTGTCTGCGGCCGTGCGTGCCGGGGCGGCTGGCGCTATCTCCGGCTCAGCCATTGTCAAGATTATCGAGAAAAACCTCGCGTCTCCCAAACAGATGTTGGCGGAGCTCAGGTCCTTTGTCTCAGCCATGAAAGCCGCCAGCCGCGCATAACGGGTTAAGCCGTCAGCATAACGCTGGCGGCTTAATGAGTGGCTGGCGCCGAACCAGAATCACGATTCAGACGTACATCATCAATCTTTTTGGTGTTATTCCCCGTCCAGCTTTCAAAAAGTCTGAAGTCAATTTTTGCTAAGAAAACTCCTAATCACGCCGAAAAATTTAGCGTTTTCACGTCAATTTCCCCCGCCTTTATGTAAAGAAATGTCAACAAACAATGCATTAACGTTTCCAGGGCCGCAGATAGTGACGATTGTTATTAATAACAATGCAAAGCATTTCCTGTTAAATCGCTTATTCTAAAAGAAAATATAGAATATATTATCTCTAAACCATTTTTTGTGCGTTATTTACGCGGTAAATCACAACTATTTCCGTCAATTTGACTAATCGGTTTAACCAACTAATTTTAATAGGGTGTCGACACGGTTAAAACCGATTTTTTCAGCAAGCAACGAGACAGGAGAAATAATATGGCCGAACATCGTGGTGGTTCAGGAAATTTCGCCGAGAATCGTGAAAAAGCATCTGAAGCGGGCCGCAAAGGCGGTCAGCATAGCGGCGGGAATTTCAAAAACGATC
>T-22
GATCTGCATACTTCGCAATATCAGGTGCTCACACGTCTGGCGCAGTTCTCACTGGCGCGCGAACCGCATACTGACAAAGTAGAAACCCCCAAAGCGGCGGTGTCGATGGCGAAGAAGGCCTCGTTTGTGGTGGTGGTGATTATTGCGTTGCTCACCATTTATGGGGCATTAGTGTAGCGTTGTGCCGATGGCGCTAACGCTTATGCTGCCTACAATGGAAAAATCTGTAGGCCCGGTAAGCGTTAGCGCCACCGGGCAGTGCAACGGCTTAGATCGTATCGGCTGGCGGCACGCCAAAATCAGGCATCCCGTTTTCGTCCCAGCGCACCCGTTTCAGACGCGTATGGCGGTTGGGATCGTACAACGGATCGCCTTCAATTTCGGTGTAATTTCTCGCGTGGTACACCAGTACATTTTCACCTTCCGGCGTTTGCGTAAAGCTGTTGTGTCCCGGTCCATACTGGCGATTCTCATAGCTGGTGGTAAAGACCGGGCGCGGTGATTTATGCCAGTTGGCCGGATCGCGCGGATCGTCGTTTACGTTAATCCACAGTAATCCCATACAGTAGTTTTCATCGGTGGCGCTGGCCGAATAGCTGATAAACAGCTTGTCGCCATGCACCACGACGGCCGGACCTTCGTTAACCCAAAAACCCCGACACTCCCAGTCGTACTCTGGCTTACTGAGTCTTACCGGCTCGCCTTTAATCGTCCACGGATTTTCCAGTTCAGCCAGATAAATATTTGAGTTACCGGCAATATCCGGCGCTTTTTGCGCCCACAGATACCACTGCTTTCCCTGATGATAAAAAGTGGTGGCATCGAGGGCGAAGGTATCAAACGGTGTTTTAATTTGACCTTTCTCCGTCCACTTGCCGGTGAGTGGGTCGGCATCGGCGCACTCCAGGGCAAACATGCGATGCTGGAACATTCCCAGCTTATCAAGCGCTTGAGTATGGGTAGCGGCAAAATAGATGTACCACTTGCCGTTGATGCGGTGCATTTCCGGTGCCCAGATGAGCTGGCTCATCGGGCCGCTTTCCGGTTTACGCCAGACGACAACGGGGTCGGCGGCGCGCAACCCTTCCAGCGAATTAGCCCGGCGGATTTCCAGCCTGTCATATTCCGGTACGGAGGCAATAAAGTAGTAATCGCTGCCGTCGCGTAAAATAAACGGATCGGCACGTTGTTCAATAAACGGGTTTGGCCAGTTTGCCATCAGGCTTTCCTTACGCTTTCAGCGGCGTTGAGCTCCTGCAACTCACTCAATTCGCGGTAATTAGTGCGACGTTTTTCCAAATCGTCCTGGATTTTTTTCATCAGTTCACGGTCAACTTTCAGCAGACGTACCACGCCTGCGGTAATCAGATAGCCAACGCCGGGGATAACGGTAAACAACAGCATAATACCGTTCATCGCCGAGGCGCTTTGCGCTTTGGCGCTGGCGTCATAACCGTACCAGGAGAGCAAAAAGCCGACCATCGCCCCGGCAATCGCCAGGCCCAGTTTCAGGAAGAACAGGTTACCGGAGAAGCTGATACCGGTAATGCGTTTGCCGGTTTTCCATTCGCCGTAATCATCGACGTCCGCCATCAGCGACCAGTGCAGCGGGGAAGGGATC
>T-23
GATCTGGGTATCATTAATAAAATCCCAGCCGTAGTAGCCCTTCAGACACAGGGTACCCTGGTTGGTTTTCCCCTGCGCCGCCTCAGCCCGGACGATTTTGCCGTTATCGACCACCAGATTGATTTTGCAACCTGATGCGCAATACGGGCAAACCGTGACGACTTTTTTCATCGGTTTCGCTCCAGTTAATCAAATCGCGCATACGCGCTGTCGCCCTCAGTATGCATGTTTTATGCCACTTTTTTAGTATGGGCTTTCCCTGATATTACGGGCGAATTTTGCTCAAAACCCTGACGAAAAACAGGCTATCGTCAGTTTTGACGTGTCGAAAGAAAATGCCGCATGACATTTATTATGGGGAGAAACGCGGAGCGCAGGCCAGATAAGGCGGAGCCGCTATCCGGCAATCTGCTCTGCCTGAAGTGGAAATTATCCGGATGGATAGACTCAAGGCCAGGGAACGCTTTGAGACTAACGCCAGGATAGAACAGGCCGCGGCAGGCAGTTCCATACCGCGGCCCGGCTTCAGAGGAGGCTGAAAAGGATTATCGCGGCAAAAATAGCAGCAATAACGAGATATTTCAGCGCGTAATAGAGAGTGCGGTTTTTCTTTTTCAGACGTTGCCCGGCATCGCGTGCTGCGTGAACGTATTTAAAGATGCGATTAATCGCACCGGTGCGATC
>T-24
GATCTGGGTATCATTAATAAAATCCCAGCCGTAGTAGCCCTTCAGACACAGGGTACCCTGGTTGGTTTTCCCCTGCGCCGCCTCAGCCCGGACGATTTTGCCGTTATCGACCACCAGATTGATTTTGCAACCTGATGCGCAATACGGGCAAACCGTGACGACTTTTTTCATCGGTTTCGCTCCAGTTAATCAAATCGCGCATACGCGCTGTCGCCCTCAGTATGCATGTTTTATGCCACTTTTTTAGTATGGGCTTTCCCTGATATTACGGGCGAATTTTGCTCAAAACCCTGACGAAAAACAGGCTATCGTCAGTTTTGACGTGTCGAAAGAAAATGCCGCATGACATTTATTATGGGGAGAAACGCGGAGCGCAGGCCAGATAAGGCGGAGCCGCTATCCGGCAATCTGCTCTGCCTGAAGTGGAAATTATCCGGATGGATAGACTCAAGGCCAGGGAACGCTTTGAGACTAACGCCAGGATAGAACAGGCCGCGGCAGGCAGTTCCATACCGCGGCCCGGCTTCAGAGGAGGCTGAAAAGGATTATCGCGGCAAAAATAGCAGCAATAACGAGATATTTCAGCGCGTAATAGAGAGTGCGGTTTTTCTTTTTCAGACGTTGCCCGGCATCGCGTGCTGCGTGAACGTATTTAAAGATGCGATTAATCGCACCGGTGCGATC
>T-25
GATCAGATTCGCCGGTTAATGCAGCGGGATAAAACGCCGCTGGCCATCCTGTTCATGGCGGCCGTTGTCGGCACGCTTACCGGACTGGTTGGCGTCGCCTTTGAAAAGGCGGTTTCCTGGGTACAAAACATGCGCATTGGCGCGCTGGTTCAGGTCGCGGACCATGCGTTTTTGCTTTGGCCGCTGGCCTTTATCCTTTCGGCGTTACTGGCGATGGTCGGCTACTTTTTGGTGCGAAAATTTGCGCCGGAAGCCGGCGGCTCGGGTATCCCGGAAATTGAAGGTGCGCTGGAGGAGTTACGTCCGGTGCGCTGGTGGCGCGTGTTGCCGGTGAAGTTTATCGGCGGGATGGGAACGTTGGGCGCAGGAATGGTATTAGGGCGCGAAGGGCCAACGGTGCAAATCGGGGGCAACCTGGGGCGCATGGTGCTGGATGTTTTTCGTATGCGCAGCGCCGAAGCGCGGCATACGTTGCTGGCGACCGGCGCGGCGGCGGGGCTCTCTGCGGCGTTTAACGCGCCGCTGGCGGGTATCCTGTTTATTATTGAAGAGATGCGCCCGCAGTTTCGCTACAACCTGATTTCGATTAAAGCGGTGTTTACCGGCGTCATCATGTCGAGCATTGTGTTCCGCATATTTAACGGCGAAGCGCCGATTATTGAAGTCGGCAAACTGTCTGATGCGCCGGTTAACACGCTGTGGTTATACCTTATCCTCGGTATTATTTTTGGCTGCGTCGGGCCGGTATTTAATTCGCTGGTGCTGCGTACCCAGGATATGTTTCAGCGCTTTCATGGCGGCGAAATTAAAAAATGGGTGCTTATGGGCGGCGCGATCGGCGGCCTGTGCGGTATTCTGGGGTTGATTGAGCCGGCCGCGGCAGGCGGCGGATTTAACCTTATTCCTATCGCCGCGGCAGGTAATTTTAGCGTCGGGCTACTGCTGTTCATCTTTATTACCCGGGTTGTCACGACGCTGCTTTGCTTTTCTTCCGGCGCGCCGGGGGGCATTTTCGCCCCGATGCTGGCGCTGGGAACGCTGCTTGGTACGGCCTTCGGTATGGCGGCGGCAGTGCTTTTCCCGCAGTATCACCTGGAGGCCGGTACCTTTGCGATTGCCGGGATGGGGGCGCTGATGGCGGCGTCGGTGCGCGCCCCGTTGACAGGGATTGTGCTGGTGCTGGAGATGACCGATAACTATCAGCTCATTTTGCCAATGATTATCACCTGTCTTGGCGCGACACTATTAGCCCAGTTTTTGGGCGGGAAACCGTTATACTCAACCATCCTGGCGCGCACTCTGGCGAAGCAGGATGCGGAACAGGCGGCGAAAAACCAGAATGCGCCTGCGGACGAGAATACTTGAACGAATTACCAGGGTATTAGATAATGACTGCCAATATTGGGTGAAAATTTATCCAATTACCGATGTCGTTTGGAGCAAAATATGAGTGATGACGTAGCGCTGCCGCTGCAGTTTACCGACGCAGCAGCGAATAAAGTAAAAAGTCTTATCGCTGACGAAGATAACCCGAATCTGAAACTGCGCGTGTATATCACCGGCGGTGGTTGCAGCGGCTTCCAGTATGGTTTTACCTTTGACGATC

**Table S2.**
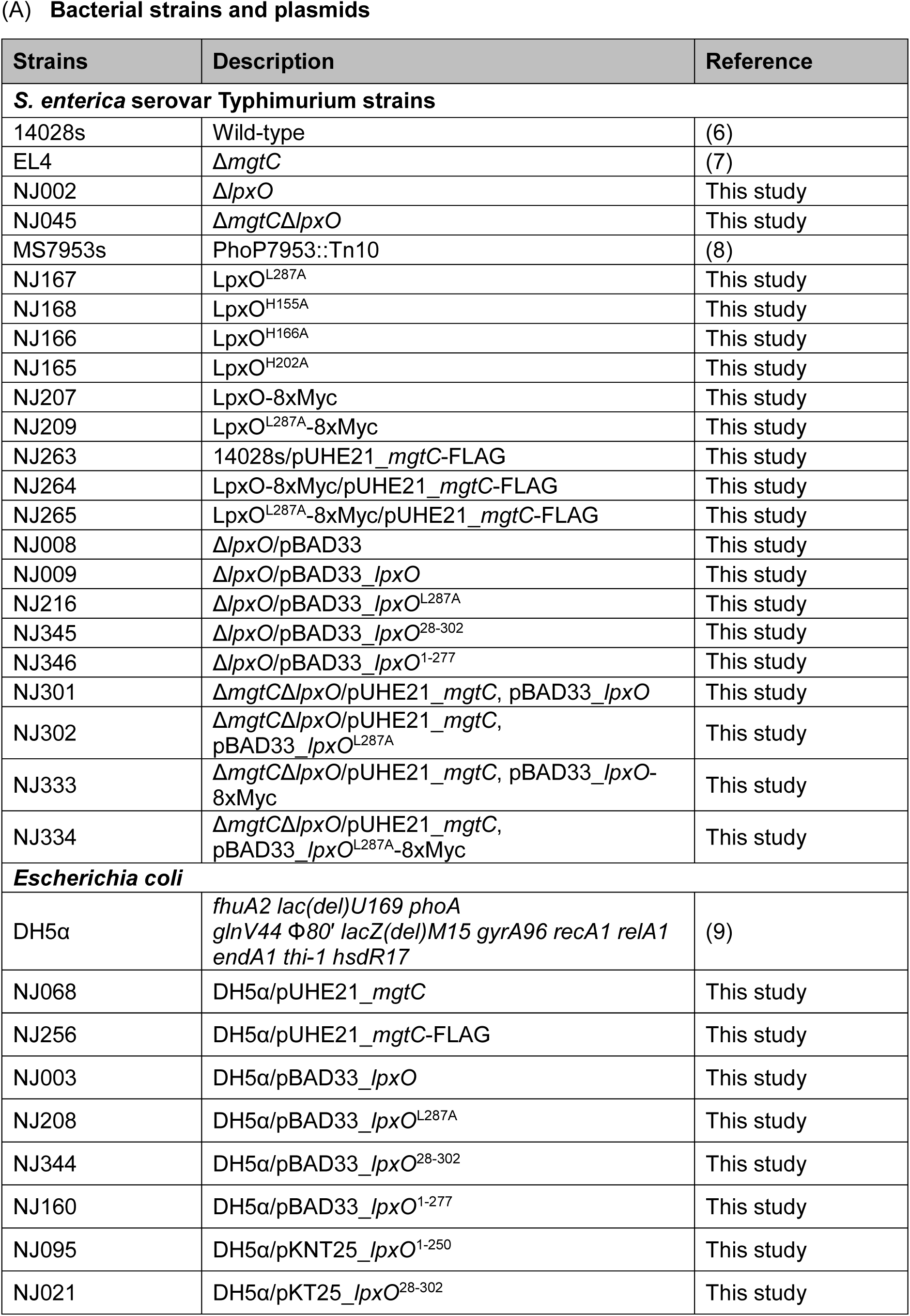
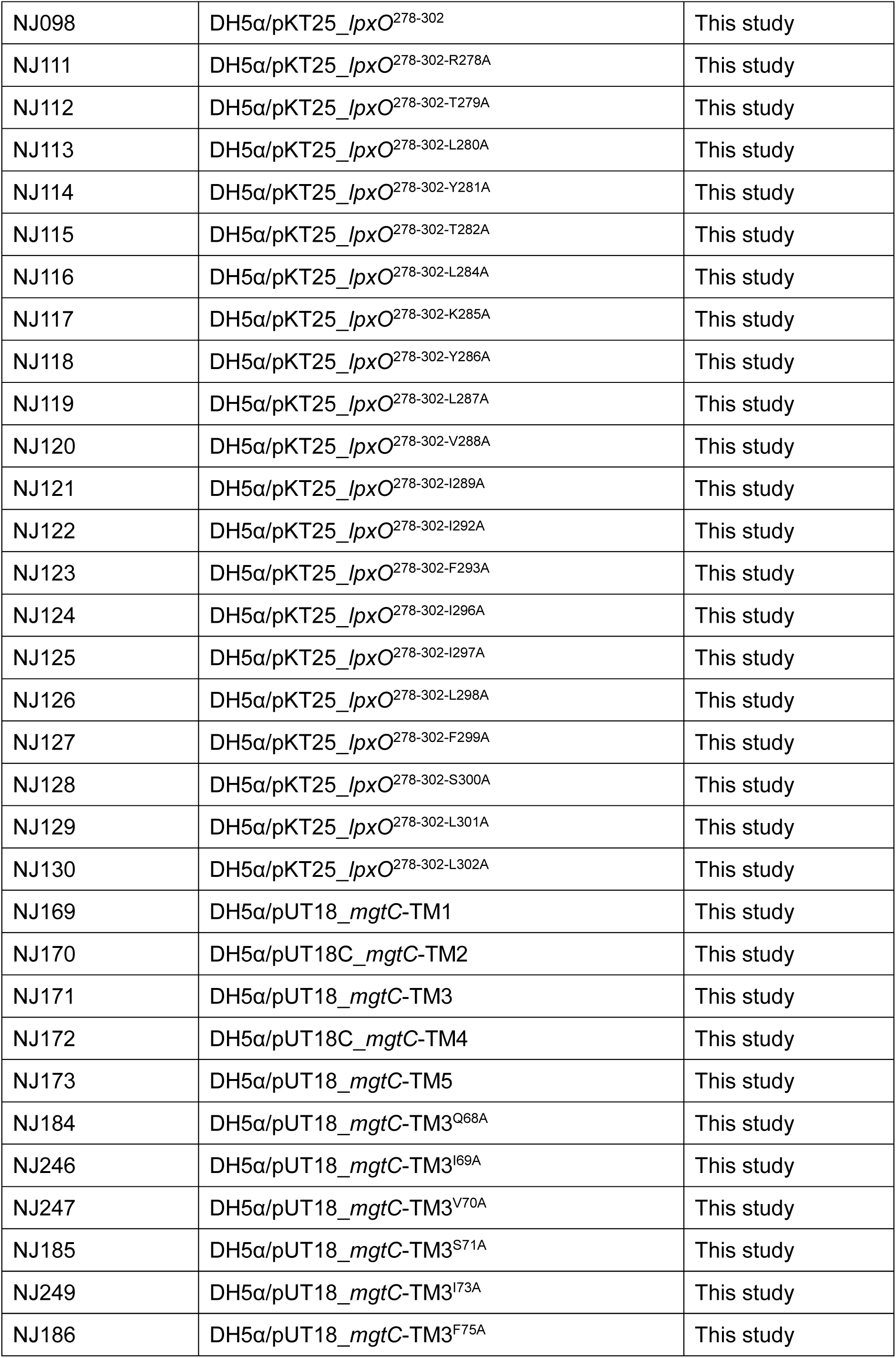
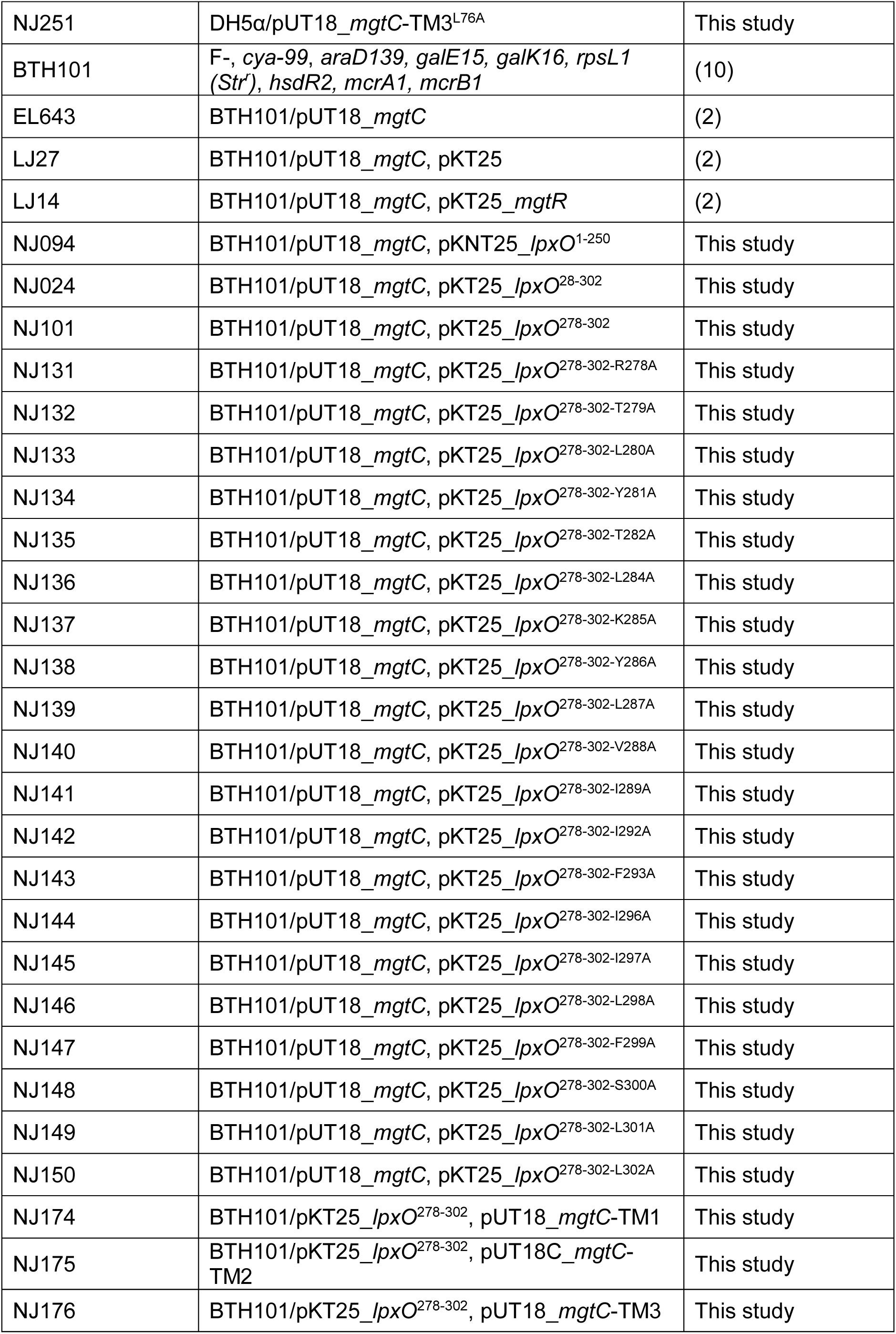
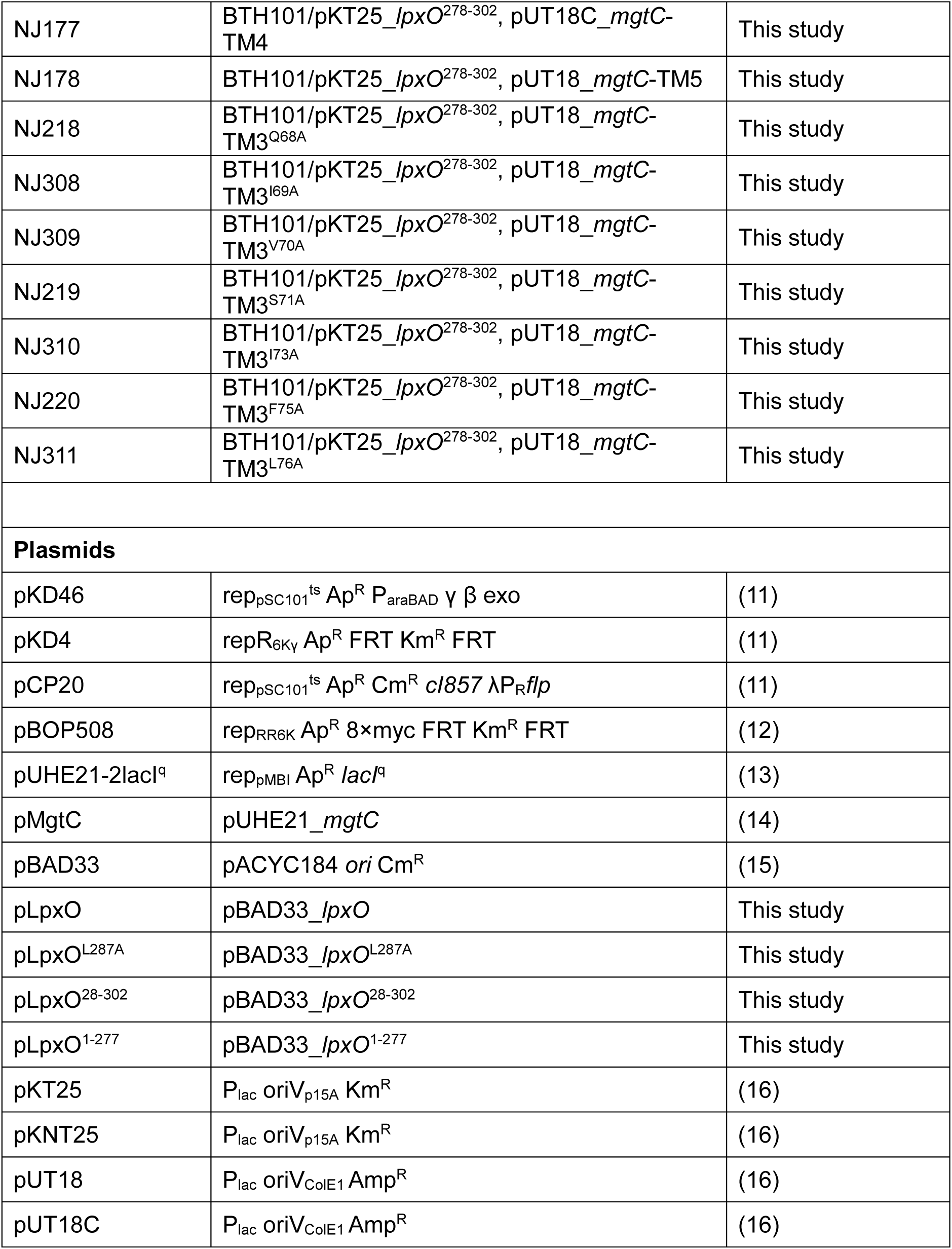
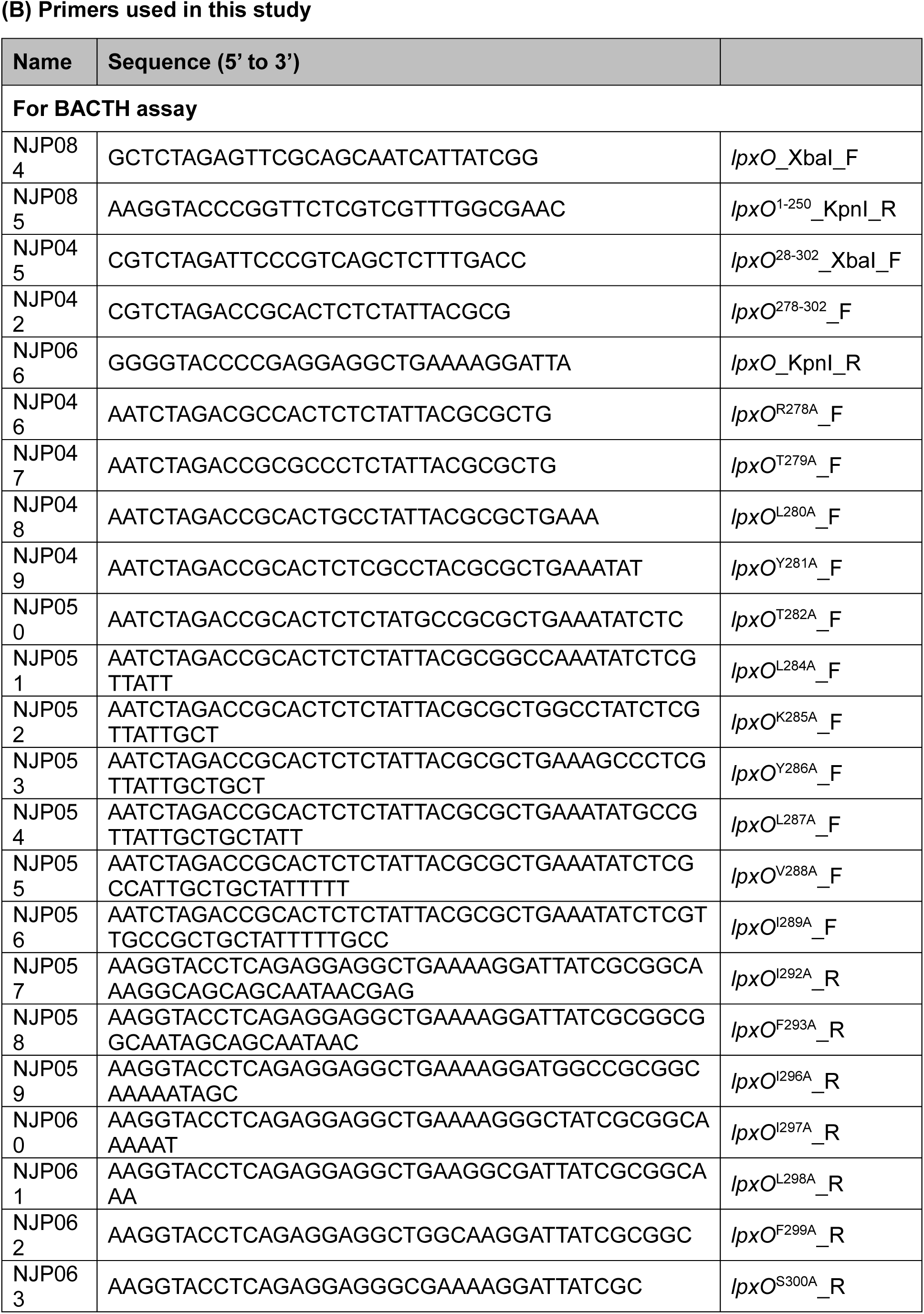
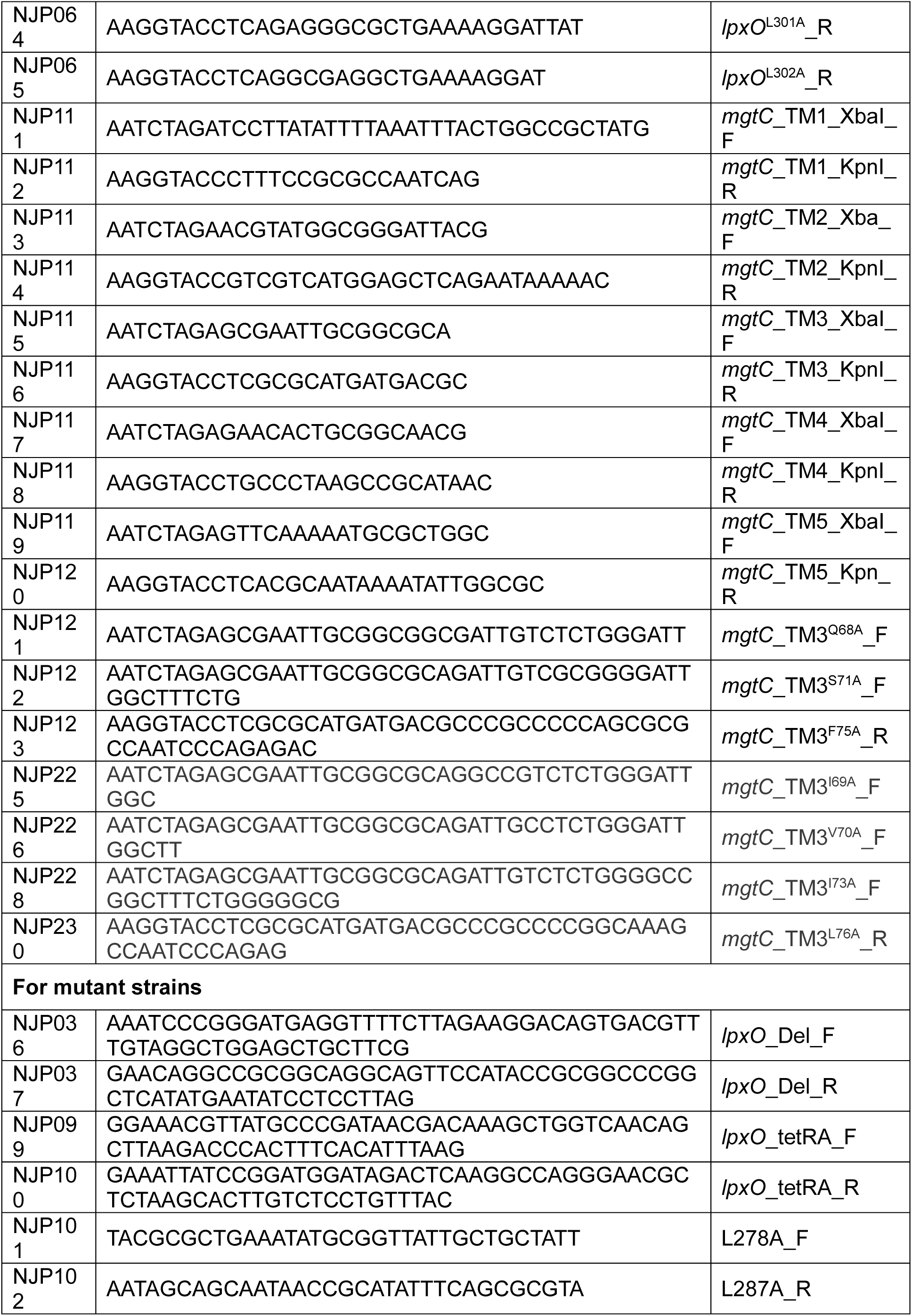
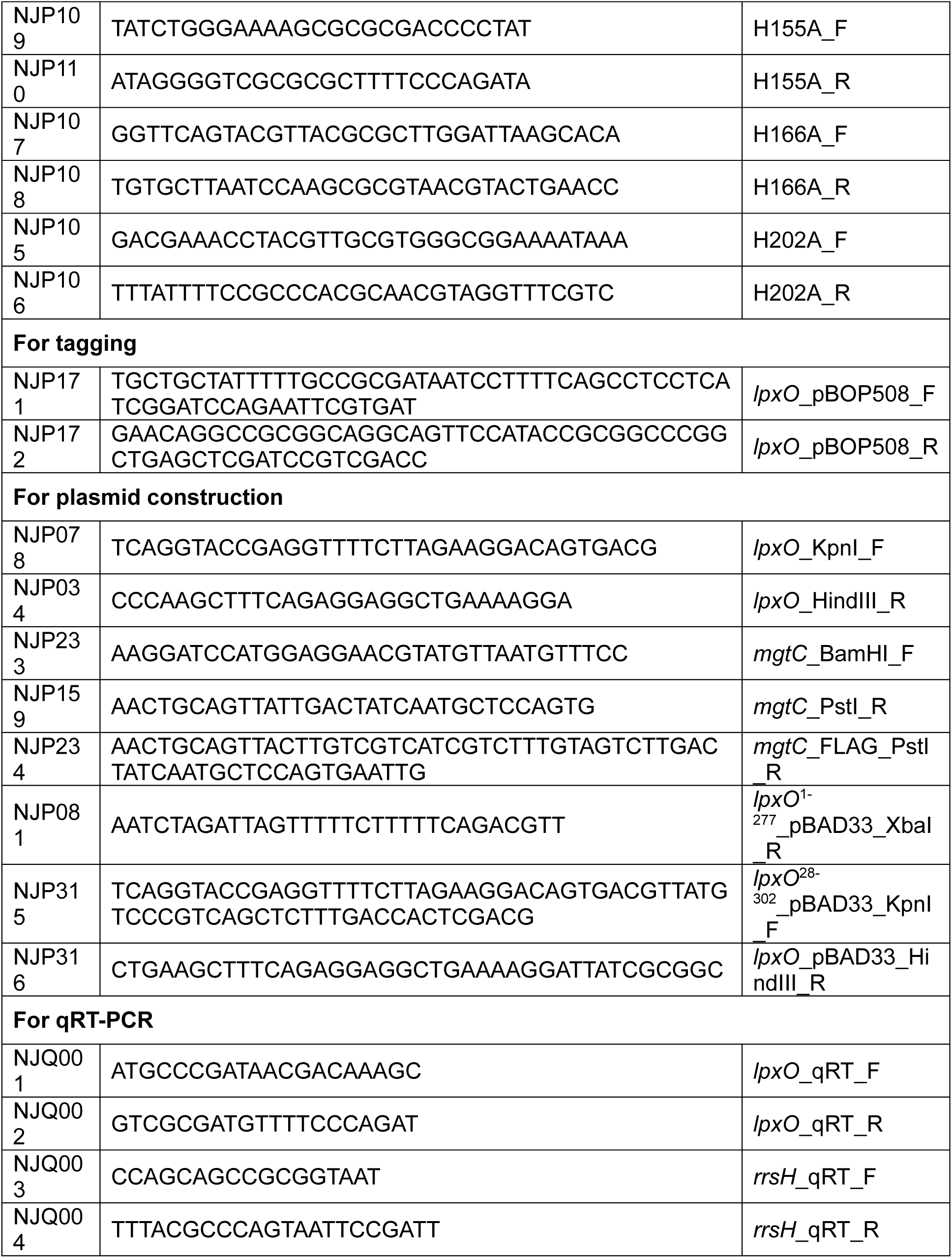
Bacterial strains, plasmids, and oligonucleotides used in this study.

## Legends for Datasets S1

## References

1. Uncategorized References

1. C. R. Raetz, C. Whitfield, Lipopolysaccharide endotoxins. Annual review of biochemistry 71, 635–700 (2002).

2. M. J. Trimble, P. Mlynárčik, M. Kolář, R. E. Hancock, Polymyxin: alternative mechanisms of action and resistance. Cold Spring Harbor perspectives in medicine 6, a025288 (2016).

3. C. R. Raetz, C. M. Reynolds, M. S. Trent, R. E. Bishop, Lipid A modification systems in gram-negative bacteria. Annu. Rev. Biochem. 76, 295–329 (2007).

4. C. R. Raetz, Regulated covalent modifications of lipid A. Journal of endotoxin research 7, 73–78 (2001).

5. M. E. Ohl, S. I. Miller, Salmonella: a model for bacterial pathogenesis. Annual review of medicine 52, 259–274 (2001).

6. H. S. Gibbons, S. R. Kalb, R. J. Cotter, C. R. Raetz, Role of Mg2+ and pH in the modification of Salmonella lipid A after endocytosis by macrophage tumour cells. Molecular microbiology 55, 425–440 (2005).

7. H. D. Chen, E. A. Groisman, The biology of the PmrA/PmrB two-component system: the major regulator of lipopolysaccharide modifications. Annu Rev Microbiol 67, 83–112 (2013).

8. A. Kato, T. Latifi, E. A. Groisman, Closing the loop: the PmrA/PmrB two-component system negatively controls expression of its posttranscriptional activator PmrD. Proc Natl Acad Sci U S A 100, 4706–4711 (2003).

9. W. M. Shafer, S. G. Casey, J. K. Spitznagel, Lipid A and resistance of Salmonella typhimurium to antimicrobial granule proteins of human neutrophil granulocytes. Infect Immun 43, 834–838 (1984).

10. A. X. Tran et al., Resistance to the antimicrobial peptide polymyxin requires myristoylation of Escherichia coli and Salmonella typhimurium lipid A. J Biol Chem 280, 28186–28194 (2005).

11. H. S. Gibbons, C. M. Reynolds, Z. Guan, C. R. Raetz, An inner membrane dioxygenase that generates the 2-hydroxymyristate moiety of Salmonella lipid A. Biochemistry 47, 2814–2825 (2008).

12. S. De Majumdar et al., Elucidation of the RamA regulon in Klebsiella pneumoniae reveals a role in LPS regulation. PLoS pathogens 11, e1004627 (2015).

13. A. Anandan, A. Vrielink, Structure and function of lipid A–modifying enzymes. Annals of the New York Academy of Sciences 1459, 19–37 (2020).

14. A. Lo Sciuto et al., Genetic basis and physiological effects of lipid A hydroxylation in Pseudomonas aeruginosa PAO1. Pathogens 8, 291 (2019).

15. L. Guo et al., Regulation of lipid A modifications by Salmonella typhimurium virulence genes phoP-phoQ. Science 276, 250–253 (1997).

16. T. Murata, W. Tseng, T. Guina, S. I. Miller, H. Nikaido, PhoPQ-mediated regulation produces a more robust permeability barrier in the outer membrane of Salmonella enterica serovar typhimurium. Journal of bacteriology 189, 7213–7222 (2007).

17. E. A. Groisman, The pleiotropic two-component regulatory system PhoP-PhoQ. Journal of bacteriology 183, 1835–1842 (2001).

18. A.-B. Blanc-Potard, E. A. Groisman, The Salmonella selC locus contains a pathogenicity island mediating intramacrophage survival. The EMBO journal 16, 5376–5385 (1997).

19. E.-J. Lee, M. H. Pontes, E. A. Groisman, A bacterial virulence protein promotes pathogenicity by inhibiting the bacterium’s own F1Fo ATP synthase. Cell 154, 146–156 (2013).

20. S. Choi et al., The Salmonella virulence protein MgtC promotes phosphate uptake inside macrophages. Nature communications 10, 3326 (2019).

21. E. Alix, A. B. Blanc-Potard, Peptide-assisted degradation of the Salmonella MgtC virulence factor. The EMBO journal 27, 546–557 (2008).

22. J. Yeom, K. J. Wayne, E. A. Groisman, Sequestration from Protease Adaptor Confers Differential Stability to Protease Substrate. Mol Cell 66, 234–246 e235 (2017).

23. E. J. Lee, E. A. Groisman, Control of a Salmonella virulence locus by an ATP-sensing leader messenger RNA. Nature 486, 271–275 (2012).

24. E. J. Lee, E. A. Groisman, Tandem attenuators control expression of the Salmonella mgtCBR virulence operon. Mol Microbiol 86, 212–224 (2012).

25. G. Karimova, J. Pidoux, A. Ullmann, D. Ladant, A bacterial two-hybrid system based on a reconstituted signal transduction pathway. Proceedings of the National Academy of Sciences 95, 5752–5756 (1998).

26. C. Rang et al., Dual role of the MgtC virulence factor in host and non-host environments. Molecular microbiology 63, 605–622 (2007).

27. A.-B. Blanc-Potard, B. Lafay, MgtC as a horizontally-acquired virulence factor of intracellular bacterial pathogens: evidence from molecular phylogeny and comparative genomics. Journal of molecular evolution 57, 479–486 (2003).

28. H. S. Gibbons, S. Lin, R. J. Cotter, C. R. Raetz, Oxygen requirement for the biosynthesis of theS-2-hydroxymyristate moiety in Salmonella typhimurium lipid A: FUNCTION OF LpxO, A new Fe2+/α-KETOGLUTARATE-DEPENDENT dioxygenase homologue. Journal of Biological Chemistry 275, 32940–32949 (2000).

29. M. Elshikh et al., Resazurin-based 96-well plate microdilution method for the determination of minimum inhibitory concentration of biosurfactants. Biotechnology letters 38, 1015–1019 (2016).

30. M. B. C. Moncrief, M. E. Maguire, Magnesium and the role of MgtC in growth of Salmonella typhimurium. Infection and immunity 66, 3802–3809 (1998).

31. E. Garcia Vescovi, F. C. Soncini, E. A. Groisman, Mg2+ as an extracellular signal: environmental regulation of Salmonella virulence. Cell 84, 165–174 (1996).

32. F. C. Soncini, E. Garcia Vescovi, F. Solomon, E. A. Groisman, Molecular basis of the magnesium deprivation response in Salmonella typhimurium: identification of PhoP-regulated genes. J Bacteriol 178, 5092–5099 (1996).

33. S. Eriksson, S. Lucchini, A. Thompson, M. Rhen, J. C. Hinton, Unravelling the biology of macrophage infection by gene expression profiling of intracellular Salmonella enterica. Mol Microbiol 47, 103–118 (2003).

34. V. I. Petrou et al., Structures of aminoarabinose transferase ArnT suggest a molecular basis for lipid A glycosylation. Science 351, 608–612 (2016).

35. A. Anandan et al., Structure of a lipid A phosphoethanolamine transferase suggests how conformational changes govern substrate binding. Proceedings of the National Academy of Sciences 114, 2218–2223 (2017).

36. R. E. Bishop, The lipid A palmitoyltransferase PagP: molecular mechanisms and role in bacterial pathogenesis. Molecular microbiology 57, 900–912 (2005).

37. E. Kintz et al., Salmonella enterica serovar Typhi lipopolysaccharide O-antigen modification impact on serum resistance and antibody recognition. Infection and immunity 85, 10.1128/iai.01021-01016 (2017).

38. J. V. Farizano, M. de las Mercedes Pescaretti, F. E. López, F.-F. Hsu, M. A. Delgado, The PmrAB system-inducing conditions control both lipid A remodeling and O-antigen length distribution, influencing the Salmonella Typhimurium-host interactions. Journal of Biological Chemistry 287, 38778–38789 (2012).

39. P. A. Fernández et al., Fnr and ArcA regulate lipid a hydroxylation in Salmonella enteritidis by controlling lpxO expression in response to oxygen availability. Frontiers in microbiology 9, 1220 (2018).

40. J. Jennewein et al., Low-oxygen tensions found in S almonella-infected gut tissue boost S almonella replication in macrophages by impairing antimicrobial activity and augmenting S almonella virulence. Cellular microbiology 17, 1833–1847 (2015).

41. D. A. Cano, M. G. Pucciarelli, M. Martínez-Moya, J. Casadesús, F. García-del Portillo, Selection of small-colony variants of Salmonella enterica serovar typhimurium in nonphagocytic eucaryotic cells. Infection and immunity 71, 3690–3698 (2003).

42. C. G. Moreira et al., Virulence and stress-related periplasmic protein (VisP) in bacterial/host associations. Proceedings of the National Academy of Sciences 110, 1470–1475 (2013).

43. W. T. Doerrler, H. S. Gibbons, C. R. Raetz, MsbA-dependent translocation of lipids across the inner membrane of Escherichia coli. J Biol Chem 279, 45102–45109 (2004).

44. H. Nikaido, Molecular basis of bacterial outer membrane permeability revisited. Microbiology and molecular biology reviews 67, 593–656 (2003).

45. P. I. Fields, R. V. Swanson, C. G. Haidaris, F. Heffron, Mutants of Salmonella typhimurium that cannot survive within the macrophage are avirulent. Proceedings of the National Academy of Sciences 83, 5189–5193 (1986).

46. M. Snavely, C. Miller, M. Maguire, The mgtB Mg2+ transport locus of Salmonella typhimurium encodes a P-type ATPase. Journal of Biological Chemistry 266, 815–823 (1991).

47. K. A. Datsenko, B. L. Wanner, One-step inactivation of chromosomal genes in Escherichia coli K-12 using PCR products. Proceedings of the National Academy of Sciences 97, 6640–6645 (2000).

48. E. J. Lee, E. A. Groisman, An antisense RNA that governs the expression kinetics of a multifunctional virulence gene. Molecular microbiology 76, 1020–1033 (2010).

49. B. R. Bochner, H.-C. Huang, G. L. Schieven, B. N. Ames, Positive selection for loss of tetracycline resistance. Journal of bacteriology 143, 926–933 (1980).

50. B.-K. Cho, E. M. Knight, B. Ø. Palsson, PCR-based tandem epitope tagging system for Escherichia coli genome engineering. Biotechniques 40, 67–72 (2006).

51. C. Y. Eugene, M. Hackett, Rapid isolation method for lipopolysaccharide and lipid A from gram-negative bacteria. Analyst 125, 651–656 (2000).

52. H. Lee, F.-F. Hsu, J. Turk, E. A. Groisman, The PmrA-regulated pmrC gene mediates phosphoethanolamine modification of lipid A and polymyxin resistance in Salmonella enterica. Journal of bacteriology 186, 4124–4133 (2004).

53. S. Y. Park, E. A. Groisman, Signal-specific temporal response by the S almonella PhoP/PhoQ regulatory system. Molecular microbiology 91, 135–144 (2014).

54. L. A. Lewis et al., Phosphoethanolamine substitution of lipid A and resistance of Neisseria gonorrhoeae to cationic antimicrobial peptides and complement-mediated killing by normal human serum. Infection and immunity 77, 1112–1120 (2009).

55. J. Abramson et al., Accurate structure prediction of biomolecular interactions with AlphaFold 3. Nature, 1–3 (2024).

56. L. Schrödinger, W. DeLano (2020) PyMOL.

## SI References

1. Uncategorized References

1. E. Alix, A. B. Blanc-Potard, Peptide-assisted degradation of the Salmonella MgtC virulence factor. The EMBO journal 27, 546–557 (2008).

2. S. Choi et al., The Salmonella virulence protein MgtC promotes phosphate uptake inside macrophages. Nature communications 10, 3326 (2019).

3. E. F. Pettersen et al., UCSF ChimeraX: Structure visualization for researchers, educators, and developers. Protein science 30, 70–82 (2021).

4. H. S. Gibbons, S. Lin, R. J. Cotter, C. R. Raetz, Oxygen requirement for the biosynthesis of theS-2-hydroxymyristate moiety in Salmonella typhimurium lipid A: FUNCTION OF LpxO, A new Fe2+/α-KETOGLUTARATE-DEPENDENT dioxygenase homologue. Journal of Biological Chemistry 275, 32940–32949 (2000).

5. H. Lee, F.-F. Hsu, J. Turk, E. A. Groisman, The PmrA-regulated pmrC gene mediates phosphoethanolamine modification of lipid A and polymyxin resistance in Salmonella enterica. Journal of bacteriology 186, 4124–4133 (2004).

6. P. I. Fields, R. V. Swanson, C. G. Haidaris, F. Heffron, Mutants of Salmonella typhimurium that cannot survive within the macrophage are avirulent. Proceedings of the National Academy of Sciences 83, 5189–5193 (1986).

7. E.-J. Lee, M. H. Pontes, E. A. Groisman, A bacterial virulence protein promotes pathogenicity by inhibiting the bacterium’s own F1Fo ATP synthase. Cell 154, 146–156 (2013).

8. P. I. Fields, E. A. Groisman, F. Heffron, A Salmonella locus that controls resistance to microbicidal proteins from phagocytic cells. Science 243, 1059–1062 (1989).

9. R. G. Taylor, D. C. Walker, R. McInnes, E. coli host strains significantly affect the quality of small scale plasmid DNA preparations used for sequencing. Nucleic acids research 21, 1677 (1993).

10. G. Karimova, J. Pidoux, A. Ullmann, D. Ladant, A bacterial two-hybrid system based on a reconstituted signal transduction pathway. Proceedings of the National Academy of Sciences 95, 5752–5756 (1998).

11. K. A. Datsenko, B. L. Wanner, One-step inactivation of chromosomal genes in Escherichia coli K-12 using PCR products. Proceedings of the National Academy of Sciences 97, 6640–6645 (2000).

12. B.-K. Cho, E. M. Knight, B. Ø. Palsson, PCR-based tandem epitope tagging system for Escherichia coli genome engineering. Biotechniques 40, 67–72 (2006).

13. F. C. Soncini, E. G. Véscovi, E. A. Groisman, Transcriptional autoregulation of the Salmonella typhimurium phoPQ operon. Journal of bacteriology 177, 4364–4371 (1995).

14. S. Chamnongpol, E. A. Groisman, Mg2+ homeostasis and avoidance of metal toxicity. Molecular microbiology 44, 561–571 (2002).

15. L.-M. Guzman, D. Belin, M. J. Carson, J. Beckwith, Tight regulation, modulation, and high-level expression by vectors containing the arabinose PBAD promoter. Journal of bacteriology 177, 4121–4130 (1995).

16. G. Karimova, A. Ullmann, D. Ladant, Protein-protein interaction between Bacillus stearothermophilus tyrosyl-tRNA synthetase subdomains revealed by a bacterial two-hybrid system. Journal of molecular microbiology and biotechnology 3, 73–82 (2001).

